# Combinatorial tumor suppressor inactivation efficiently initiates lung adenocarcinoma with therapeutic vulnerabilities

**DOI:** 10.1101/2021.10.20.464849

**Authors:** Maryam Yousefi, Gábor Boross, Carly Weiss, Christopher W. Murray, Jess D. Hebert, Hongchen Cai, Emily L. Ashkin, Saswati Karmakar, Laura Andrejka, Leo Chen, Minwei Wang, Min K. Tsai, Wen-Yang Lin, Chuan Li, Pegah Yakhchalian, Caterina I. Colón, Su- Kit Chew, Pauline Chu, Charles Swanton, Christian A. Kunder, Dmitri A. Petrov, Monte M. Winslow

**Affiliations:** Department of Genetics, Stanford University School of Medicine, Stanford, CA, USA; Cancer Biology Program, Stanford University School of Medicine, Stanford, CA, USA; Department of Pathology, Stanford University School of Medicine, Stanford, CA, USA; Department of Biology, Stanford University, Stanford, CA, USA; Cancer Evolution and Genome Instability Laboratory, University College London Cancer Institute, London, UK; Cancer Evolution and Genome Instability Laboratory, The Francis Crick Institute, London, UK; Department of Medicine, David Geffen School of Medicine at University of California, Los Angeles, Los Angeles, CA, USA

## Abstract

Lung cancer is the leading cause of cancer death worldwide, with lung adenocarcinoma being the most common subtype. Many oncogenes and tumor suppressor genes are altered in this cancer type and the discovery of oncogene mutations has led to the development of targeted therapies that have improved clinical outcomes. However, a large fraction of lung adenocarcinomas lacks mutations in known oncogenes, and the genesis and treatment of these oncogene-negative tumors remain enigmatic. Here, we perform iterative *in vivo* functional screens using quantitative autochthonous mouse model systems to uncover the genetic and biochemical changes that enable efficient lung tumor initiation in the absence of oncogene alterations. Through the generation of hundreds of diverse combinations of tumor suppressor alterations, we demonstrate that the inactivation of suppressors of the RAS and PI3K pathways drive the development of oncogene-negative lung adenocarcinoma. Human genomic data and histology identified RAS/MAPK and PI3K pathway activation as a common event in oncogene-negative human lung adenocarcinomas. We demonstrate that these Onc-negative^RAS/PI3K^ tumors and related cell lines are vulnerable to pharmacological inhibition of these signaling axes. These results transform our understanding of this prevalent yet understudied subtype of lung adenocarcinoma.

## INTRODUCTION

Lung cancer is the leading cause of cancer death^1^. Lung adenocarcinoma, the most prevalent subtype of lung cancer, has frequent alterations in receptor tyrosine kinase and RAS/RAF pathway oncogenes, including mutations in *EGFR* and *KRAS*^2^. The identification of driver oncogenes has enabled a shift from toxic chemotherapies to less toxic and more effective therapies that often target the oncogenes^3^. However, approximately 30 percent of lung adenocarcinomas are thought to lack a driving oncogene^4–6^. Consequently, developing targeted therapies for these tumors remains a major unmet challenge for precision thoracic oncology.

Extensive genomic and transcriptomic studies suggest that neither technical reasons nor the presence of novel oncogenes likely explain this large and clinically significant population of lung cancer patients^1, 2, 4–12^. Thus, despite the diagnosis of more than 150,000 patients per year with oncogene-negative lung adenocarcinomas worldwide, the genetic events and biochemical pathway changes that drive the initiation and growth of these tumors remain almost entirely unknown.

Oncogenes and tumor suppressor genes are parts of signaling networks that generate and sustain the biochemical changes that drive tumor initiation and growth^13–16^. Combinatorial alterations in tumor suppressor genes could co-operate to activate pathways driving oncogene-negative lung tumors. Human lung adenocarcinoma have complex patterns of mutations across many putative tumor suppressor genes^4^. However, the ability to predict which combinations of genomic alterations drive cancer in the absence of oncogene activation based on human genomic data alone remains challenging. While human genomic data can predict combinations of genomic mutations as likely cancer drivers when the mutations co-occur at very high frequencies ^17–20^, identifying pathogenic combinations of less frequently mutated genes poses a nearly insurmountable statistical challenge. Furthermore, the large numbers of mutations in lung cancers, non-genomic mechanism that often inactivate tumor suppressor genes, and generation of similar biochemical effects through inactivation of different genes further reduce the ability of human cancer genomic studies to identify combinatorial alterations that activate driver pathways in lung cancer ^21–24^.

Functional genomic studies within autochthonous cancer models can help identify the pathways involved in tumorigenesis *in vivo* ^25^. Here, we leveraged quantitative mouse model systems to assess the ability of hundreds of combinatorial alterations of tumor suppressor genes, acting across many different signaling pathways, to generate oncogene-negative lung adenocarcinomas *in vivo*. We uncover pathway-level changes that drive lung cancer in the absence of oncogene mutations, translate these findings to human oncogene-negative lung adenocarcinoma, and leveraged these results to identify therapeutic vulnerabilities.

## RESULTS

### A large fraction of human lung adenocarcinomas lack oncogene mutations

To better understand the genomics of lung adenocarcinomas that lack oncogene mutations, we analyzed data from The Cancer Genome Atlas (TCGA) and AACR Genomics Evidence Neoplasia Information Exchange (GENIE) ^26, 27^. We classified tumors as oncogene-positive if they had high-confidence oncogenic alterations in previously described proto-oncogenes, oncogene-indeterminate if they had alterations of unknown significance in known proto-oncogenes, and oncogene-negative if they had no alterations in known proto-oncogenes (**Methods**). Consistent with previous publications, we found that 17-18% of lung adenocarcinomas were oncogene-negative (**Figure 1a** and **S1a**) ^28–30^. Additionally, 15-27% of lung adenocarcinomas were oncogene-indeterminate and thus 32-45% of lung adenocarcinomas lack known oncogene mutations. Patients with oncogene-negative, oncogene-indeterminate, and oncogene-positive lung adenocarcinomas have broadly similar mutational burden and clinical characteristics (**Figure S1b-e**).

**Figure 1.**
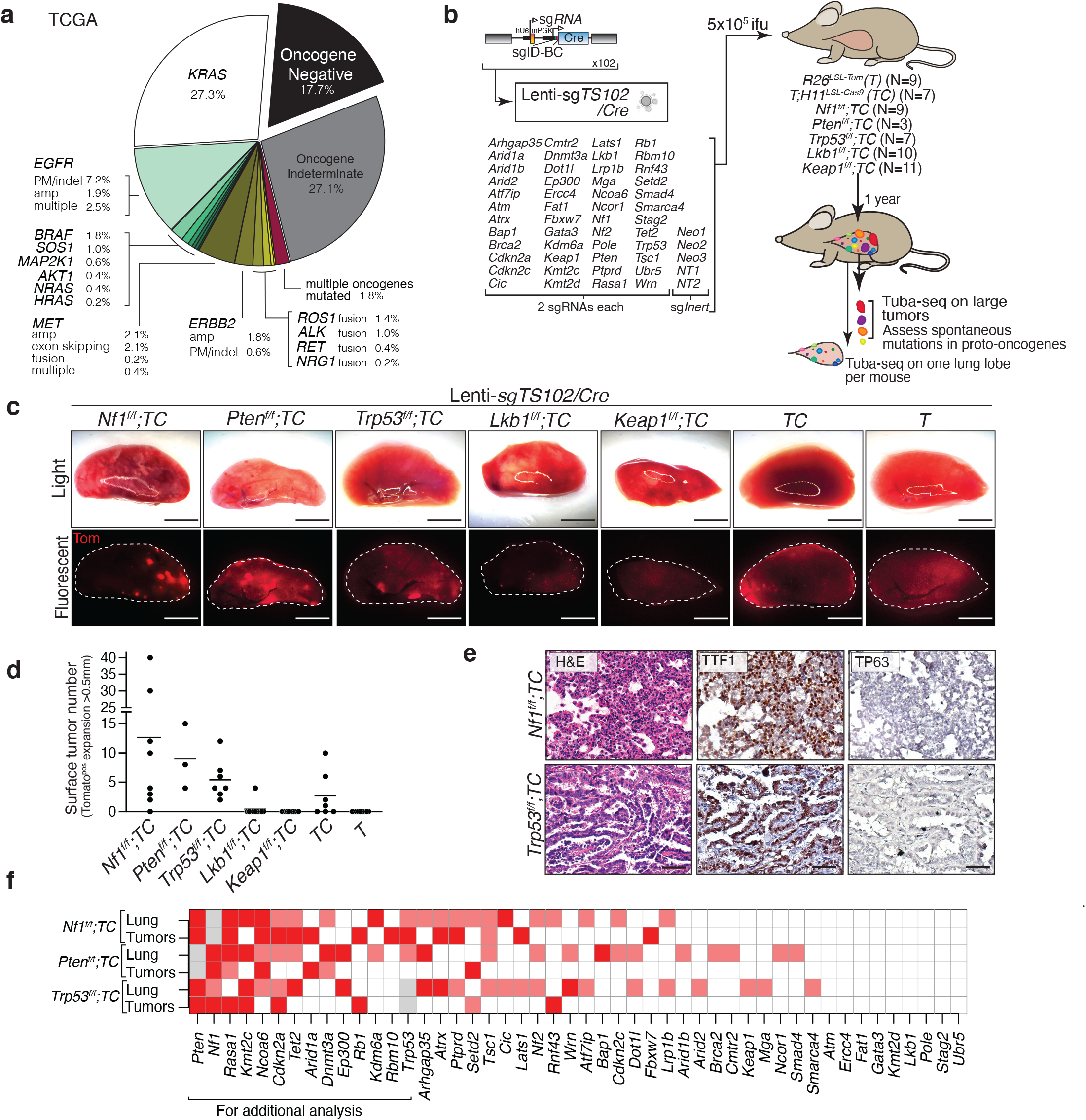
Combinatorial tumor suppressor inactivation enables lung tumor development in the absence of engineered oncogenes. **a.** Frequency of human lung adenocarcinomas with likely oncogenic alterations in proto-oncogenes (oncogene-positive), with alterations of unknown effects in proto-oncogenes (oncogene-indeterminate), and without any alterations in proto-oncogenes (oncogene-negative). Data from TCGA. PM: point mutation, indel: insertion and deletion, amp: amplification, multiple: multiple alterations in the same gene. **b.** Combined Cre/lox and CRISPR/Cas9-mediated tumor suppressor gene inactivation to generate lung epithelial cells with diverse genotypes. The number of mice in each group is indicated. **c.** Representative light and fluorescence images of lung lobes from the indicated genotypes of mice one year after transduction with the Lenti-sg*TS102/Cre* pool. Lung lobes are outlined with white dotted lines. Scale bar = 4 mm **d.** The number of surface tumors (defined as Tomato-positive expansions greater than 0.5 mm in diameter) quantified by direct counting. Each dot represents a mouse, and the bar is the mean. **e.** Representative Hematoxylin and Eosin (H&E), TTF1, and TP63 stained sections of the indicated genotypes of mice. Scale bar = 100 µm **f.** Heatmap showing two measures of tumor suppressor strengths in each genotype detected using Tuba-seq analysis: (1) in rows labeled as “Tumors” we assessed the occurrence of tumor suppressor gene targeting vectors in dissected tumors. p < 0.001 (red), p < 0.1 (pink) (see **Figure S4**) (2) In rows labeled as “Lung” we measured increases in median sizes of clonal expansions in the presence of indicated tumor suppressor alterations in bulk single lung lobe samples. Gene mutations showing significant increases (p<0.05) in sizes of clonal expansions using all sgRNAs are shown in red, and those with only one significant sgRNA are shown in pink (see **Figure S5**). Gray boxes indicate redundant targeting of genes by both Cre/*loxP* and CRISPR/Cas9.

### Combinatorial tumor suppressor gene inactivation enables lung tumor development

To determine whether combinatorial tumor suppressor gene inactivation can drive lung tumor initiation in the absence of oncogene activation, we coupled Cre/*loxP*-based genetically engineered mouse models and somatic CRISPR/Cas9-based genome editing with tumor barcoding and high-throughput barcode sequencing (Tuba-seq) ^31–35^. We used Cre*/loxP* to inactivate each of five “core” tumor suppressor genes (*Trp53*, *Lkb1/Stk11*, *Keap1*, *Nf1*, and *Pten)*. These genes are within diverse pathways and are frequently inactivated in human lung cancers, including oncogene-negative lung adenocarcinomas (**Figure S2a-b**) [35-38]. We used CRISPR/Cas9 to coincidentally inactivate panels of additional tumor suppressor genes in lung epithelial cells in mice with floxed alleles of each of the “core” tumor suppressors, a Cre-reporter allele (*R26^LSL-Tom^* (*T*) ^36^), and a Cre-regulated *Cas9* allele (*H11^LSL-Cas^*^9^ (*C*) ^37^).

We transduced *Nf1^f/f^;TC, Pten^f/f^;TC, Trp53^f/f^;TC, Lkb1^f/f^;TC, Keap1^f/f^;TC, TC*, and *T* mice with two pools of barcoded Lenti-sgRNA*/Cre* vectors that target ∼50 putative tumor suppressor genes that we previously investigated in KRAS^G12D^-driven lung tumors (Lenti-sg*TS*15*/Cre* and Lenti-sg*TS*102*/Cre*) (**Figure 1b, S2c-d**, **S3a**, and **Table S1**) ^31, 32, 35^. The mutation frequency of these genes varied, and mutations in some were enriched in oncogene-negative human lung adenocarcinomas (**Table S1)** (**Figure S2c-d**). The combination of Cre*/LoxP* and CRISPR/Cas9- based genome editing should generate hundreds of combinations of genomic alterations in lung epithelial cells. We previously found that a small percent of lung tumors initiated with Lenti-sgRNA*/Cre* vectors in other lung cancer models contained multiple sgRNAs, consistent with the transduction of the initial cell with multiple Lenti-sgRNA*/Cre* vectors ^31, 32^. Thus, we used a high titer of the Lenti-sgRNA/*Cre* pools in these experiments to increases the likelihood of finding higher-order genetic interactions that drive tumorigenesis.

One year after transduction with the Lenti-sg*RNA/Cre* pools, *Nf1^f/f^;TC*, *Pten^f/f^;TC,* and *Trp53^f/f^;TC* mice developed a modest number of tumors (defined as Tomato^positive^ expansion >0.5 mm in diameter), while *Lkb1^f/f^;TC* and *Keap1^f/f^;TC* mice rarely developed any tumors (**Figure 1c-d, S3b-c)**. Interestingly, *Nf1^f/f^;TC*, *Pten^f/f^;TC,* and *Trp53^f/f^;TC*, and *TC* mice transduced with the larger Lenti-sg*TS102/Cre* pool developed many more tumors than those transduced with the Lenti-sg*TS15/Cre* pool. These tumors were positive for TTF1/NKX2-1, a marker for lung adenocarcinoma, and negative for P63 and UCHL1, markers for squamous cell and small cell lung cancer, respectively (**Figure 1e**).

To determine whether these tumors contained spontaneous oncogene mutations, we PCR-amplified and sequenced 10 genomic regions in *Kras, Braf*, *Nras,* and *Egfr* (**Figure S3d**, **Table S2,** and **Methods**) ^33, 38–46^. Across 29 samples, we detected only one oncogene mutation (a *Kras^G12V^* mutation in a tumor from a *Pten^f/f^;TC* mouse). Thus, the majority of these tumors arose in the absence of hotspot mutations in these proto-oncogenes. This is consistent with the low mutation rate in mouse models of lung cancer ^47^ and suggests that the inactivation of combinations of specific tumor suppressor genes in *Nf1^f/f^;TC*, *Pten^f/f^;TC,* and *Trp53^f/f^;TC* mice drives the development of lung cancer *in vivo.* Notably, the overall low number of tumors indicates that inactivation of the “core” tumor suppressor genes alone, and most combinations of tumor suppressor genes tested, are insufficient to generate lung tumors.

### Identification of top candidate tumor suppressor genes involved in oncogene-negative lung tumor formation

The Lenti-sgRNA/Cre vectors contain two-component barcodes in which an sgID identifies the sgRNA and a random barcode (BC) uniquely tags each clonal tumor. Thus, high throughput sequencing of the sgID-BC region can identify the sgRNA(s) present in each tumor and quantify the number of cancer cells in each tumor (**Figure 1b**). To determine which sgRNAs were present in the largest tumors, we PCR-amplified the sgID-BC region from genomic DNA from dissected tumors and performed high-throughput sgID-BC sequencing.

Most large tumors contained multiple Lenti-sgRNA*/Cre* vectors therefore, we calculated the statistical enrichment of each sgRNA based on their relative representation in the dissected tumors (**Figure 1f** and **S4**, see **Methods**).

To further quantify the impact of inactivating each tumor suppressor gene on clonal expansion of lung epithelial cells, we performed tumor barcode sequencing (Tuba-seq) on bulk DNA from one lung lobe from each *Nf1^f/f^;TC*, *Pten^f/f^;TC, Trp53^f/f^;TC,* and *TC* mouse (**Figure 1c**). Analysis of the number of cells in clonal expansions further nominated tumor suppressor genes that may contribute to tumor initiation and growth (**Figure 1f**, and **S5**). Based on these two analyses, we selected 13 genes for further analysis (**Figure 1f**). The potential importance of these tumor suppressor genes was often supported by both sgRNAs targeting each gene, consistent with on-target effects. Finally, Lenti-sg*Pten/Cre* enrichment in tumors in *Nf1^f/f^;TC* mice and Lenti-sg*Nf1/Cre* enrichment in tumors in *Pten^f/f^;TC* mice cross-validate our screen (**Figure 1f** and **S4-5**).

### Inactivation of candidate tumor suppressors efficiently generates lung tumors

To determine the potential of the top candidate tumor suppressor genes to initiate oncogene-negative tumors, we generated a pool of Lenti-sgRNA*/Cre* vectors targeting each of these tumor suppressor genes and one vector with an inert sgRNA (Lenti-sg*TS14/Cre* pool; **Figure 2a**). We targeted each gene with the sgRNA that had the most significant effect on tumor growth and used five times higher titer of each lentiviral vector per mouse than we used in Lenti- sg*TS102/Cre* pool, thus increasing the potential for the transduction of the initial cell with multiple Lenti-sgRNA*/Cre* vectors.

**Figure 2.**
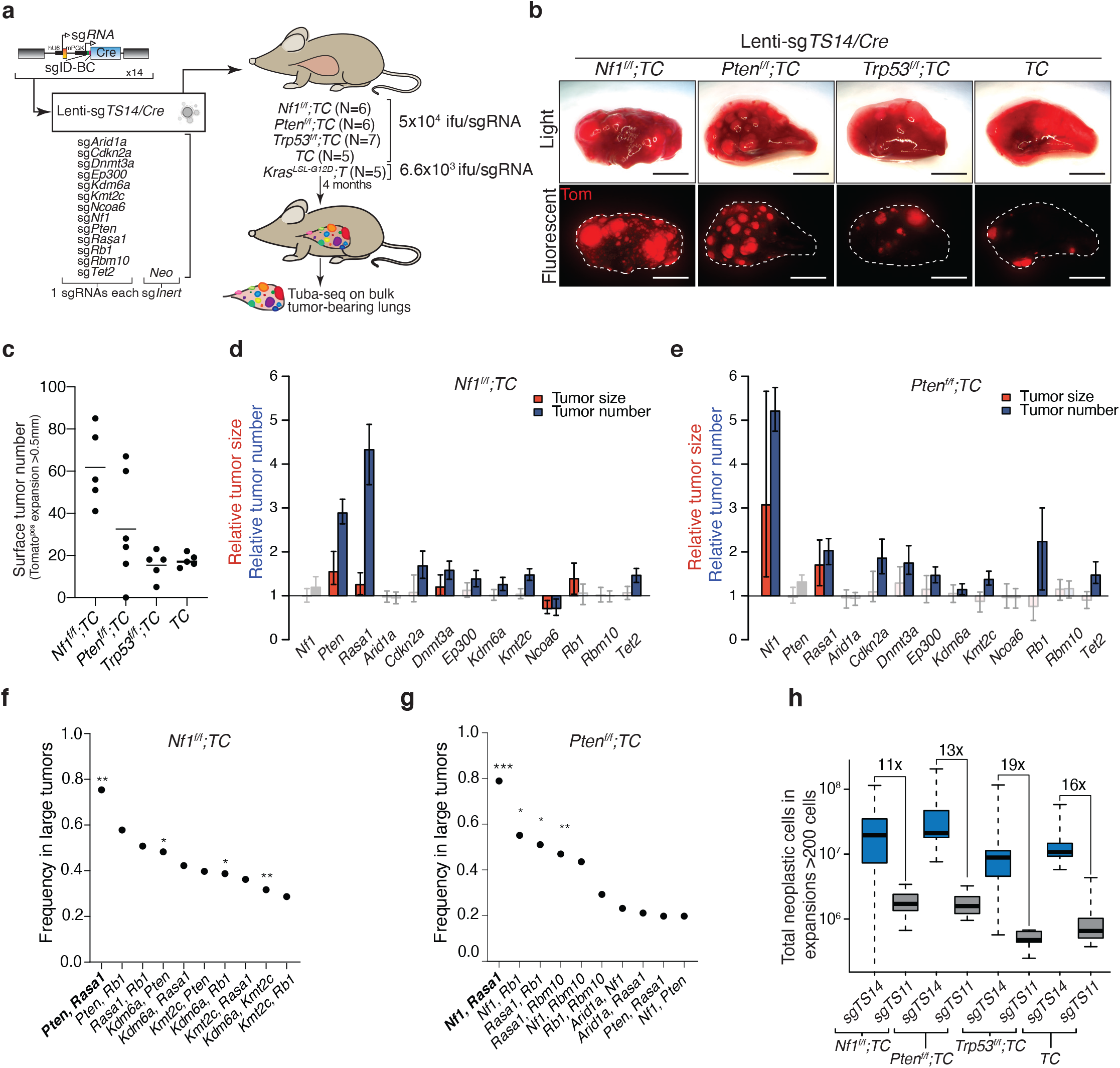
*Nf1*, *Rasa1*, and *Pten* emerge as key drivers of oncogene-negative lung adenocarcinoma. **a.** Combined Cre/lox and CRISPR/Cas9-mediated tumor suppressor gene inactivation to generate lung epithelial cells with diverse genotypes. The number of mice in each group is indicated. **b.** Representative light and fluorescence images of lung lobes from the indicated genotypes of mice. Lung lobes are outlined with white dotted lines. Scale bar = 4mm **c.** The number of tumors (defined as Tomato-positive expnsions larger than 0.5 mm in diameter) quantified by direct counting. Each dot represents a mouse, and the bar is the mean. **d,e.** The number of tumors with a minimum size of 1000 neoplastic cells relative to the inert sgRNA containing expansions is shown as blue bars. 90^th^ percentile of tumor sizes relative to the inert sgRNAs is shown as a red bar. sgRNAs resulting in significantly different tumor number or size (p<0.05) are shown in darker colors. Whiskers show 95% confidence intervals. Mouse genotypes are indicated. **f,g.** Barcodes with the highest counts in each mouse were investigated for coinfection with multiple Lenti-sg*TS/Cre* vectors(i.e., tumors initiated from cells transduced with multiple viruses, which result in complex tumor suppressor genotypes, see **Methods**). The top 10 pairs of tumor suppressors that were most frequently co-mutated are shown. Combinations of sgRNAs that lead to the generation of *Nf1*, *Rasa1*, and *Pten* mutant cancer cells are in bold. *p<0.05, **p<0.01, ***p<0.001 based on a permutation test. **h.** Total number of neoplastic cells in clonal expansions with more than 200 cells in the indicated genotypes of mice after receiving Lenti-*sgTS14/Cre* or Lenti-*sgTS11/Cre,* which lacks lentiviral vectors containing sg*Nf1*, sg*Rasa1*, and sg*Pten.* The magnitude of neoplastic cell number reduction in each group is indicated.

We initiated tumors with Lenti-sg*TS14*/*Cre* in *Nf1^f/f^;TC*, *Pten^f/f^;TC, Trp53^f/f^;TC, TC,* and *Kras^LSL-G12D^;T* (*KT*) mice. Less than four months after tumor initiation, several *Nf1^f/f^;TC* and *Pten^f/f^;TC* mice showed signs of extensive tumor burden. These mice developed many more tumors than mice of the same genotypes one year after transduction with the Lenti-sg*TS102/Cre* (compare **Figure 2b-c** with **Figure 1c-d**, and **S10c**). Thus, this pool of candidate tumor suppressor genes is enriched for those that generate oncogene-negative lung tumors.

We performed Tuba-seq on DNA from bulk tumor-bearing lungs to determine the number and size of tumors with each barcoded Lenti-sgRNA/*Cre* vector. Inactivation of *Nf1, Rasa1*, and *Pten* most dramatically increased tumor size and/or tumor number across all mouse genotypes (**Figure 2d-e**, **S6a-b**, and **Methods**). Inactivation of some of the other tumor suppressor genes less dramatically but significantly increased tumor size and/or tumor number in a genotype-specific manner. This suggests that additional molecular pathways altered by these tumor suppressor genes may also lead to early epithelial expansions.

The largest tumors in *Nf1^f/f^;TC*, *Pten^f/f^;TC, Trp53^f/f^;TC,* and *TC* mice were frequently generated through the inactivation of multiple tumor suppressor genes. Vectors targeting *Nf1*, *Rasa1*, and/or *Pten* were often present in the largest tumors, and the coincident targeting of *Nf1*, *Rasa1*, and *Pten* was the most frequent combination (**Figure 2f-g**, **S6c-h**). To gain greater insight into the contribution of *Nf1*, *Rasa1,* and *Pten* inactivation to the generation of oncogene-negative tumors, we transduced *Nf1^f/f^;TC*, *Pten^f/f^;TC,Trp53^f/f^;TC, TC,* and *KT* mice with a pool of Lenti-sgRNA*/Cre* vectors that lacked the vectors targeting *Nf1*, *Rasa1*, and *Pten* (*Lenti-sgTS11/Cre)* (**Figure S7a**). Approximately four months after transduction, these mice had many fewer tumors than mice transduced with Lenti-*sgTS14/Cre* pool (**Figure S7b-c** and **S10c**). Tuba-seq analysis confirmed a dramatic decrease in tumor burden relative to mice that received the Lenti- sg*TS14/Cre pool* (**Figure 2h**). Thus, the inactivation of *Nf1, Rasa1*, and *Pten* emerged as the most important contributors to the generation of oncogene-negative lung tumors.

Extensive experiments generating single and pairwise inactivation of tumor suppressor genes in individual mice led to the development of very few tumors even after long periods of time (**Figure S8-S9**). Thus, single and pairwise tumor suppressor gene inactivation is rarely sufficient to generate lung tumors and combinatorial inactivation of three or more tumor suppressor genes increases the efficiency of tumor development and/or accelerates the growth of oncogene-negative lung tumors.

### Combinatorial inactivation of Nf1, Rasa1, and Pten drives lung adenocarcinoma development comparably to oncogenic Kras mutation

To dissect the higher-order genetic interactions between *Nf1*, *Rasa1*, and *Pten*, we transduced *TC* and *Trp53^f/f^;TC* mice with a pool of eight lentiviral vectors that would inactivate *Nf1, Rasa1,* and *Pten* individually, in pairwise combinations, and all three simultaneously (Lenti- sg*TS^Triple-pool^/Cre*, **Figure 3a**). Three months after tumor initiation, *TC* mice had hundreds of large adenomas and adenocarcinomas (**Figure 3b-c** and **Figure S10a-e**). Tuba-seq analysis showed that most of the tumor burden arose as a consequence of concomitant inactivation of all three tumor suppressors, with single and pairwise inactivation of these genes generating very few tumors consistent with our previous observations (**Figure 3e** and **S8**). Additional inactivation of *Trp53* in *Trp53^f/f^;TC* mice did not increase tumor initiation suggesting that *Trp53* is not a major suppressor of oncogene-negative lung adenocarcinoma development at these early stages (**Figure 3b-f** and **Figure S10a-h**). Finally, to compare the tumor initiation potential of combinatorial *Nf1*, *Rasa1*, and *Pten* inactivation with that of a known oncogene, we transduced *Kras^LSL-G12D^;T* mice (which lack Cas9) with Lenti-sg*TS^Triple-pool^/Cre* (**Figure 3a**). Strikingly, coincident inactivation of *Nf1, Rasa1,* and *Pten* in *TC* mice was nearly as potent as oncogenic KRAS^G12D^ in driving lung tumor initiation *in vivo* (**Figure 3g** and **Methods**).

**Figure 3.**
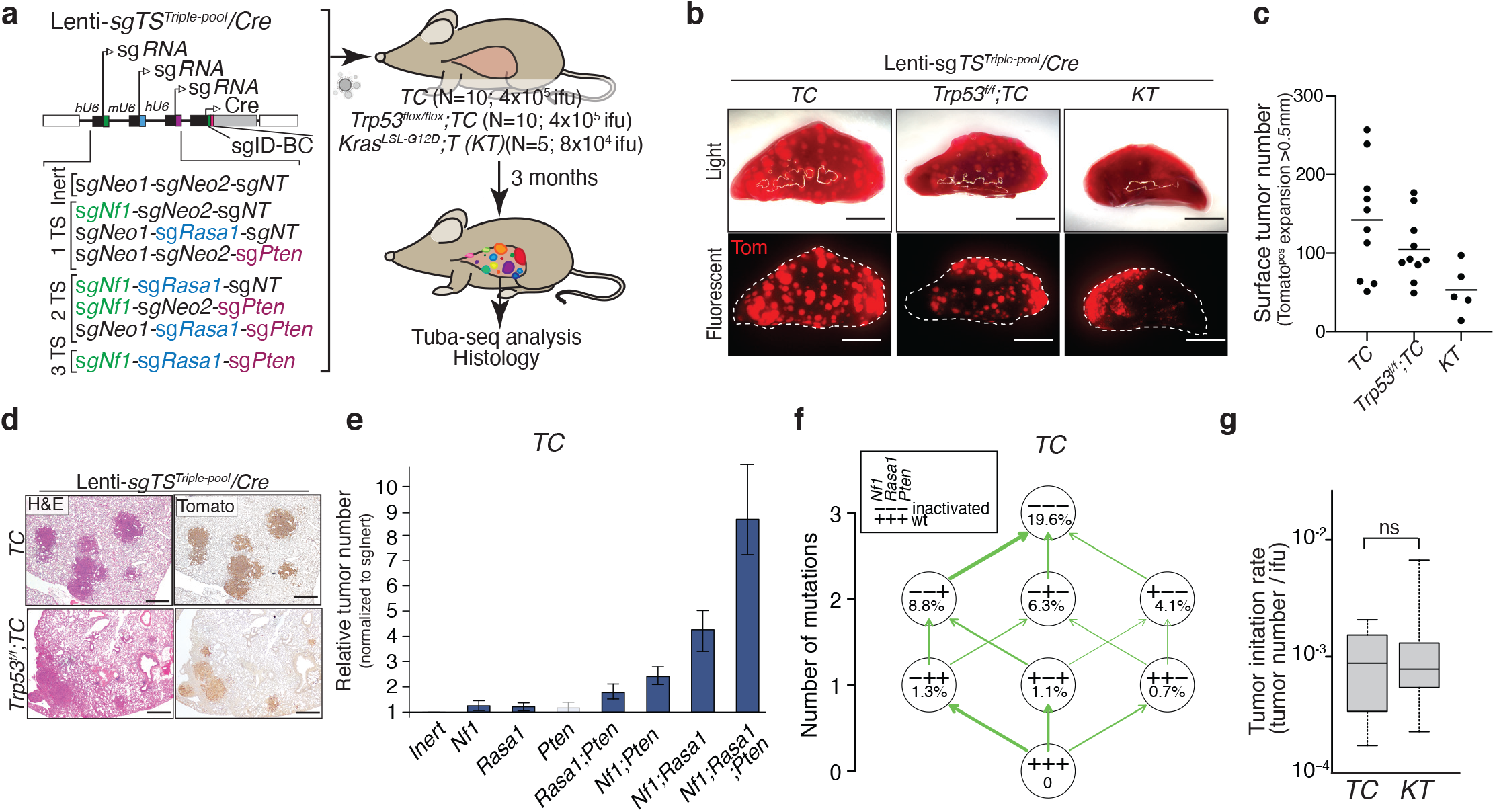
Inactivation of *Nf1*, *Rasa1*, and *Pten* allows a stepwise acquisition of growth advantage during lung adenocarcinoma development. **a.** 8 barcoded triple sgRNA vectors for CRISPR/Cas9-mediated inactivation of all combinations of *Nf1*, *Rasa1*, and *Pten* in *TC* and *Trp53^flox/flox^;TC* mice to assess genetic interactions between these tumor suppressors. sg*Neo1* and sg*Neo2* are active cutting, but inert sgRNAs that target *Neo^R^* in the *R26^LSL-tdTomato^* allele. *sgNT* is a non-targeting inert sgRNA. Mouse genotype, mouse number, and titer of virus delivered to each mouse are indicated. Tuba-seq was performed on tumor-bearing lungs 3 months after tumor initiation. **b.** Bright-field and fluorescence images of lungs from the indicated mouse genotypes. Lung lobes are outlined with a dashed white line. Scale bar = 4 mm **c.** The number of surface tumors (defined as Tomato-positive expansions larger than 0.5 mm in diameter) quantified by direct counting. Each dot represents a mouse, and the bar is the mean. **d.** Representative H&E and Tomato stained sections of lungs from *TC* and *Trp53^flox/flox^;TC* mice 3 months after transduction with Lenti-sg*TS^Triple-pool^/Cre.* Scale bar = 500 mm **e.** Numbers of tumors (with >1000 neoplastic cells) relative to the Inert sgRNA containing expansions. sgRNAs resulting in a significantly higher number of tumors than the inert vector (p<0.05) are shown in a darker color. Mean +/- 95% confidence interval is shown. **f.** Adaptive landscape of *Nf1*, *Rasa1*, and *Pten* inactivation in *TC* mice is shown. Nodes represent genotypes shown as a string of +(wild-type) and - (inactivat- ed) symbols representing *Nf1*, *Rasa1*, and *Pten*. Numbers in the nodes indicate fitness increase compared to wild-type. The relative probability of each beneficial mutation is shown as arrow widths (see **Methods**). **g.** Quantification of the ability of combined *Nf1/Rasa1/Pten* inactivation in *TC* mice and oncogenic *Kras^G12D^* in *KT* mice to initiate tumors. The number of tumors (with >1000 neoplastic cells) per infectious unit (ifu) is shown. The bar is the median, the box represents the interquartile range, and the whiskers show minimum and maximum values. ns: non-significant

In molecular evolution studies, generating combinations of genomic alterations and measuring the fitness of each genotype (growth rate) is used to infer the possible and the most probable paths from a wild-type state to a complex genotype ^48^. Through the generation of all possible combinatorial alterations of *Nf1*, *Rasa1*, and *Pten*, we quantified the fitness conferred by each mutation and the relative probabilities of different adaptive paths leading to the triple mutant genotype. Our data suggest that inactivation of these three genes can occur in any order, with each additional alteration further increasing the fitness (**Figure 3f**). The *Nf1 Rasa1 Pten* mutation sequence emerged as the most probable of all six possible paths.

To further analyze tumors driven by inactivation of *Nf1, Rasa1,* and *Pten*, we initiated tumors in *TC* and *Trp53^f/f^;TC* mice using only the lentiviral vector that targets all three genes (Lenti-sg*Nf1*-sg*Rasa1*-sg*Pten*/*Cre*) (**Figure S11a**). After only three months, these mice developed very large numbers of lung adenomas and adenocarcinomas (**Figure S11b-e**). We confirmed the inactivation of *Nf1*, *Rasa1*, and *Pten* in these tumors and whole-exome sequencing uncovered no putative oncogene mutations and only a few putative tumor suppressor mutations, none of which occurred in more than one tumor (**Figure S11f** and **Table S3**). Interestingly, at later timepoints after initiation, tumors in *Trp53^f/f^;TC* mice progressed to an invasive NKX2-1^negative^ HMGA2^positive^ state and metastasized to other organs such as liver similar to what has been reported in *Kras^G12D^;Trp53* mutant lung adenocarcinoma models (**Figure S12**) ^49^.

### Oncogene-negative murine lung adenocarcinomas have activated RAS and PI3K pathways

NF1 and RASA1 are negative regulators of RAS, while PTEN is a negative regulator of the PI3K-AKT pathway. Therefore, we investigated the impact of inactivating these tumor suppressor genes on RAS and PI3K pathway activation by immunohistochemistry, as well as by RNA-sequencing (RNA-seq) on FACS-isolated Tomato^positive^ cancer cells. We generated autochthonous tumors in which *Nf1*, *Rasa1*, and *Pten* were inactivated (*TC* mice with Lenti- sg*Nf1-*sg*Rasa1-*sg*Pten*/*Cre*; Nf1/Rasa1/Pten tumors), KRAS^G12D^ was expressed (*KT;H11^LSL-Cas^*^9^ mice with Lenti-sg*Inert*/*Cre*; Kras tumors), or KRAS^G12D^ was expressed and *Pten* was inactivated (*KT;H11^LSL-Cas^*^9^ mice with Lenti*-*sg*Pten*/*Cre*; Kras/Pten tumors) (**Figure S13a**).

Nf1/Rasa1/Pten tumors had positive staining for pERK (indicative of RAS pathway activation) and pAKT (indicative of PI3K pathway activation) (**Figure 4a**). Compared with Kras/Pten tumors, the average pERK staining in Nf1/Rasa1/Pten tumors was less intense and pAKT staining was similar (**Figure 4b-c**). Single-sample gene set variation analysis (ssGSVA) for previously reported gene sets representing RAS and PI3K-AKT regulated genes ^50, 51^ on our RNA-seq data confirmed that Nf1/Rasa1/Pten tumors had lower RAS pathway gene signature scores than Kras/Pten tumors **(Figure S13b)**. PI3K-AKT pathway gene signature scores were similar in Nf1/Rasa1/Pten and Kras tumors **(Figure S13c).** The rare tumors that eventually developed after pairwise inactivation of *Nf1*, *Rasa1*, and *Pten* also had strong activation of RAS and PI3K pathways (**Figure S8 and S13d**). Based on these analyses, we propose that these tumors represent a subtype of oncogene-negative lung adenocarcinomas with activated RAS and PI3K pathways (Onc-negative^RAS/PI3K^ subtype).

**Figure 4.**
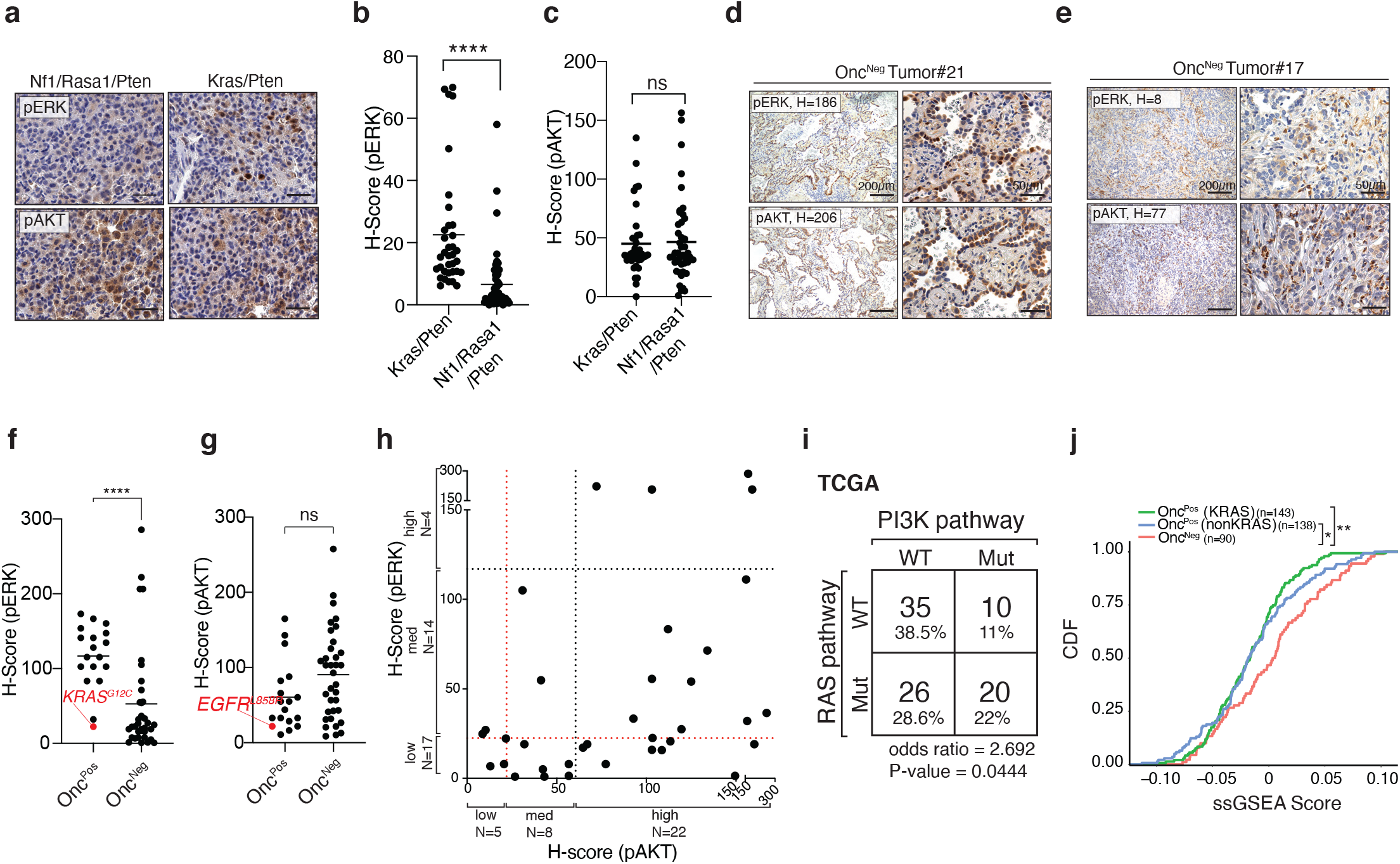
Oncogene-negative mouse and human lung adenocarcinomas have frequent activation of RAS and PI3K pathways. **a-c.**Representative immunohistochemistry for pERK and pAKT to determine activation of RAS and PI3K pathway in tumors with the indicated genotypes and quantification of these stainings. The bar is the mean. n.s: non-significant, ****<p<0.0001 using Mann–Whitney U test. Scale bars= 50µm **d,e.** Representative p-AKT and p-ERK-stained sections of oncogene-negative human tumors. H-scores for the whole section are indicated for each representative image. Scale bar= 200 µM (right), 50 µm (left) **f, g.** Quantification of pAKT and pERK staining on 35 oncogene-negative and 18 oncogene-positive human lung adenocarcinomas. Genotypes of oncogene- positive tumors with the lowest pERK and pAKT staining intensities are highlighted in red. Significance between groups was determined using Mann-Whitney U test, ns: non-significant, ****p<0.0001 **h.** pERK and pAKT H-scores for oncogene-negative human tumors are replotted from Figure 4f**,g**. Red dotted lines: the thresholds for low versus medium pERK and pAKT stains based on the lowest pERK staining intensity of oncogene-positive lung adenocarcinomas and the lowest pAKT staining level of *EGFR* mutant lung adenocarcinomas. Black dotted lines: the thresholds for medium versus high pERK and pAKT staining based on the mean pERK and pAKT H-scores in oncogene-positive tumors. The number of tumors in each staining intensity group (low, medium, high) is indicated on each axis of the plot. **i.** Alteration frequency of well-established components of RAS and PI3K pathways (see **Table S6**) and assessment of their co-occurrences based on TCGA data sets. p-value calculated by two-sided Fisher’s Exact Test. **j.** Cumulative distribution function (CDF) plot of the signature scores for human tumors stratified by genes upregulated in mouse oncogene-negative tumors generated by inactivation of *Nf1*, *Rasa1,* and *Pten* (**Figure S14a** and see **Table S4**). The cohort size and the P-value calculated by Kolmogorov–Smirnov test are indicated on the plot.

### Oncogene-negative human lung adenocarcinomas frequently have activation of RAS and PI3K pathways

To investigate the activation of RAS and PI3K pathways in human oncogene-negative lung adenocarcinomas, we analyzed oncogene-negative (N=35) and oncogene-positive (N=18) lung adenocarcinomas. Immunohistochemistry for pERK and pAKT showed that ∼45% of oncogene-negative human tumors had moderate to strong activation of both RAS and PI3K pathways and thus represent the Onco-negative^RAS/PI3K^ subtype (**Figure 4d-h, S13e-j)**. These tumors were genomically characterized by Stanford’s Solid Tumor Actionable Mutation Panel (STAMP)^52^. Activation of the RAS and PI3K pathways were rarely explained by mutations in *NF1*, *PTEN*, or other genes profiled by STAMP (**Table S5 and S6)**, likely due to the noncomprehensive tumor suppressor gene panel characterized by STAMP, as well as epigenetic mechanisms of RAS and PI3K pathway activation. Epigenetic silencing and other non-genomic mechanisms have been well documented to inhibit tumor suppressor genes including *PTEN* ^22, 23, 53, 54^ . Therefore, we performed immunohistochemistry for PTEN on 20 oncogene-negative lung adenocarcinomas that did not have genomic *PTEN* mutations. Consistent with previous reports, we observed low PTEN protein levels in 13 out of 20 of these tumors (**Figure S14a-f**) ^22^.

To assess a larger set of oncogene-negative lung adenocarcinomas for alterations that could lead to the activation of RAS and PI3K pathways, we analyzed oncogene-negative tumors in TCGA and GENIE datasets. We queried a set of well-established negative regulators of the RAS and PI3K pathways for alterations in oncogene-negative tumors (**Table S6**). Consistent with previous reports, *NF1* and *RASA1* alterations were enriched in oncogene-negative tumors; however, coincident genomic alterations in *NF1*, *RASA1*, and *PTEN* were rare (**Figure S14g-h**) ^55, 56^. However, over 60% of oncogene-negative lung adenocarcinomas in TCGA had alterations in either the RAS or PI3K pathways, and 22% of these tumors had alterations in components of both pathways, likely representing oncogene-negative^RAS/PI3K^ lung adenocarcinomas (**Figure 4i)**. These frequencies were lower in the GENIE dataset, possibly because only a fraction of the known genes in these pathways were analyzed (**Figure S14i)**. These histological and genomic analyses support a model in which activation of the RAS and PI3K pathways in Onc- negative^RAS/PI3K^ tumors can be generated by diverse genomic and/or epigenetic alterations.

Finally, we assessed whether Onc-negative^RAS/PI3K^ tumors in our mouse model more broadly exhibit transcriptional features that are consistent oncogene-negative human lung adenocarcinoma. We generated a gene expression signature of murine Onc-negative^RAS/PI3K^ tumors comprised of genes that are higher in Nf1/Rasa1/Pten tumors relative to Kras tumors in mice. We then calculated gene signature activity scores for each TCGA lung adenocarcinoma for this Onc-negative^RAS/PI3K^ gene expression signature using single-sample GSEA **(Table S4**).

Interestingly, the Onc-negative^RAS/PI3K^ signature was highest in oncogene-negative human lung adenocarcinomas relative to lung adenocarcinomas driven by oncogenic *KRAS* or other known oncogenes (**Figure 4j**). Collectively, these data indicate that the molecular and biochemical state of mouse Onc-negative^RAS/PI3K^ tumors recapitulates that of a substantial fraction of oncogene-negative human lung adenocarcinomas.

### Onc-negative^RAS/PI3K^ tumors are vulnerable to inhibition of RAS and PI3K-AKT pathways

Understanding the biochemical changes that drive tumor development can nominate potential therapeutic strategies ^38^. To investigate the therapeutic benefit of targeting key nodes in Onc-negative^RAS/PI3K^ lung cancer, we initiated tumors in *TC* mice with a smaller pool of barcoded sgRNA viral vectors targeting *Nf1*, *Rasa1*, and *Pten*. We treated these mice with the SHP2 inhibitor RMC-4550 ^57^, AKT1/2 inhibitor capivasertib ^58, 59^, or a combination of the two (**Figure 5a** and **S15a-b**). These drugs were chosen based on their ongoing clinical development and ability to reduce activation RAS and PI3K pathways ^57, 59^.

**Figure 5.**
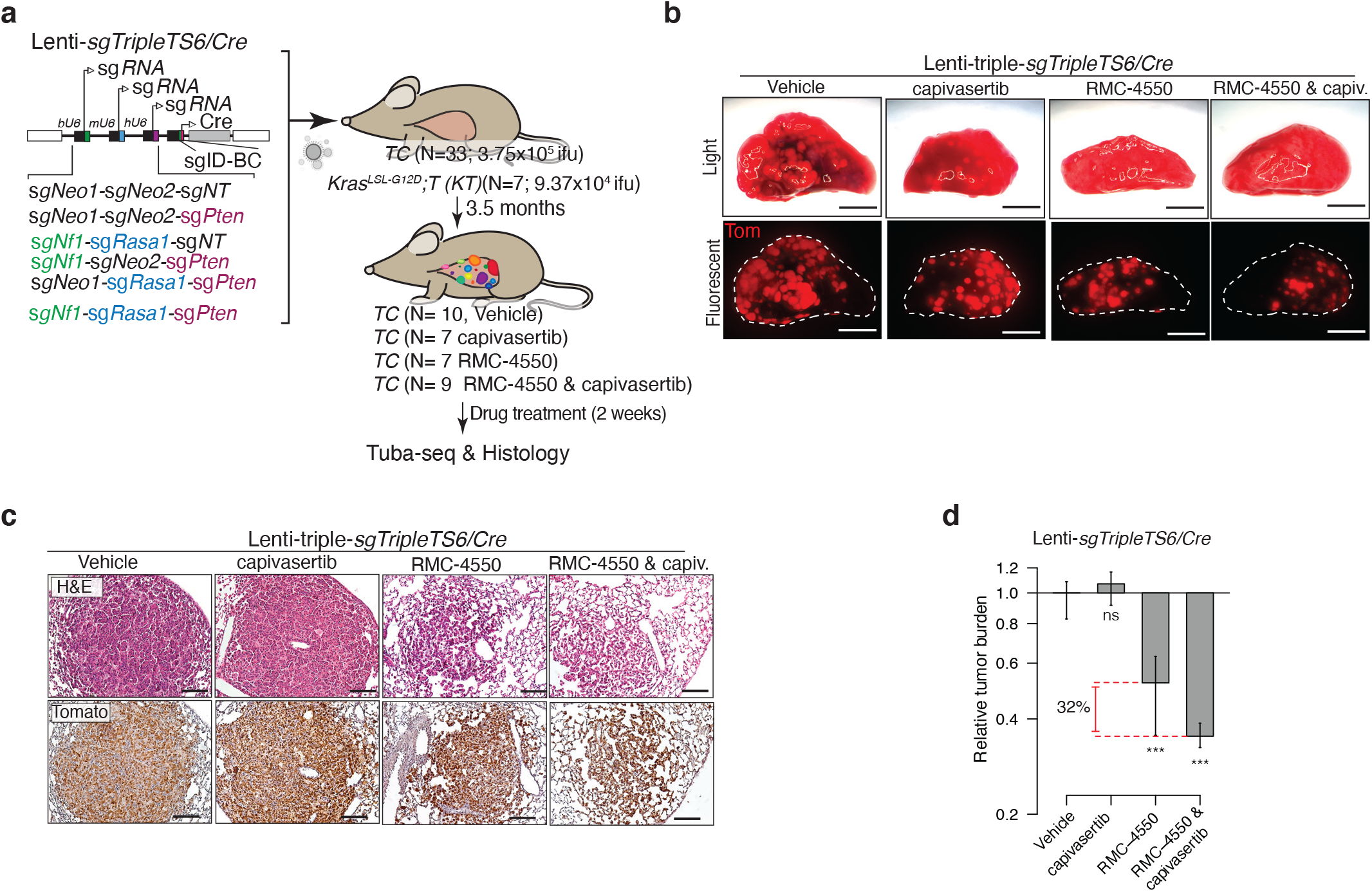
SHP2 and AKT inhibition synergize to reduce the growth of autochthonous oncogene-negative lung tumors. **a.** Barcoded triple sgRNA vectors for CRISPR/Cas9-mediated inactivation of combinations of *Nf1*, *Rasa1*, and *Pten* in *TC* mice to determine the response of oncogene-negative tumors to pharmacological inhibition of RAS and PI3K pathways. Indicated numbers of mice were treated with RMC-4550 (SHP2 inhibitor), capivasertib (AKT inhibitor), or combination of these two drugs for two weeks 3.5 months after tumor initiation. Tuba-seq and histological analysis were performed on tumor-bearing lungs followed by analysis of tumor response to therapies. **b.** Bright-field and fluorescence images of lungs from the indicated mice. Lung lobes are outlined with a dashed white line. Scale bars = 4 mm **c.** Representative H&E and Tomato-stained sections of tumors from *TC* mice 3.5 months after transduction with Lenti-sg*TripleTS6/Cre* and two weeks after treatment with the indicated drugs. Scale bars = 100 µm **d.** Relative tumor burden in mice after treatment with capivasertib, RMC-4550, and combination of these two drugs compared with tumor burden in vehicle-treated mice. ns: non-significant, ***p< 0.001. Drug response is shown for all the tumors.

Direct fluorescence imaging and histology indicated that SHP2 inhibition and combined SHP2 and AKT1/2 inhibition greatly reduced tumor burden (**Figure 5b-c** and **S15c**). Tuba-seq analysis provided greater insights into the overall and genotype-specific responses of tumors to the therapeutic interventions. Capivasertib monotherapy was ineffective *in vivo* while RMC- 4550 reduced the total tumor burden. The combination of RMC-4550 and capivasertib trended towards being the most efficient therapeutic approach reducing tumor burden by ∼30% compared with RMC-4550 alone (**Figure 5d, S15d-g**).

We confirmed the inhibition of RAS and PI3K pathways in oncogene-negative^RAS/PI3K^ tumors in mice treated with RMC-4550 and capivasertib by immunohistochemistry (**Figure S15h**). Furthermore, global gene expression analysis confirmed the downregulation of RAS and PI3K-AKT gene expression signatures after coincident SHP2 and AKT1/2 inhibition (**Figure S16a-d**). Treated tumors tended to have higher expression of an apoptosis gene expression signature and lower expression of a G2/M gene expression signature, suggesting that this combination treatment induces broad cellular changes in oncogene-negative tumors (**Figure S16e-f**).

### Inhibition of SHP2 and AKT synergizes to reduce the growth of Onc-negative^RAS/PI3K^ lung adenocarcinoma cell lines

To more extensively characterize the responses to SHP2 and AKT inhibition, we generated *Nf1/Rasa1/Pten* deficient Onc-negative^RAS/PI3K^ cell lines from tumors initiated with Lenti-sg*Nf1*-sg*Rasa1*-sg*Pten*/*Cre* in *Trp53^flox/flox^*;*TC* mice (**S17a-c**). As anticipated, RAS and PI3K signaling was reduced in response to treatment with RMC-4550 and capivasertib, respectively (**Figure S17d**). RMC-4550 and capivasertib each decreased the overall growth of three oncogene-negative^RAS/PI3K^ cell lines in a dose-dependent manner (**Figure 6a** and **S17e, g**). Consistent with our *in vivo* observations, RMC-4550 and capivasertib synergized to inhibit the growth of these cell lines (**Figure 6b,** and **S17f, h**). RAS and PI3K signaling can promote cell growth and survival [58, 59], and RMC-4550 and capivasertib inhibited proliferation and induced apoptosis to a greater extent than either RMC-4550 or capivasertib alone (**Figure 6c-d**).

**Figure 6.**
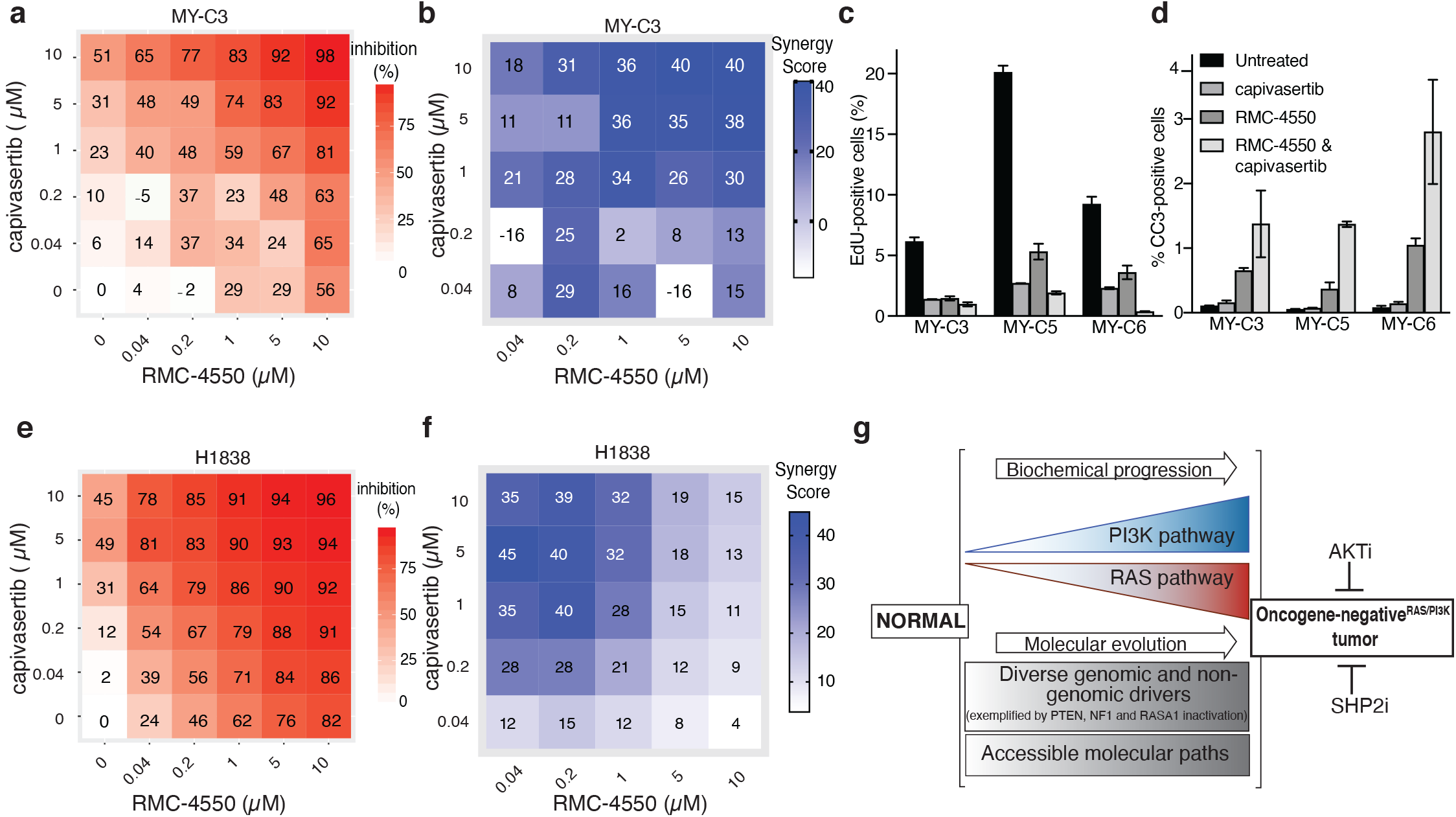
RMC-4550 and capivasertib synergize to inhibit the growth of Onc-negative^RAS/PI3K^ lung adenocarcinoma cell lines. **a.** Drug dose-response matrix depicting % growth inhibition of amurine Onc-negative^RAS/PI3K^ cell line after four days of treatment with the indicated doses of RMC-4550 and capivasertib. The average responses of three to four replicates are shown for each drug/drug combination. (see also **Figure S17 a-h)** **b.** Loewe’s synergy score calculated based on drug responses in Figure 6a. Synergy scores indicate the percentage of response beyond expectation. **c,d.** Cell proliferation and apoptosis analysis using EdU incorporation, cleaved caspase 3 staining, and flow-cytometry analysis. Three independent Onc-negative^RAS/PI3K^ murine cell lines were treated with 10 µM of the indicated drug/drugs for 2 days before the analysis. **e.** Drug dose-response matrix depicting % growth inhibition of H1838, a human oncogene-negative^RAS/PI3K^ lung adenocarcinoma cell line. **f.** Loewe’s synergy score calculated based on drug responses in Figure 6e. **g.** Model of biochemical progression and molecular drivers of Onc-negative^RAS/PI3K^ tumors.

Building on these findings, we assessed activation of RAS and PI3K pathways and driver pathway vulnerabilities in two oncogene-negative human lung adenocarcinoma cell lines, NCI- H1838 (*NF1^LOF^*) and NCI-H1623 (*RASA1^LOF^*). H1838 and H1623 had activation of RAS and PI3K pathways (**Figure S17i**). Consistent with our findings in mouse Onc-negative^RAS/PI3K^ cell lines, RMC-4550 synergizes with capivasertib to inhibit the growth of these human Onc- negative^RAS/PI3K^ lung adenocarcinoma cell lines (**Figure 6e-f and S17j-k**). These *in vivo* and cell culture analyses indicate that Onc-negative^RAS/PI3K^ tumors are vulnerable to therapeutic inhibition of these pathways.

## DISCUSSION

It is often overlooked that lung adenocarcinomas without genomic alterations in oncogenes afflict as many patients as those driven by either oncogenic KRAS or EGFR. To identify whether combinatorial inactivation of multiple tumor suppressor genes drives the initiation and growth of lung adenocarcinoma in the absence of oncogene activation, we performed a series of multiplexed *in vivo* functional genomic screens. By querying an extensive set of tumor suppressor gene alterations, we uncovered combinatorial tumor suppressor inactivation as a key driver of oncogene-negative lung adenocarcinomas. Importantly, combinatorial inactivation of negative regulators of RAS and PI3K pathways are as potent as oncogenic KRAS^G12D^ in initiating lung tumors *in vivo*.

Furthermore, while *NF1* inactivation is sometimes suggested to be an “oncogenic driver” in lung adenocarcinoma ^4, 29, 60^, *Nf1* inactivation alone is insufficient to initiate lung tumors (**Figure S8**). Even pairwise inactivation of *Nf1* and *Rasa1,* as well as many other tumor suppressor genes, generated very few tumors even after long time periods (**Figure S8**). These data suggest that genomic and/or epigenetic alterations in multiple genes within and across pathways may be required to surpass the thresholds necessary for Onc-negative^RAS/PI3K^ lung adenocarcinoma initiation and growth.

Although cancers harbor diverse genomic and epigenomic alterations, these alterations often converge on key pathways and generate similar biochemical changes ^15, 61^. For example, myeloid leukemia can be driven by gain-of-function mutations in *KRAS*, *NRAS*, or the receptor tyrosine kinase *FLT3*, or combined inactivation of multiple negative regulators of RAS pathway such as *SPRY4* and *NF1* ^62, 63^. Pathway activation through genomic and epigenomic inactivation of tumor suppressors can be very diverse, precluding the identification of non-oncogene drivers from gene-centric analysis of human cancer genomic data. Notably, our pathway analysis in oncogene-negative lung adenocarcinomas indicated that mutations in different genes that converge on the RAS and PI3K pathways frequently co-occur (**Figure 4i** and **S14i**).

Furthermore, previous reports and our observations suggest frequent non-genomic mechanisms of downregulation of RAS GAPs and PTEN (**Figure 4f-h, S14a-f**) ^4, 22–24, 53, 54^. Thus, genomic alterations should be viewed as a floor, not a ceiling, in estimating the frequency of pathway alteration.

We assessed the ability of hundreds of complex tumor suppressor genotypes to generate lung tumors. While activation of RAS and PI3K pathway emerged as the most potent driver of oncogene-negative lung adenocarcinomas, our data also suggest that combinatorial inactivation of tumor suppressor genes outside these two pathways can likely initiate tumorigenesis (**Figure 2** and **S6**). Given the mutational diversity and complexity of oncogene-negative human lung adenocarcinomas ^64^, there remain many other mutational combinations to be investigated. We anticipate that additional studies will uncover other oncogene negative tumor subtypes beyond Onc-negative^RAS/PI3K^ lung adenocarcinomas.

Knowledge of the genes underlying human cancer is a pillar of cancer diagnostics, personalized medicine, and the selection of rational combination therapies. Additionally, our data demonstrate RAS and PI3K pathway activation in the absence of oncogene mutations in a sizable fraction of human lung adenocarcinoma that could predict therapeutic vulnerability to SHP2 and AKT inhibitors. Thus, biochemical assessment of oncogenic pathways in tumors is a strong foundation for rational selection of therapies and clinical trial designs. Beyond SHP2 and AKT, extensive efforts have generated inhibitors for many other components of the RAS and PI3K pathways. Thus, further investigation of the therapeutic targeting of key nodes within the RAS pathway (*e.g.,* SOS, MEK, ERK) and PI3K pathway (*e.g.,* PI3K, mTOR), could contribute to the development of the most effective therapies for Onc-negative^RAS/PI3K^ lung adenocarcinomas.

Our findings uncover tumorigenic mechanisms and clinical features of oncogene-negative lung adenocarcinomas. This work identifies biomarkers and new therapeutic targets for Onc-negative^RAS/PI3K^ tumors. The generation of comprehensive molecular and pharmacogenomic maps of oncogene-negative lung adenocarcinomas will transform our understanding of these heretofore poorly characterized lung cancer subtypes.

## ACKNOWLEDGEMENTS

We thank the Stanford Shared FACS Facility for flow cytometry and cell sorting services, the Stanford Veterinary Animal Care Staff for expert animal care, Human Pathology/Histology Service Center, Stanford Protein and Nucleic Acid Facility; A. Orantes and S. Mello for administrative support; Stanford’s Molecular Genetic Pathology Laboratory and Henning Stehr for their help in providing genetically profiled tumor tissues. David Feldser, Joseph Lipsick, Eric Collisson, Christopher McFarland, and members of the Winslow and Petrov laboratories for helpful discussions and reviewing the manuscript. We thank Florent Elefteriou and Alejandro Sweet-Cordero for providing mouse strains. M.Y. was supported by a Stanford University School of Medicine Dean’s fellowship, an American Lung Association senior research training grant, and an NIH Ruth L. Kirschstein National Research Service Award (F32- CA236311). G.B., H.C., and J.D.H. were supported by a Tobacco-Related Disease Research Program (TRDRP) Postdoctoral Fellowships (T31FT-1772, 28FT-0019, and T31FT-1619). C.W.M. was supported by the NSF Graduate Research Fellowship Program and an Anne T. and Robert M. Bass Stanford Graduate Fellowship. W-Y.L. was supported by an American Association of Cancer Research Postdoctoral fellowship (17-40-18-LIN). C.L. was the Connie and Bob Lurie Fellow of the Damon Runyon Cancer Research Foundation (DRG-2331). E.L.A and C.I.C were supported by PHS Grant Number CA09302, awarded by the National Cancer Institute, DHHS. E.L.A. was also supported by HHMI Gilliam Fellowship for Advanced Study (GT14928). This work was supported by NIH R01-CA231253 (to M.M.W and D.A.P), NIH R01-CA230919 (to M.M.W.) and NIH R01-CA234349 (to M.M.W and D.A.P.), as well as by the Stanford Cancer Institute, an NCI-designated Comprehensive Cancer Center.

---

**Figure S1.**
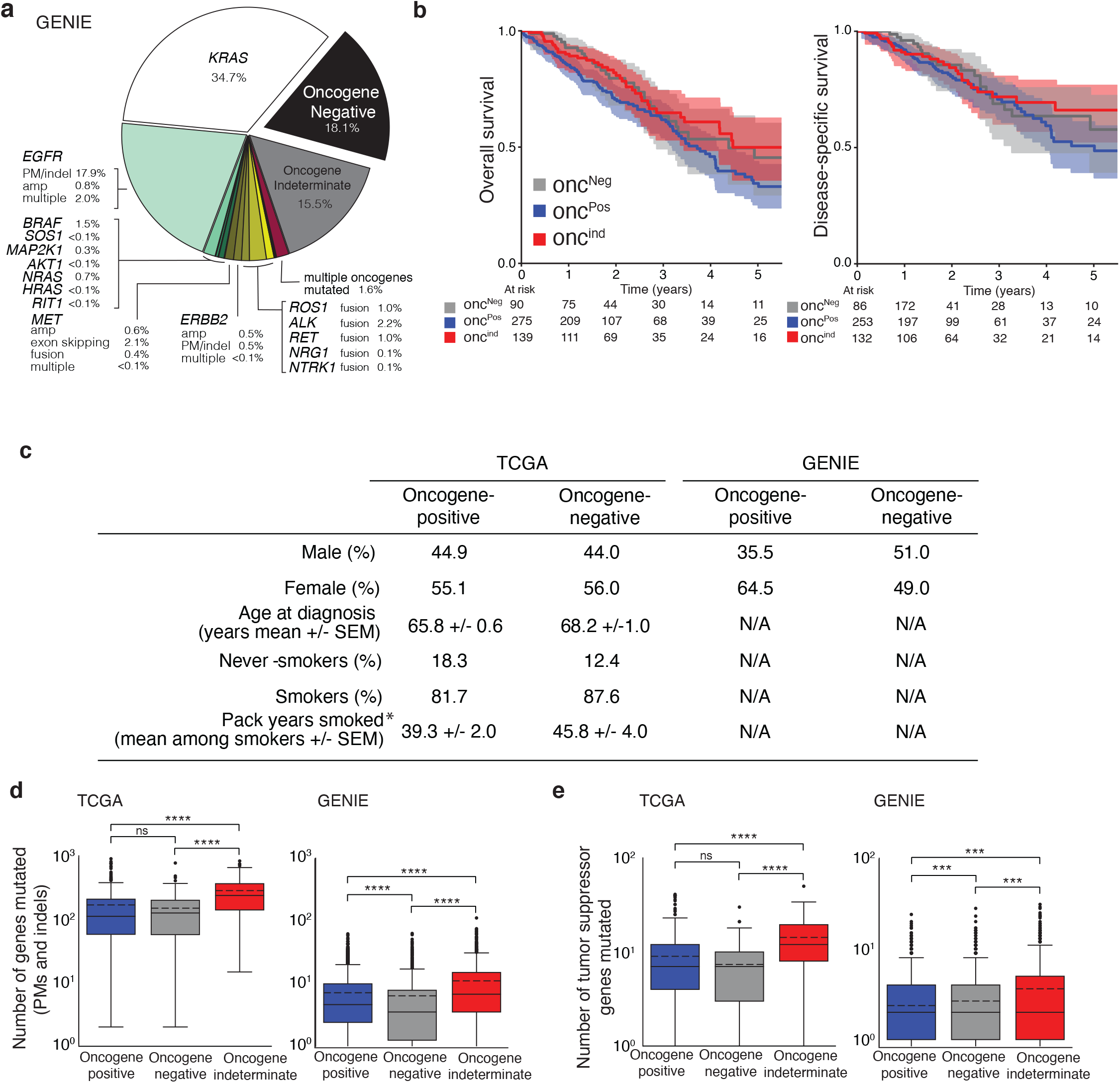
Clinical and molecular features of oncogene-negative human lung adenocarcinomas. **a.** Frequency of human oncogene-positive, oncogene-indeterminate, and oncogene-negative lung adenocarcinomas based on GENIE data sets. PM: point mutation, indel: insertion and deletion, amp: amplification, multiple: multiple oncogenic alterations in the same gene (see **Methods**) **b.** Overall survival and disease-specific survival of oncogene-positive (Onc^Pos^), oncogene-indeterminate (Onc^Ind^), and oncogene-negative (Onc^Neg^) lung adenocarcinoma patients based on TCGA data.The numbers below the plots are the numbers of patients alive at each time point. **c.** Clinical characteristics of oncogene-positive and oncogene-negative patients based on TCGA and GENIE data sets. SEM - standard error of the mean. N/A - information not present in this dataset. The p-values were calculated using Mann Whitney U test, * p<0.05 . **d,e.** The number of mutated genes (**d**,by point mutations (PMs) and indels) and number of mutated tumor suppressor genes (**e**, by point mutations, indels, or deletions) in oncogene-positive, oncogene-indeterminate, and oncogene-negative tumors based on TCGA and GENIE data sets. The mean is represented by the dashed line and the median by the straight line. ****p<0.0001, p-values were calculated using Mann Whitney U test.

**Figure S2.**
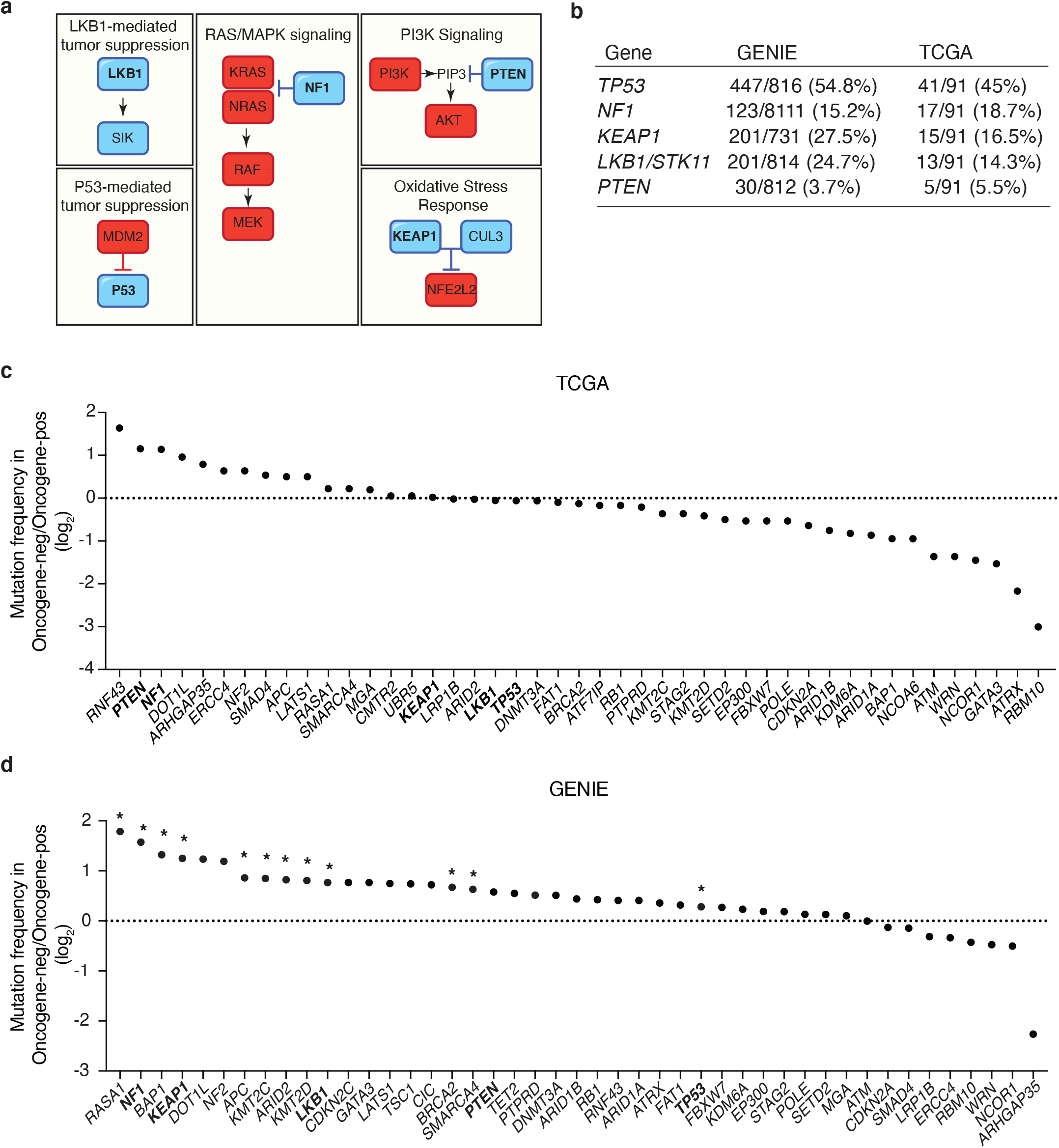
Tumor suppressor gene alterations in oncogene-negative lung adenocarcinoma. **a.** Schematic of the pathways controlled by the five tumor suppressor genes inactivated using floxed alleles (“core” tumor suppressor genes) in this study. The tumor suppressors represent different key cancer pathways. **b.** Alteration frequency of “core” tumor suppressor genes (number of tumors with potentially inactivating missense or nonsense mutations or focal DNA copy number losses/ total tumor number) in oncogene-negative lung adenocarcinomas based on GENIE and TCGA data sets. **c,d.** The ratio of the frequency of inactivating alterations of tumor suppressor genes (point mutations, indel, and copy number loss) of the genes in the Lenti-sg*TS102/Cre* and Lenti-sg*TS15/Cre pools* in oncogene negative versus oncogene-positive lung adenocarcinomas. Data from TCGA (**c**) and GENIE (**d**) data sets are shown. The dotted line represents equal frequency in oncogene-negative and oncogene-positive lung adenocarcinomas. The “Core” tumor suppressors are in bold. *FDR<0.05

**Figure S3.**
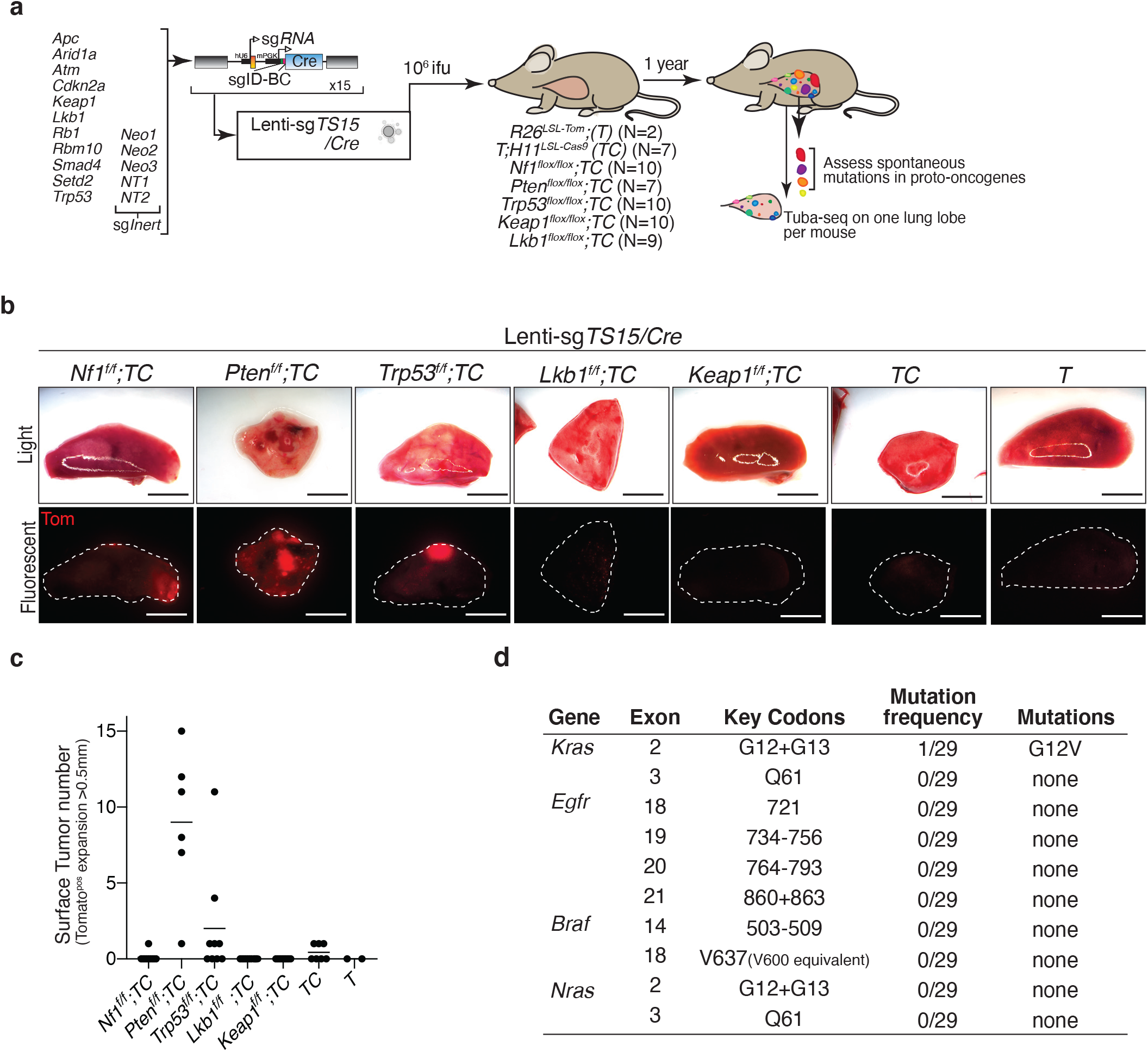
Most tumors in *Nf1^f/f^;TC*, *Pten^f/f^;TC*, and *Trp53^f/f^;TC* mice arise in the absence of mutations in the proto-oncogenes. **a.** Schematic of combined Cre/lox and CRISPR/Cas9-mediated tumor suppressor gene inactivation to generate lung epithelial cells with diverse genotypes. **b.** Representative light and fluorescence images of lung lobes from the indicated genotypes of mice. Lung lobes are outlined with white dotted lines. Scale bar = 4 mm **c.** The number of tumors (defined as Tomato-positive expansions greater than 0.5 mm in diameter) was quantified by direct counting. Each dot represents a mouse, and the bar is the mean. **d.** Exons in known proto-oncogenes that were analyzed by targeted sequencing. Key codons are those in which mutations are associated with oncogenic activity. Mutation frequency is the number of tumors with putative oncogenic mutations over the total number of samples analyzed. Putative oncogenic mutations are defined in **Methods**.

**Figure S4.**
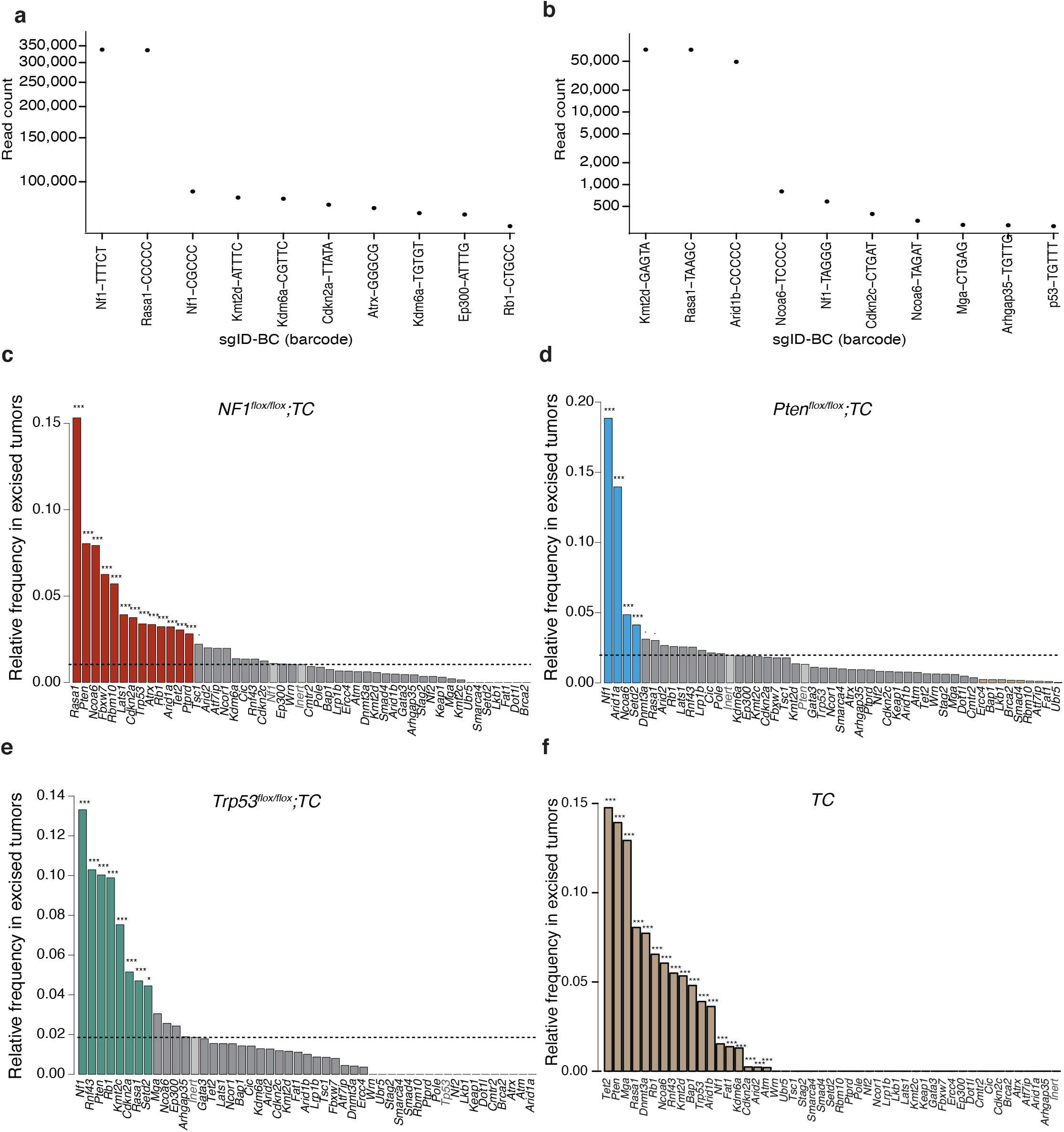
Identification of tumor suppressor genes that constrain lung tumor formation *in vivo*. **a,b.** Example plots indicating evidence of transduction with multiple barcoded lentiviral vectors. The10 sgID-BCs with the highest read counts from two excised tumors are shown. Each dot represents a sgID-BC, the y-axis shows read count, and the sgID-BCs are sorted on the x-axis by decreasing read counts (the first 5 nucleotides of the random barcode are shown with the targeted gene symbol). The first two and three barcodes (sgID-BC) in subpanels a and b, respectively, that have very similar read counts likely represent a single clonal tumor initiated from a cell transduced with multiple barcoded Lenti-sgRNA/Cre vectors (see **Methods** Multiple transduction section). **c-f.** Relative frequency of sgRNAs targeting each tumor suppressor gene in tumors harvested from the indicated genotypes of mice. Tumors were dissected under a fluorescence microscope based on their tdTomato fluorescent signal and were subjected to genomic DNA extraction. The sgID-BC region was PCR amplified and sequenced using Illumina high-throughput sequencing. The dotted lines represent the frequency of the inert sgRNA (average of all inert sgRNAs). Genes significantly overrepresented compared to the inert sgRNA are shown as: *** p < 0.001, * p < 0.05, · p < 0.1. Light gray bars indicate sgRNAs targeting the “core” tumor suppressor gene that is inactivated with floxed alleles in each plot and the inert sgRNAs.

**Figure S5.**
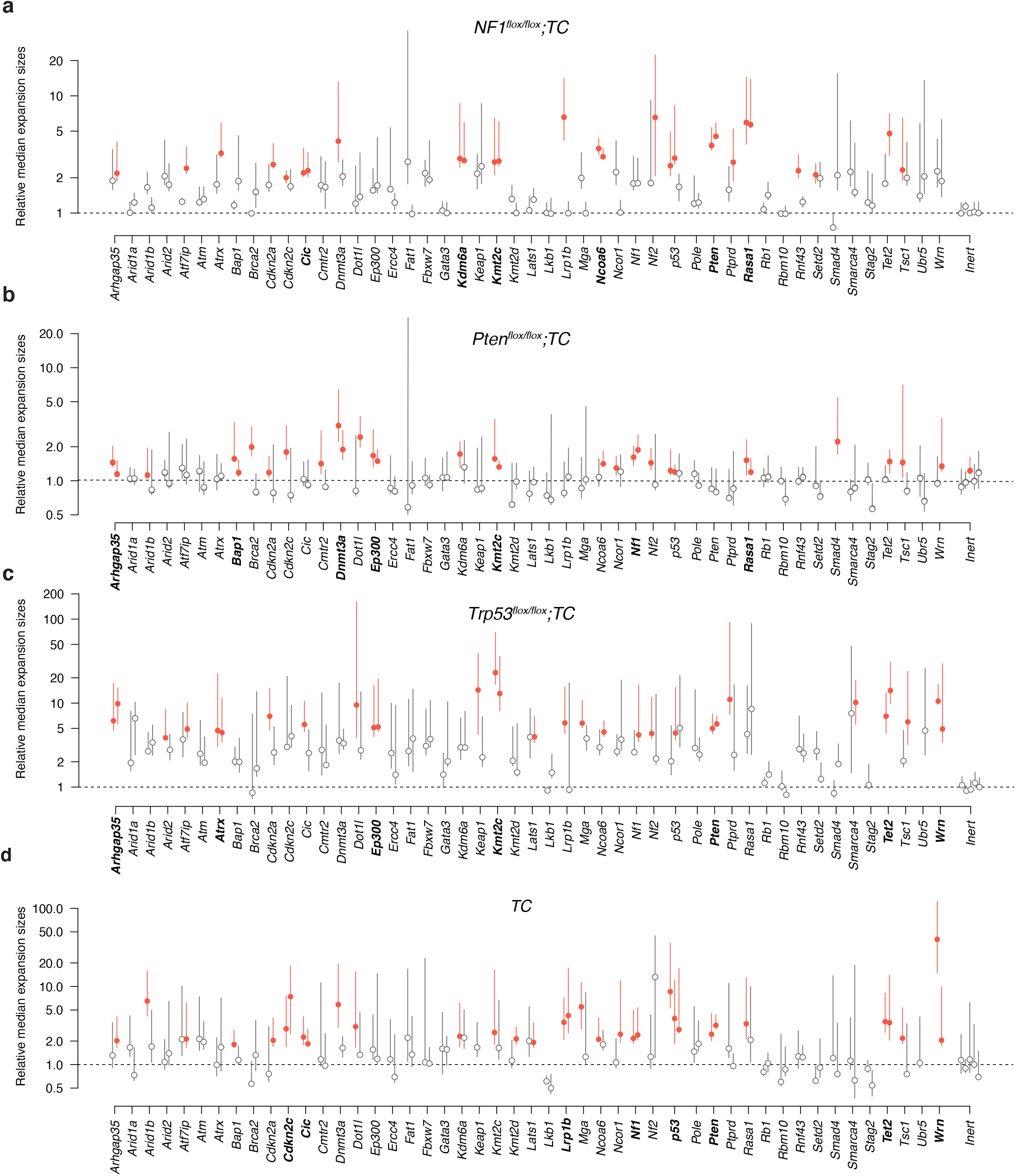
*In vivo* lung epithelial cell expansion is suppressed by diverse tumor suppressor genes. **a-d.** Median cell-expansion sizes (normalized to the sizes of inert sgRNA containing cell expansions) for each putative tumor suppressor targeting sgRNAs in one lung lobe harvested from the indicated mouse genotype are shown. Dotted lines indicate the median value for inert sgRNAs. Cell expansions are defined as clonal expansions containing a minimum of 50 cells. Bootstrap 95% confidence intervals are shown as whiskers. sgRNAs with median sizes significantly (p < 0.05) higher than the median effect for all sgRNAs are shown in red. Gene with significant effect upon inactivation using both sgRNAs are in bold.

**Figure S6.**
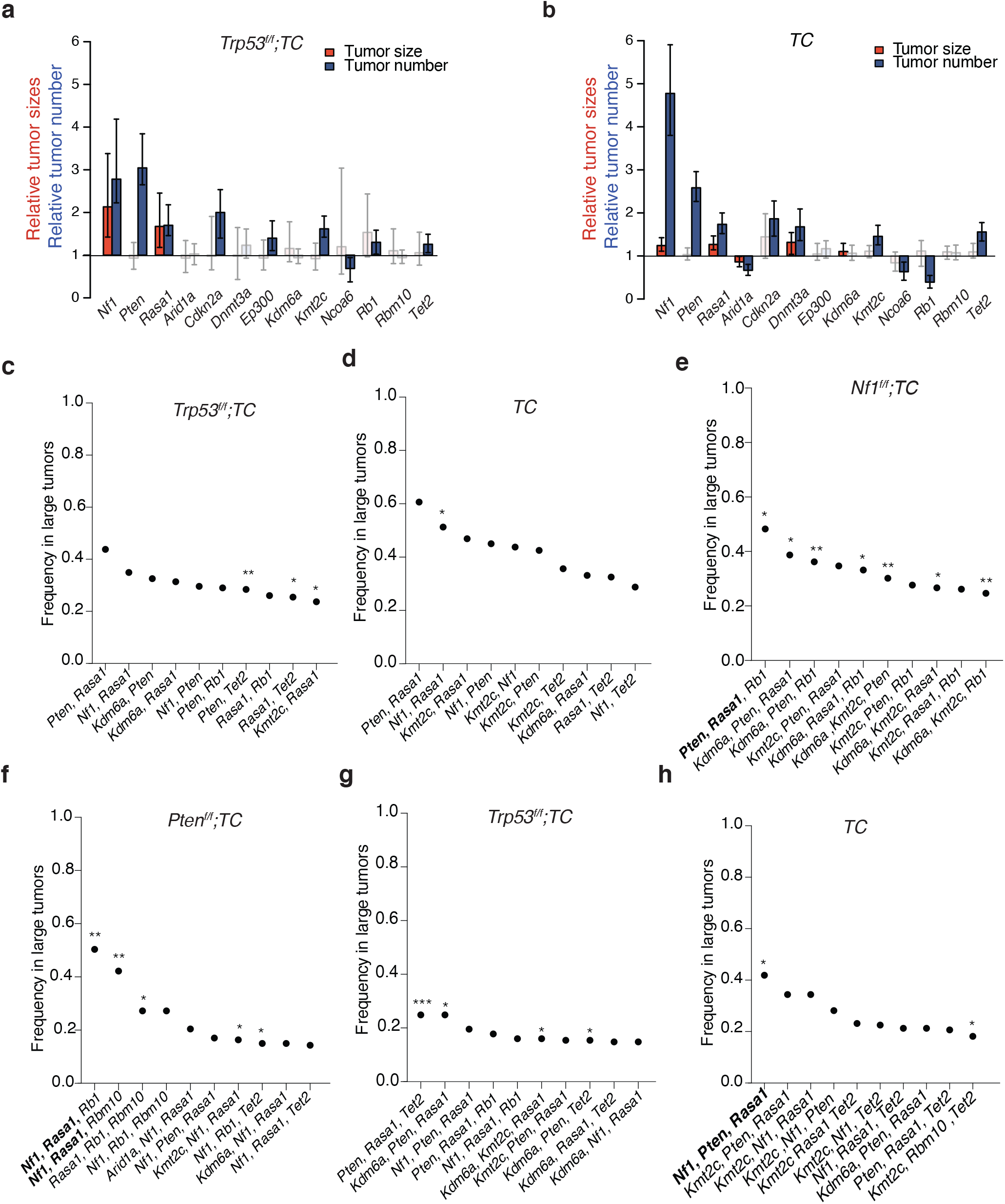
*Nf1*, *Rasa1*, and *Pten* are frequently targeted in the largest tumors of *Nf1^f/f^;TC, Pten^f/f^;TC, Trp53^f/f^;TC,* and *TC* mice. **a, b.** The number of tumors with a minimum size of 1000 cells relative to the inert guide is shown as a blue bar. 90^th^ percentile of tumor sizes relative to the inert sgRNA is shown as a red bar. sgRNAs resulting in a significantly higher number or larger tumors than the inert sgRNA (p<0.05) are shown in color. Whiskers show 95% confidence intervals. Mouse genotypes are indicated on each plot. **c-h.** Depiction of the top 10 most frequently co-occurring tumor suppressor alterations in each indicated genotype. Barcodes with the highest cell count in each mouse were investigated for coinfection for multiple viruses (see **Methods**). The top 10 pairs of tumor suppressors found co-mutated in the largest tumors are shown. *p<0.05, **p<0.01, ***p<0.001 based on a permutation test. Combinations of sgRNAs that lead to the generation of *Nf1*, *Rasa1*, and *Pten* mutant cancer cells in a statistically significant manner are in bold.

**Figure S7.**
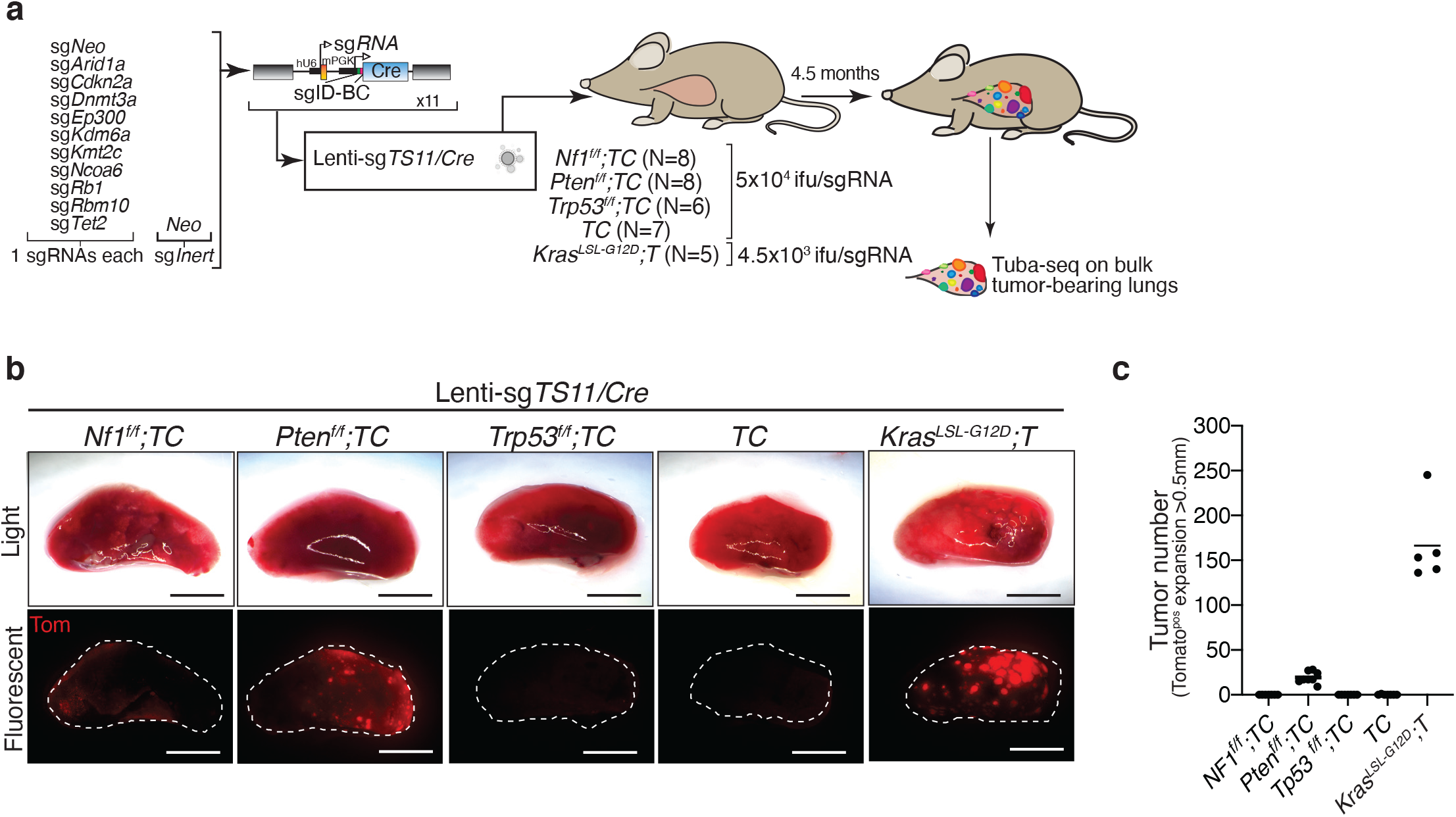
Very few tumors develop in *Nf1^f/f^;TC, Pten^f/f^;TC, Trp53^f/f^;TC,* and *TC* mice after delivery of the Lenti-sg*TS*11/*Cre* pool. **a.** Schematic of tumor initiation with a pool of 11 barcoded Lenti-sgRNA/Cre vectors (Lenti-sgTS11/Cre) similar to Lenti-sg*TS*14/*Cre* but excluding the Lenti-sgRNA/Cre vectors with sg*Nf1*, sg*Rasa1* and sg*Pten*. Each gene is targeted with a single sgRNA. Mouse genotype, mouse number, and titer of virus delivered to each group are indicated. Tuba-seq was performed on each tumor-bearing lung 4.5 months after tumor initiation. **b.** Representative light and fluorescence images of lung lobes from the indicated genotypes of mice. Lung lobes are outlined with white dotted lines. Scale bars=4mm **c.** The number of surface tumors (defined as Tomato-positive expansions larger than 0.5 mm in diameter) quantified by direct counting. Each dot represents a mouse, and the bar is the mean.

**Figure S8.**
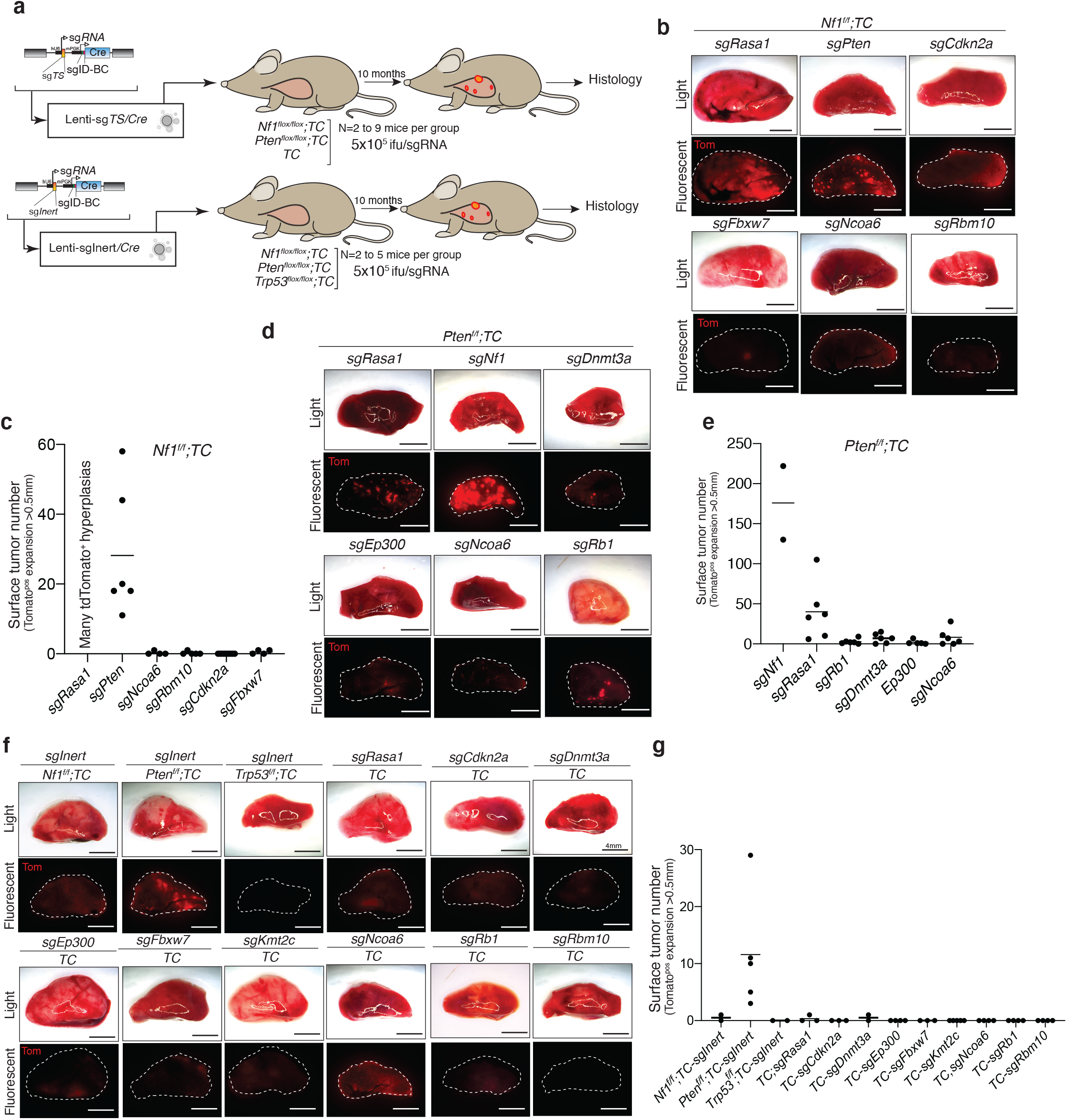
Single and Pairwise tumor suppressor gene inactivation is rarely sufficient to generate lung tumors. **a.** Schematic of experiments to assess the potential of single or paired tumor suppressor inactivation to generate lung tumors. **b,d,f.** Representative light and fluorescence images of lung lobes from the indicated genotypes of mice. Lung lobes are outlined with white dotted lines. Scale bar=4mm **c,e,g.** The numbers of tumors (defined as Tomato-positive cell expansions greater than 0.5mm in diameter) was quantified by direct counting. Each dot represents a mouse, and the bar is the mean. The genotypes of the recipient mice and the gene targeted by sgRNA are indicated. Inactivation of *Rasa1* in *Nf1^f/f^,TC* mice in **S8b,c** generated numerous tdTomato^+^ hyperplasias without distinguishable boundaries under the microscope. As a result, surface tumor number was not quantifiable for this group.

**Figure S9.**
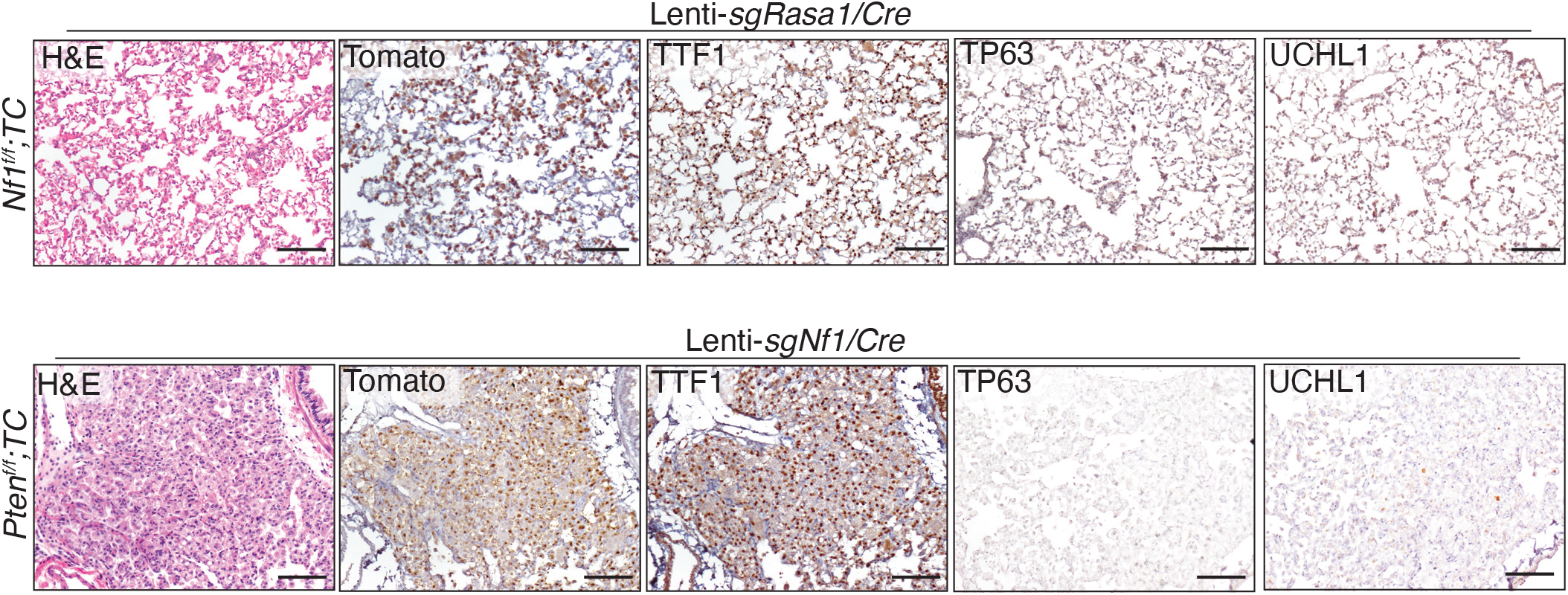
Pairwise inactivation of tumor suppressor genes rarely generates lung adenocarcinomas. Representative H&E, Tomato, TTF1, TP63, and UCHL1-stained sections of tumors from *Nf1^f/f^;TC* and *Ptenf^/f^;TC* mice 10 months after transduction with Lenti-*sgRasa1/Cre* or Lenti-*sgNf1/Cre.* Scale bars= 100 µm

**Figure S10.**
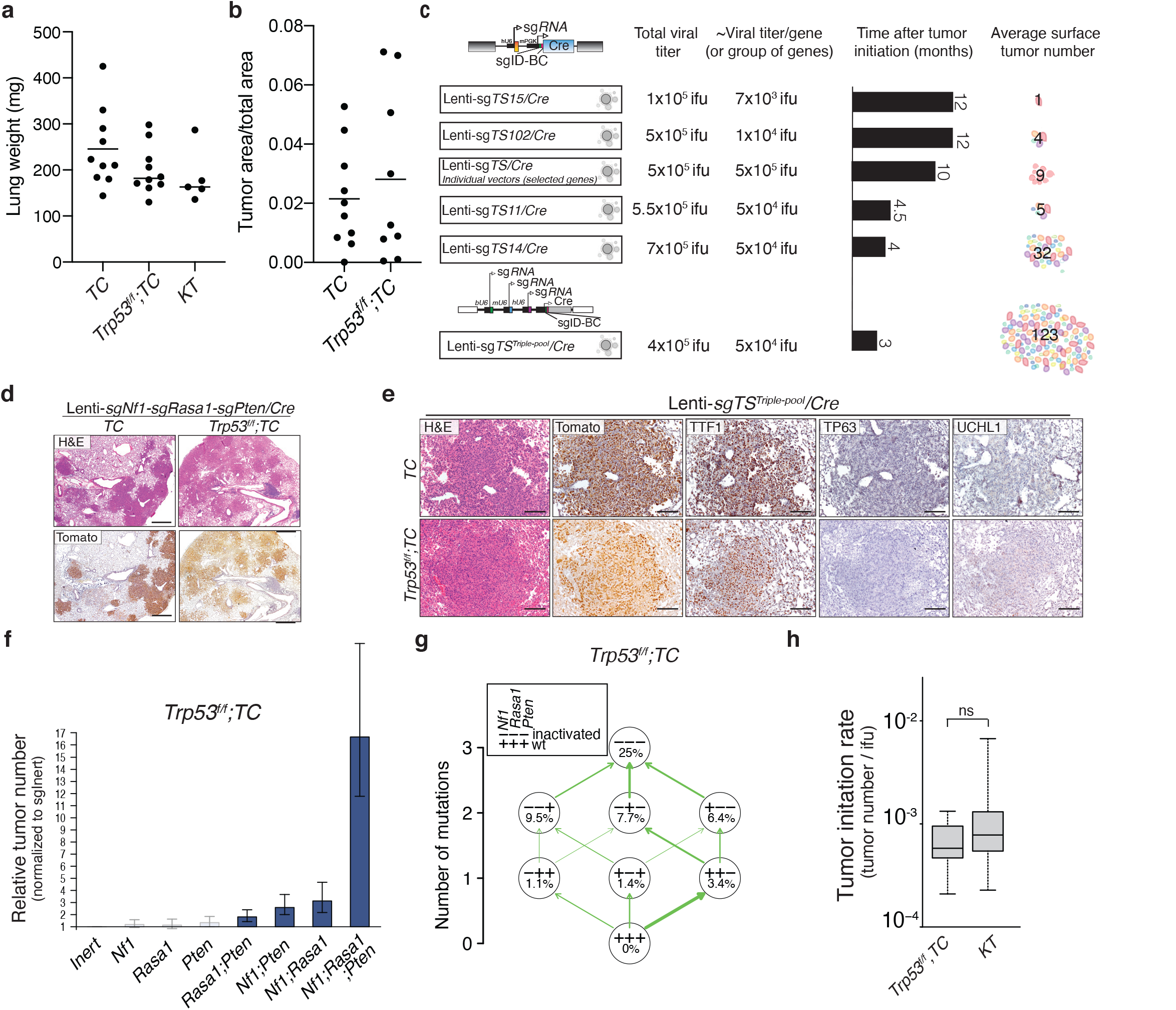
The relative contribution of *Nf1, Rasa1*, and *Pten* inactivation to oncogene-negative lung tumor development is not impacted by *Trp53* inactivation. **a.** Tumor burden, represented by lung weight. Each dot represents a mouse and the bar is the mean. **b.** Quantification of tumor burden based on H&E images. Each dot represents one lung lobe from each mouse, and the bar is the mean. **c.** Schematic of the increase in the number of oncogene-negative lung tumors generated in mice by enriching sgRNAs targeting the most potent tumor suppressor genes in each round of functional genomic screening *in vivo*. The viral titer, number of months after tumor initiation, and average number of tumors are indicated. **d.** Representative H&E and Tomato stained sections from *TC* and *Trp53^flox/flox^;TC* mice 3 months after transduction with Triple-Len- ti-sg*Nf1-*sg*Rasa1-*sg*Pten/Cre.* Scale bar= 500µm **e.** Representative H&E, Tomato, TTF1, TP63, and UCHL1-stained sections of tumors from *TC* and *Trp53^flox/flox^;TC* mice 3 months after transduction with Lenti-*sgTS^Triple-pool^/Cre.* Scale bar = 100 µm **f.** Numbers of tumors (with >1000 neoplastic cells) are shown relative to Inert sgRNA. sgRNAs resulting in a significantly higher number of tumors than sgInert (p<0.05) are shown in color. Mean +/- 95% confidence interval is shown. **g.** Adaptive landscape of *Nf1*, *Rasa1*, and *Pten* inactivation in *Trp53^flox/flox^;TC* is shown. Nodes represent genotypes shown as a string of +(wild-type) and - (inactivated) symbols representing *Nf1*, *Rasa1*, and *Pten*. Numbers in the nodes indicate fitness increase compared to wild-type. The relative probability of each beneficial mutation is shown as arrow widths (see **Methods**). **h.** Quantification of the ability of combined *Nf1/Rasa1/Pten* inactivation in *TC* mice and oncogenic Kras^G12D^ in *KT* mice to initiate tumors. Number of tumors (with >1000 neoplastic cells) per infectious unit (ifu) is shown. The bar is the median, the box represents the interquartile range, and the whiskers show minimum and maximum values. n.s: non-significant

**Figure S11.**
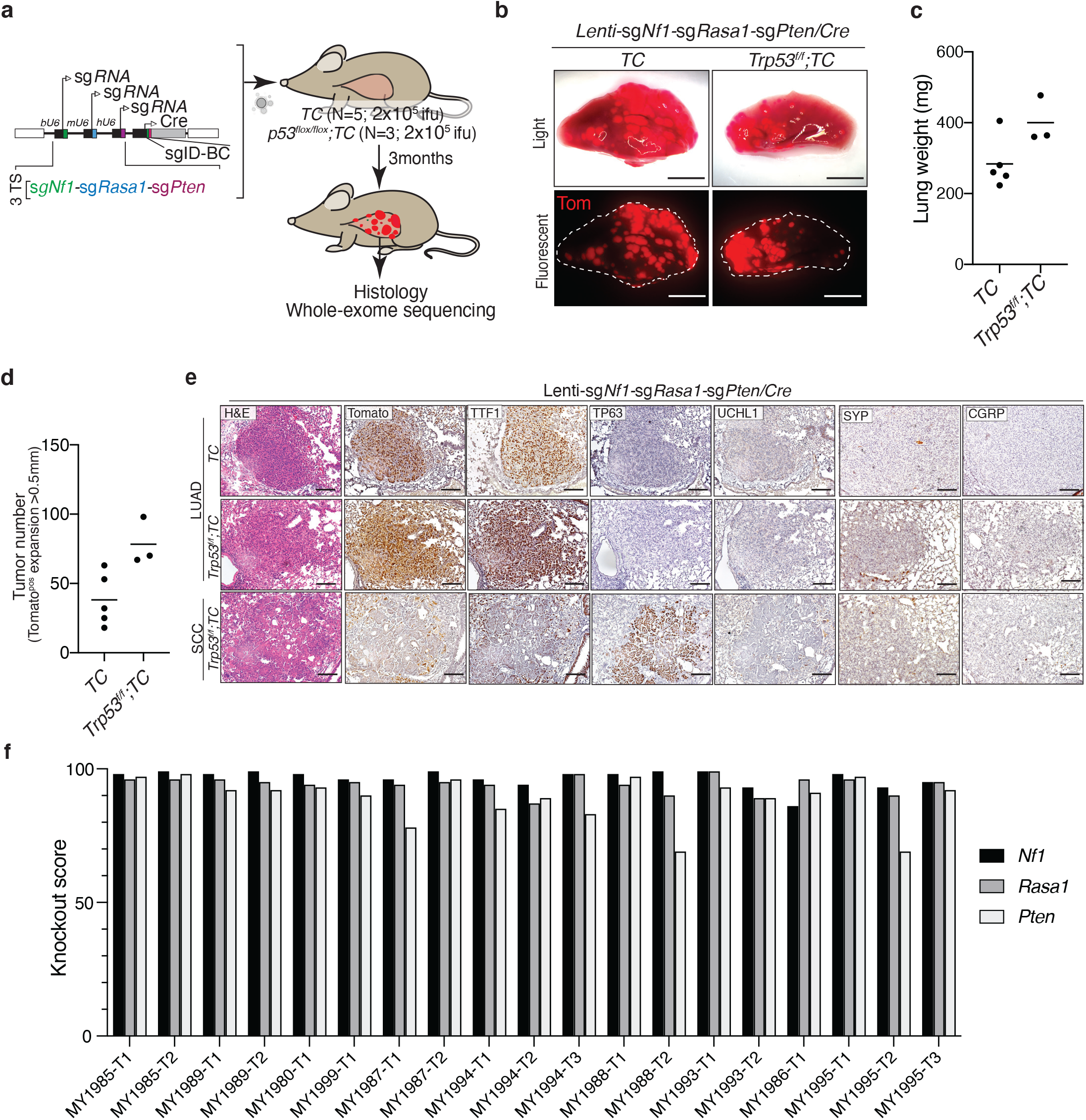
Oncogene-negative lung tumors driven by inactivation of *Nf1*, *Rasa1*, and *Pten* are almost exclusively adenomas/adenocarcinoma. **a.** Schematic of inactivation of *Nf1*, *Rasa1*, and *Pten* in *TC* and *Trp53^flox/flox^;TC* mice utilizing triple guide vectors and CRISPR/Cas9- mediated gene-inactivation. Mouse genotype, mouse number, and titer of virus delivered to each mouse are indicated. ifu, infection unit **b.** Bright-field and fluorescence images of lungs from the indicated mice 3 months after tumor initiation with Lenti-sg*Nf1*-sg*Rasa1*-sg*Pten*/*Cre* virus. Lung lobes are outlined with a dashed white line. Scale bars=4 mm **c.** Tumor burden, represented by lung weight. Each dot represents a mouse, and the bar is the mean. **d.** Quantification of tumor number based on H&E images of one lung lobe from each mouse. Each dot represents one lung lobe from each mouse, and the bar is the mean. **e.** Representative H&E, Tomato, TTF1, TP63, UCHL1, SYNAPTOPHYSIN (SYP), and CGRP-stained tumor sections from *TC* and *Trp53- ^flox/flox^;TC* mice 3 months after transduction with Lenti-*sgNf1-sgRasa1-sgPten/Cre.* Squamous cell lung cancer was only rarely observed in *Trp53^flox/flox^;TC* mice (3 out 264 tumors). Scale bars= 100 µm **f.** Analysis of insertion and deletion in genomic DNA from FACS sorted tumors of 19 TC mice 4 months after transduction with 5x10^4^ ifu of Lenti-*sgNf1-sgRasa1-sgPten/Cre*. sgRNA targeted regions were PCR amplified, and knockout scores, representing the proportion of cells that have either a frameshift-inducing indel or a large indel in a protein-coding region, were calculated using Synthego’s ICE.

**Figure S12.**
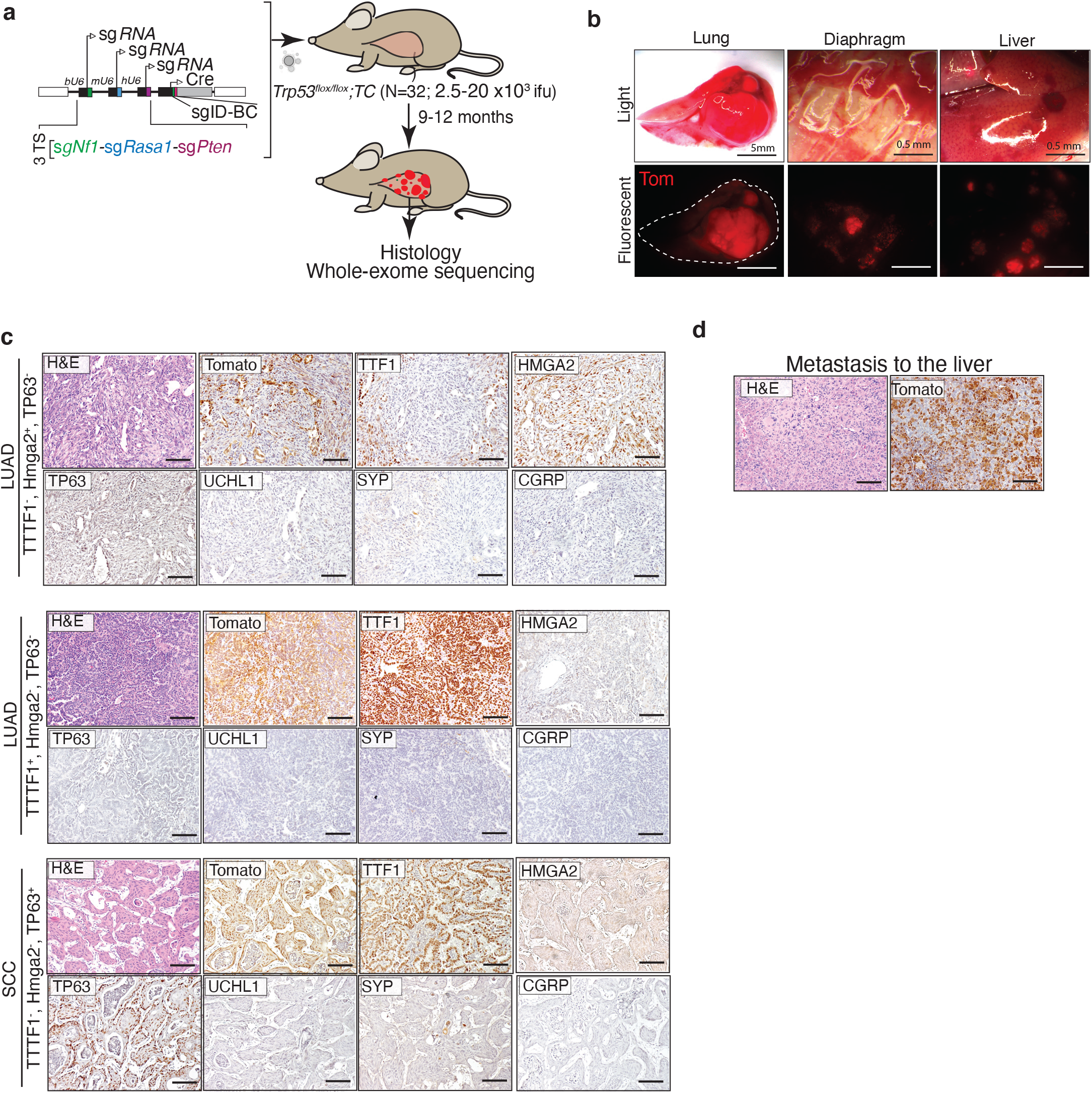
Inactivation of *Nf1*, *Rasa1*, and *Pten* generates lung tumors with the ability to metastasize to other organs. **a.** Schematic of inactivation of *Nf1*, *Rasa1*, and *Pten* in *Trp53^flox/flox^;TC* mice using the *Lenti-sgNf1-sgRasa1-sgPten/Cre* vector. Mouse genotype, mouse number, and titer of virus delivered mice are indicated. ifu, infection unit **b.** Bright-field and fluorescence images of lungs, diphragm, and liver from the *Trp53^flox/flox^;TC* mice 12 months after tumor initiation with Lenti-sg*Nf1*-sg*Rasa1*-sg*Pten*/*Cre* virus. Lung lobes are outlined with a dashed white line. Scale bars=5 and 0.5 mm (4 out of 32 mice had obvious metastasis). **c.** Representative H&E, Tomato, TTF1, HMGA2, TP63, UCHL1, SYNAPTOPHYSIN (SYP), and CGRP-stained tumor sections from*Trp53^flox/flox^;TC* mice 9-12 months after transduction with Lenti-*sgNf1-sgRasa1-sgPten/Cre.* Scale bars= 100 µm. **d.** H&E and tdTomato staining of liver sections from one of the *Trp53^flox/flox^;TC* mice with metastasis. Scale bars= 100 µm.

**Figure S13.**
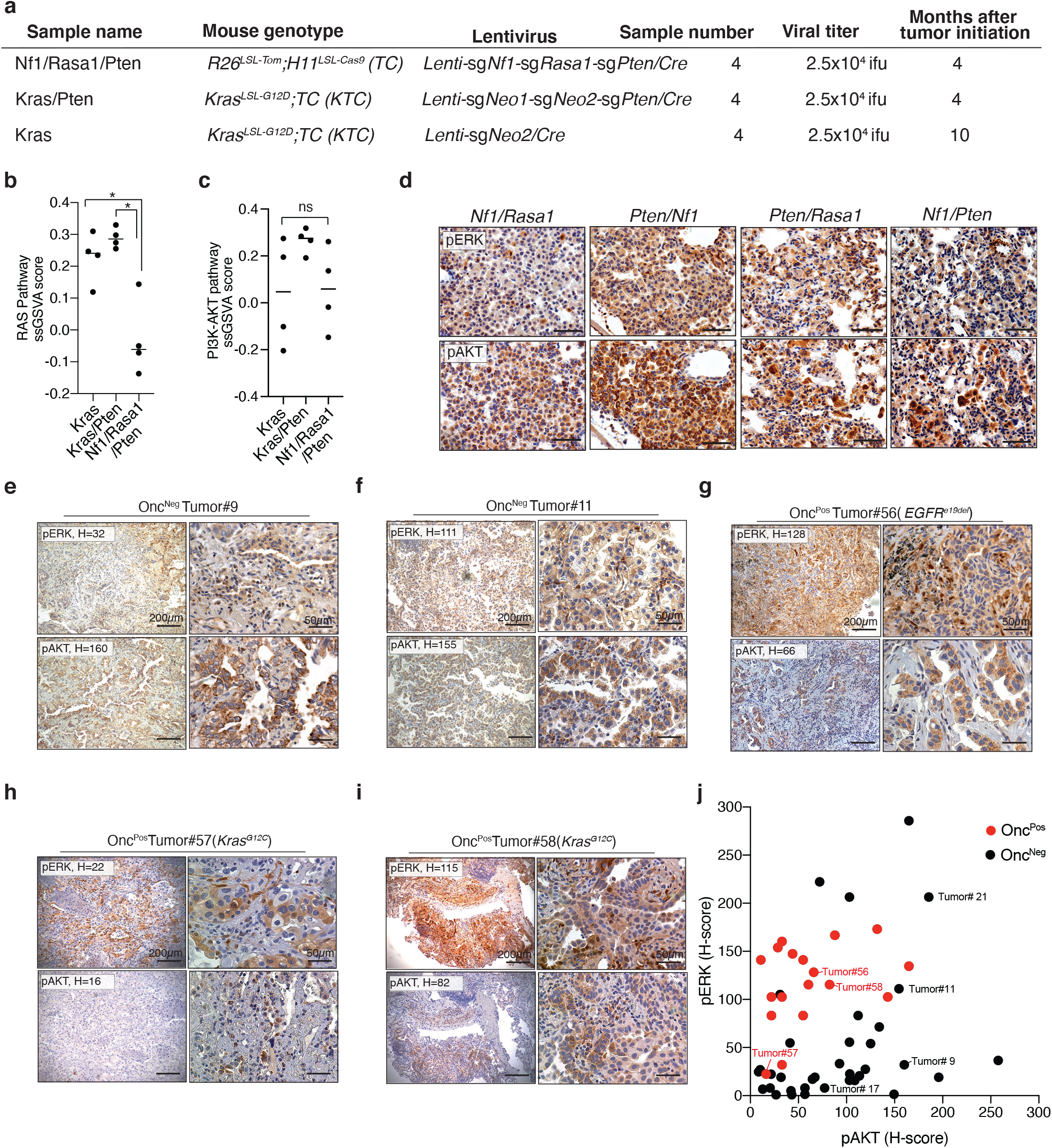
Onc-negative ^RAS/PI3K^ subtype of lung adenocarcinomas activate RAS and PI3K pathways biochemically. **a.** Summary of the mouse tumors sorted using FACS and analyzed by RNA-sequencing and immunohistochemistry. **b,c.** RAS and PI3K-AKT pathway gene-set profiles estimated by single-sample Gene Set Enrichment Analysis (ssGSVA). Tumors from *Kras^G12D^;TC* (*KTC*+ sg*Inert and KTC*+sg*Pten:* Kras and Kras/Pten) mice are compared with *Nf1*, *Rasa1*, and *Pten* mutant tumors(Nf1/Rasa1/Pten). The bar is the mean. ns: non-significant, *p<0.05 using Mann–Whitney U test. **d.** Representative immunohistochemistry for pERK and pAKT to determine activation of RAS and PI3K pathway in tumors with the indicated genotypes. The first gene is mutated using floxed alleles, and the second gene is inactivated using sgRNA/Cas9 (see **Figure S8** for more details). Scale bar = 50 µm **e-i.** Representative pAKT and pERK-stained sections of tumors from human oncogene-negative and oncogene-positive tumors. H-score for the whole section is indicated on each representative image. Scale bar= 200 µM (right), 50 µm (left) **j.** Replotting of pAKT and pERK staining on 35 oncogene-negative and 18 oncogene-positive human lung adenocarcinomas (Figure 4f**, g**). The tumors shown as IHC examples in Figure 4d**,e**, and **S14e-i** are labeled on this plot.

**Figure S14.**
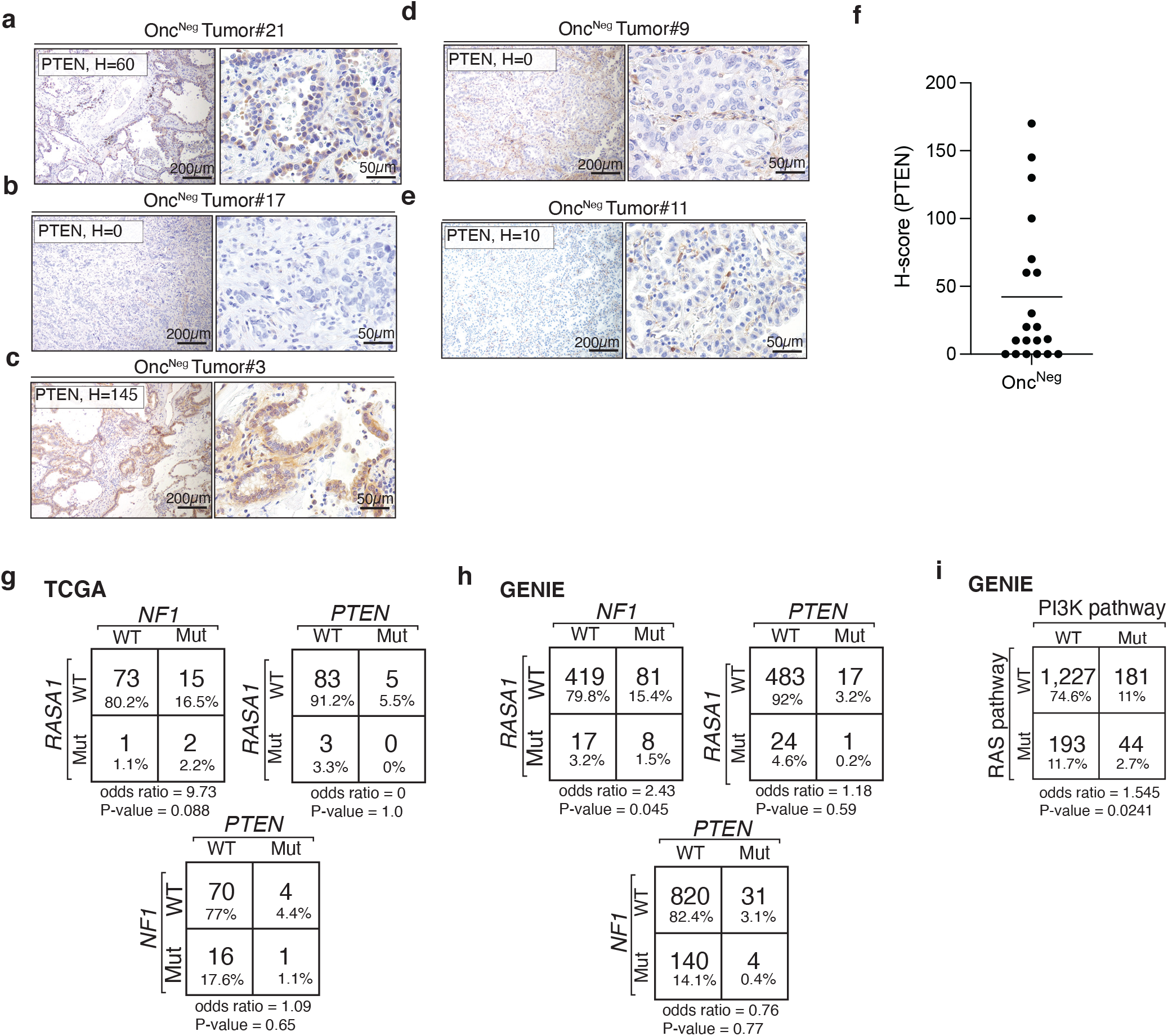
Alterations in RAS and PI3K pathways are enriched in Onc-negative ^RAS/PI3K^ subtype of human lung adenocarcinomas. **a-e.** Representative PTEN-stained sections of oncogene-negative human tumors. H-score for the whole section is indicated for each representative image. Scale bar= 200 µM (right), 50 µm (left) **f.** PTEN H-scores for oncogene-negative human lung adenocarcinoma tumors. **g, h.** Alteration frequencies of *NF1*, *RASA1*, and *PTEN* (point mutation, CNV, and indel) and assessment of their co-occurrences, the p-values were calculated by two-sided Fisher’s Exact Test. 91 oncogene-negative tumors were from the TCGA datasets. 525, 995, and 525 tumors were analyzed for *RASA1/PTEN*, *NF1/PTEN*, and *RASA1/NF1* alterations from the GENIE dataset. **i.** Frequency of alteration of well-established components of RAS and PI3K pathways (**Table S6**) queried in GENIE data set and their co-occurrences, the p-value calculated by two-sided Fisher’s Exact Test.

**Figure S15.**
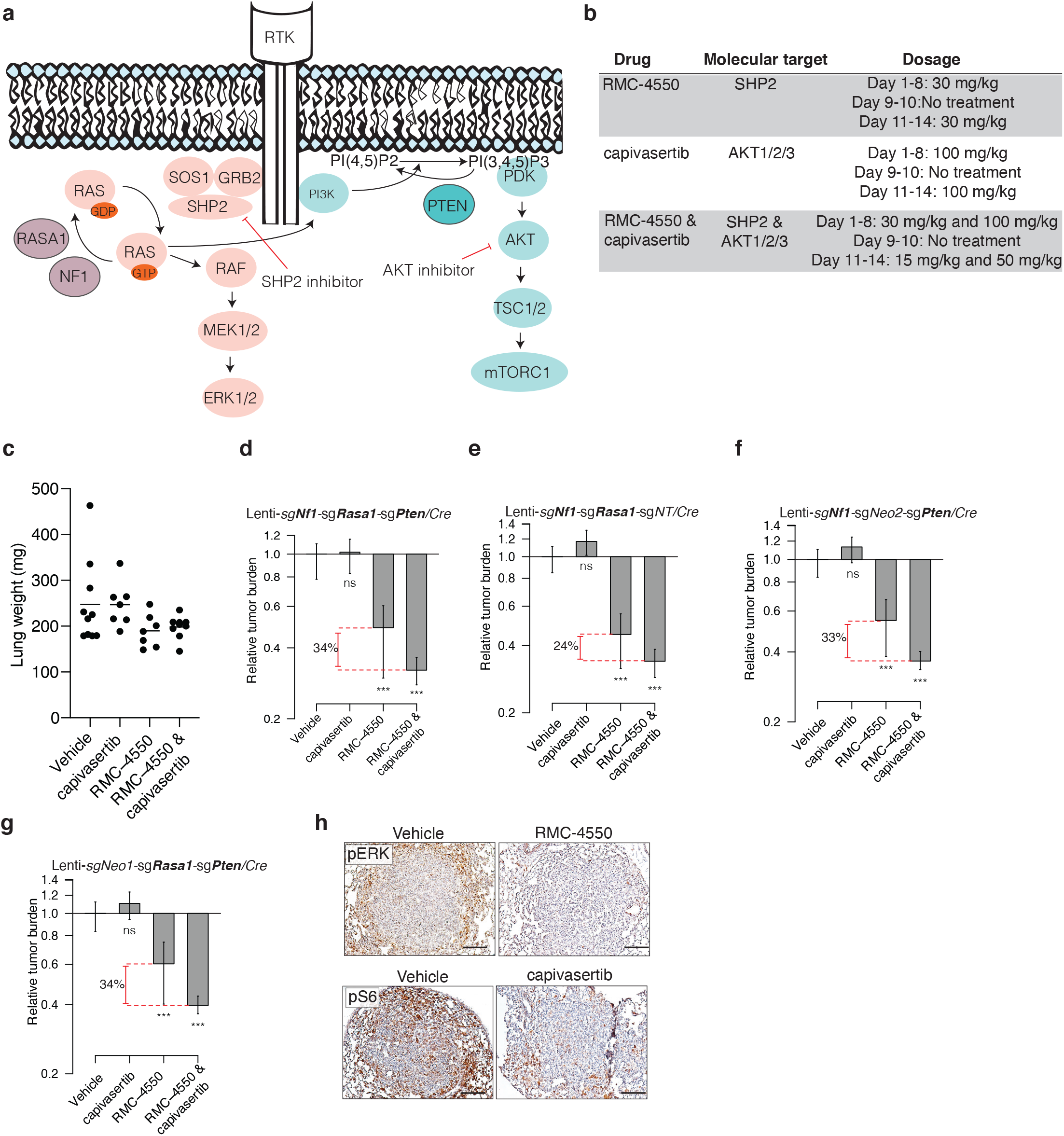
*Nf1*, *Rasa1*, and *Pten* mutant oncogene-negative lung tumors respond to inhibition of PI3K and RAS pathways. **a.** RAS and PI3K pathways are activated by alterations of *Nf1*, *Rasa1*, and *Pten* and targeted by SHP2 and AKT inhibitors. **b.** Drugs used to inhibit RAS and PI3K pathways *in vivo* and their dosages. **c.** Lung weight of mice described in Figure 5a-b. **d-g.** Relative tumor burden in mice after treatment with capivasertib, RMC-4550, and combination of these two drugs compared with tumor burden in vehicle-treated mice. ns: non-significant, ***p< 0.001. Drug response is shown for tumors driven by inactivation of different combinations of *Nf1*, *Rasa1*, and *Pten*. **h.** Representative pERK and pS6-stained sections of oncogene-negative^RAS/PI3K^ tumors from *TC* mice described in **S16** after treatment with the indicated drugs. The mice were injected with one last dose of indicated drugs 4 hours before tissue harvest. Scale bars= 100 µm

**Figure S16.**
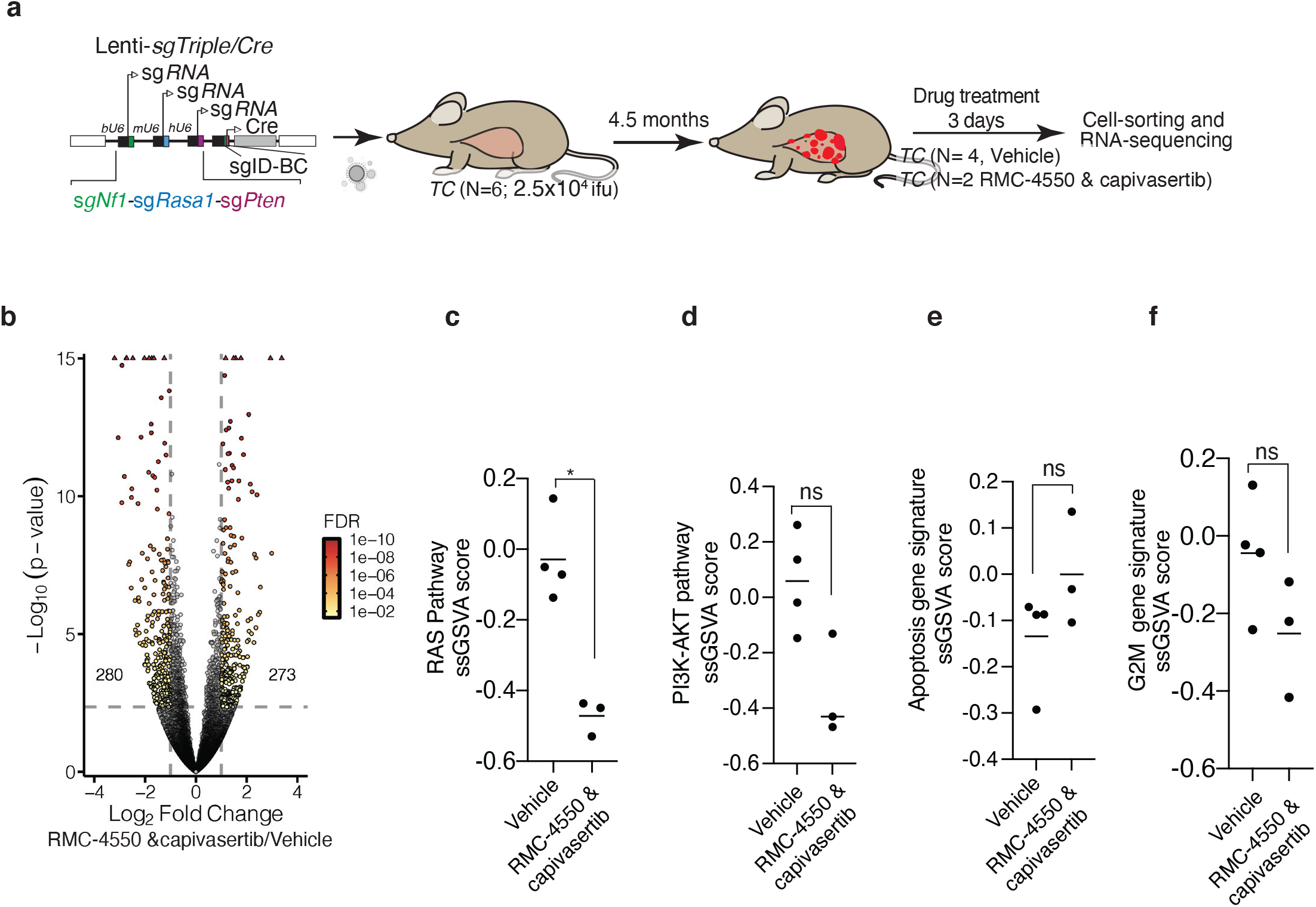
RMC-4550 and capivasertib treatment induces apoptosis gene signature and suppresses G2/M gene signature in Onc-negative^RAS/PI3K^ tumors. **a.** Generation of Onc-negative^RAS/PI3K^ tumors in *TC* mice to determine gene expression changes to pharmacological inhibition of RAS and PI3K pathways. Indicated number of mice were treated with vehicle or combination of RMC-4550 and capivasertib 4.5 months after tumor initiation for three days. RNA-sequencing was performed on sorted Tomato^positive^ epithelial cells in tumors. **b.** Volcano plots depicting a global overview of differential gene expression in Onc-negative^RAS/PI3K^ tumors in the absence and presence of treatment with RMC-4550 and capivasertib for three days as described above. Significant differential expression is defined as an absolute log2(Fold Change) > 1 and FDR < 0.01. The numbers of significantly differentially expressed genes are indicated on the plot. **c-f.** Comparison of RAS, PI3K-AKT, apoptosis, and G2M gene-set profiles estimated by single-sample Gene Set Enrichment Analysis (ssGSVA) in mouse Onc-negative^RAS/PI3K^ tumors after treatment with vehicle or RMC-4550 and capivasertib for three days. Each dot represents one tumor. ssGSVA data points shown for vehicle-treated tumors are the same as **Figure 13b, c** as Nf1/Rasa1/Pten. The bar is mean. ns: non-significant, *p<0.05 using Mann–Whitney U test.

**Figure S17.**
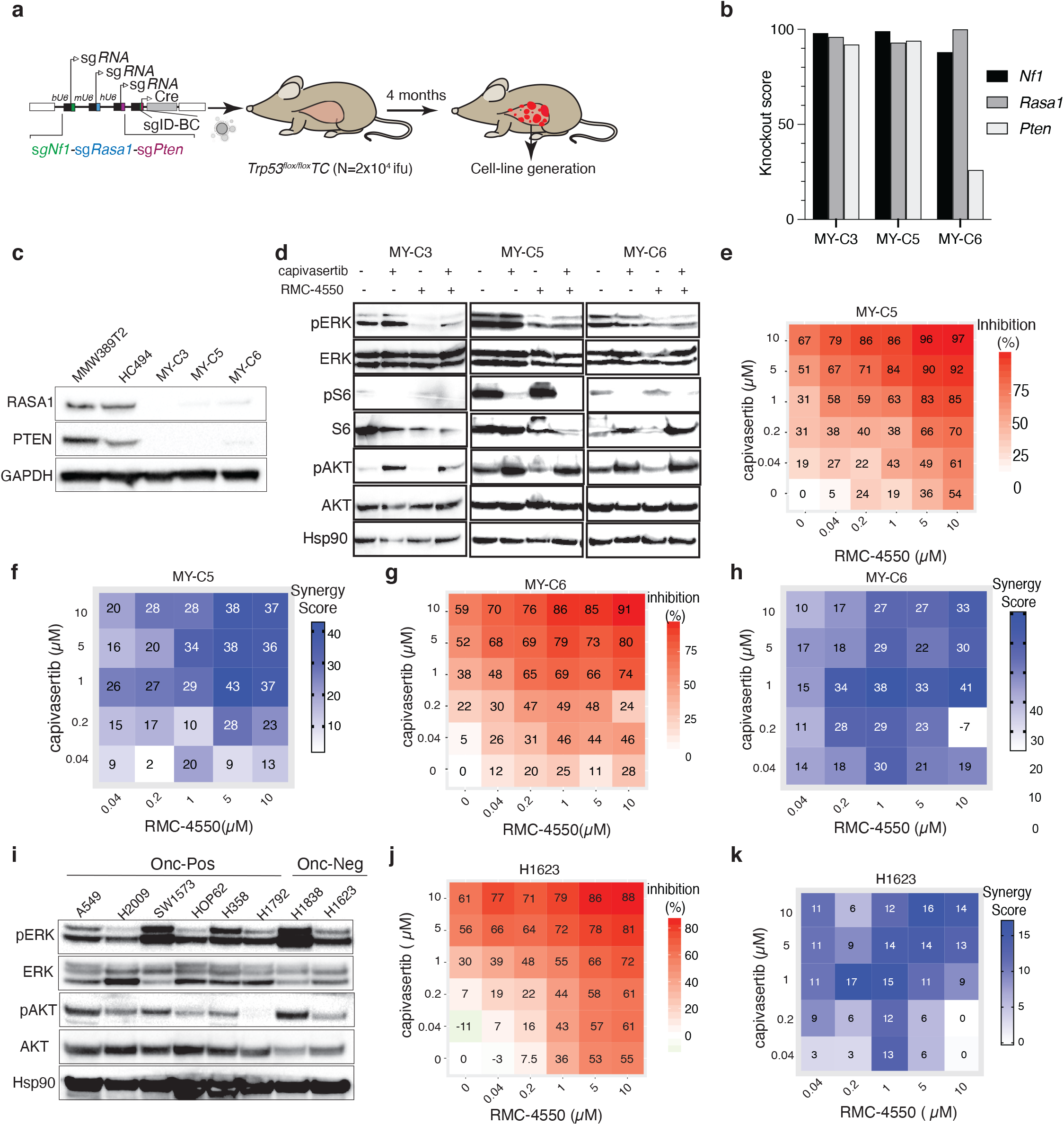
RMC-4550 synergizes with capivasertib to inhibit proliferation and induce cell death in Onc-negative^RAS/PI3K^ lung adenocarcinoma cell lines. **a.** Cell line generation from Onc-negative^RAS/PI3K^ tumors developed in *Trp53^flox/flox^;TC* mice. **b.** Indel analysis of 3 distinct mouse oncogene-negative^RAS/PI3K^ cell lines described above. Regions targeted by sg*Nf1*, sg*Rasa1*, and sg*Pten* were PCR amplified and analyzed using Synthego ICE after sanger sequencing. Knockout score represents indels causing frameshift mutations. **c.** Immunoblot of 2 murine oncogene-positive cell lines (MMW398T2 and HC494: *Kras^G12D^* and *Trp53* mutant and *Nf1*, *Rasa1*, and *Pten* wild type) and 3 murine Onc-negative^RAS/PI3K^ mouse cell lines (described above) to assess loss of RASA1 and PTEN in oncogene-negative cell lines. **d.** Immunoblot of 3 distinct oncogene-negative cell-lines treated with 10uM of indicated drugs for 24 hours. **e,g.** Drug dose-response matrix depicting % growth inhibition after treatment with various doses of RMC-4550 and capivasertib indicated on the plots. The cell-line used for the generation of each matrix is noted on top of each heatmap. **f,h.** Loewe’s synergy score was calculated for each drug dose combination shown in **e** and **g**. Synergy score indicates the percentage of inhibition beyond what is expected if there is no interaction between the drugs. **i.** Immunoblot of 6 human oncogene-positive and 2 human oncogene-negative cell lines for markers of RAS and PI3K pathway activation. **j.** Drug dose-response matrix depicting % growth inhibition of H1623 human Onc-negative^RAS/PI3K^ cell line. **k.** Loewe’s synergy score calculated based on drug responses in **Figure S17j**.

## METHODS

### Analysis of human lung adenocarcinoma datasets

Somatic mutation data (SNPs and indels, including silent mutations) for 513 TCGA lung adenocarcinoma (LUAD) tumors were downloaded from the UCSC Xena Browser (http://xena.ucsc.edu/) (Link 1 below). TCGA-LUAD clinical and exposure data were downloaded from the GDC Data Portal (https://portal.gdc.cancer.gov/projects/TCGA-LUAD) and the UCSC Xena Browser (Link 2 below). GISTIC2 thresholded copy number variation (CNV) data were downloaded from the UCSC Xena Browser (Link 3 below). Amplifications were defined as “2” and deletions as “-2”. Genes with conflicting CNV values within a single tumor were ignored. Fusion data were obtained from ^1^. Fusion and CNV data were filtered to include only data from the 513 samples within the somatic mutation set. Duplicate fusions were collapsed into single fusions. MET-exon skipping data were taken from ^2^. Curated survival data from ^3^ were downloaded from the UCSC Xena Browser (Link 4 below).

Links:

1. https://tcga.xenahubs.net/download/mc3/LUAD_mc3.txt.gz
2. https://tcga.xenahubs.net/download/TCGA.LUAD.sampleMap/LUAD_clinicalMatrix
3. https://tcga.xenahubs.net/download/TCGA.LUAD.sampleMap/Gistic2_CopyNumber_Gistic2_all_thresholded.by_genes.gz
4. https://tcga.xenahubs.net/download/survival/LUAD_survival.txt.gz

Data from AACR Project GENIE (hereinafter referred to as GENIE) v8 were downloaded from https://www.synapse.org/#!Synapse:syn22228642 ^3^, specifically: somatic mutations, copy number alteration (CNA) data, fusion data, panel information (genomic_information.txt), and clinical data (both sample- and patient-level). All data were filtered to only include LUAD tumors. A single tumor was kept for patients with multiple different tumor samples, with priority for earlier sequenced samples and those from primary tumors. If tumor samples appeared identical within the clinical meta-data, the related patient data were excluded.

### Determination of oncogenes

To have a conservative estimate of the fraction of lung adenocarcinomas without known oncogenic drivers (oncogene-negative tumors), we generated a list of oncogenes that included any gene that met at least one of these criteria: 1) Genes that have hotspot mutations or specific alterations where cancers or cancer cells with that mutation respond to therapies targeted to the protein product of that mutant gene in patients, 2) The particular alteration in that gene can generate lung adenocarcinoma in genetically-engineered mouse models, 3) The altered gene can generate tumors in other tissues in genetically-engineered mouse models, and 4) Alteration of the indicated gene can lead to the transformation of cells or predicts response to targeted therapies *in vitro*. Additionally, we excluded genes if their supposed oncogenic alterations co-occur with alterations in other proto-oncogenes (listed below) in more than 50% of cases.

| Gene | Oncogenic alterations | Patients respond to oncogene inhibition | Sufficient to generate lung tumors in GEMMs | Oncogene in GEMMs of other tumor types | Cellular transformation and/or drug response in cultured cells | Co-occurring with other proto-oncogenes listed here |
| --- | --- | --- | --- | --- | --- | --- |
| *KRAS* | Mutations at codons 12,13, and 61 | 4, 5 | 6-8 | 7, 9, 10 |  | 4% |
| *EGFR* | Diverse mutations and deletions | 11, 12 | 13-15 | 16 |  | 9% |
| *BRAF* | Diverse mutations and fusions | 17 | 18, 19 | 20, 21 | 22 | 18% |
| *HRAS* | Mutations at codons 12, 13, and 61 |  |  | 23 | 24, 25 | 13% |
| *NRAS* | Mutations at codons 12, 13, and 61 |  |  | 26, 27 |  | 14% |
| *MET* | Exon skipping | 28, 29 | 2 |  | 2, 30, 31 | 47% |
| *MEK1* | Diverse mutations |  |  |  | 32 | 24% |
| *SOS1* | Diverse mutations |  |  |  | 33 | 17% |
| *ALK* | Translocations/fusions | 34 | 35, 36 | 37 | 38 | 2% |
| *RET* | Translocations/fusions | 39 | 40, 41 |  |  | 10% |
| *ROS1* | Translocations/fusions | 42 |  | 43, 44 |  | 4% |
| *NTRK1* | Translocations/fusions | 45 |  |  | 46 | 29% |
| *NRG1* | Translocations/fusions | 47 |  |  |  | 0% |
| *AKT1* | Point mutation (E17K) | 48 |  |  | 49 | 50% |
| *RIT1* | Diverse mutations |  |  |  | 50 | 33% |
| *ERBB2* | Amplification, point mutation, deletion | 51 | 52, 53 | 54, 55 | 56 | 19% |
| *PIK3CA* | Diverse mutations |  | Opposing evidence<sup>57, 58</sup> |  |  | 68% |

### Classification of mutations and tumors

Mutations (somatic mutations, fusions, CNVs, and MET exon skipping [TCGA only]) were classified as within proto-oncogenes (described above) or not. Mutations within these proto-oncogenes were classified as “accepted oncogenic” mutations if those alterations met at least one of the criteria described above. Any tumor with one accepted oncogenic alteration was classified as “*oncogene-positive*”. Tumors with accepted oncogenic mutations in more than one gene were classified as “multiple oncogenes mutated”. Any tumor with alterations in a proto-oncogene that was not considered an accepted oncogenic alteration based on the four criteria above was classified as “*oncogene-indeterminate*”. Thus, these tumors contain variants of unknown significance (VUS) in proto-oncogenes ^59^. The remaining tumors, without any mutations in any proto-oncogene, were classified as “*oncogene-negative*”.

Tumor type counts per database:

|  | TCGA | GENIE |
| --- | --- | --- |
| Total | 513 | 9,099 |
| Oncogene-negative | 91 | 1,645 |
| Oncogene-positive | 283 | 6,041 |
| Oncogene-indeterminant | 139 | 1,413 |

Oncogene-positive tumors were further classified by the type of oncogenic mutation they had (**Figure 1a** and **S1a**).

### Clinical characteristics

We divided patients into males or females based on the sex reported by either TCGA or GENIE, if provided. For TCGA, the arithmetic mean for age at diagnosis was computed and reported with a standard error of the mean (SEM). Non-smokers were defined as having tobacco smoking history values of 1 (see public ID 2181650 at https://cdebrowser.nci.nih.gov), while smokers were defined as anything > 1 (current or reformed smokers). The arithmetic mean pack-years smoked for smokers, if reported, was described, along with SEM.

### Pan-cancer tumor suppressor genes

We generated a list of tumor suppressor genes based on two previously published reports to compare the number of altered tumor suppressor genes in oncogene-negative tumors with oncogene-positive and oncogene-indeterminate tumors ^60, 61^. We manually removed genes with conflicting evidence as a tumor suppressor gene in LUAD. The final list of TSGs is in **Table S1**.

### Calculation of mutation frequencies and absolute number of genes mutated

In general, mutation frequencies for a given gene were calculated as the number of tumors with that gene mutated, divided by the number of tumors screened for mutations in that gene (for TCGA: all tumors were screened for all genes, for GENIE: the panel sequencing information was obtained from genomic_information.txt to determine which tumors were screened for which genes). Mutation frequencies were calculated for point mutations (PM), insertion/deletions (indels), and deletions separately. Additionally, the frequency for a combination of PMs, indels, and deletions was also calculated. The screened set of tumors in GENIE for the latter included only those tumors which were screened for both PMs/indels as well as CNVs for each gene. Reported in **Figure S2b** are oncogene-negative tumors with either point mutations, indels, or deletions in the indicated gene. In **Figure S2c-d**, for each gene, a ratio of enrichment of mutations in oncogene-negative over oncogene-positive tumors was calculated as:

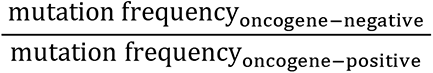

The *p*-values for enrichments were calculated using the two-sided Fisher’s Exact test as implemented by SciPy. For a given set of genes with at least a single tumor screened, the false discovery rate (FDR) was calculated using the Benjamini-Hochberg method on the Fisher’s Exact *P*-values.

To measure the total number of genes mutated (**Figure S1d**), a gene was considered mutated if it had at least one point mutation or indel. All these mutations in a tumor were collated, and the number of the unique set of genes was counted as the total number of genes mutated. For counting the number of individual tumor suppressors mutated (**Figure S1e**), deletions were also included, and the list of pan-cancer tumor suppressors as defined above was used. The Mann-Whitney U test was conducted on the number of respective genes mutated in either oncogene-negative or oncogene-positive tumors.

### Survival Analysis

Survival data from ^3^ were obtained as described above. Kaplan-Meier analysis was performed to estimate probability curves for overall survival (OS) and disease-specific survival (DSS). The logrank test was used to compare oncogene-negative and oncogene-positive tumors.

### Gene and pathway alteration co-occurrences

For analysis of simultaneous pairwise alterations of *NF1*, *RASA1*, or *PTEN* within oncogene-negative tumors, we determined the number of tumors with no mutation in *NF1, RASA1*, or *PTEN,* with mutation(s) in one gene, or mutations in two genes. Point mutations, indels, and deletions in each gene were included. A tumor needed to have one or more mutations in that gene to be considered mutated. For GENIE, only those tumors screened for both genes for point mutations and indels (according to the panel information file) were investigated. For TCGA, all oncogene-negative tumors were considered. A one-sided Fisher’s exact test was conducted to determine if there were more than the expected number of tumors with both genes mutated.

Gene lists and their acceptable alterations (*i.e*., not known to be an oncogene alteration) were generated as being in RAS or PI3K pathways ^60–80^ (**Table S6)**. We determined the list of all tumors screened for each gene in each pathway for the respective type of mutation (point mutation/indel, amplification, deletion, or fusion). For each alteration within each pathway, we determined whether it could activate the corresponding pathway or not according to the above list. A gene was considered mutated if it had at least one accepted mutation within it. A tumor was considered mutated in a given pathway if it had at least one gene mutated in that pathway.

### Animal Studies

The use of mice for the current study has been approved by Institutional Animal Care and Use Committee at Stanford University, protocol number 26696. *Kras^LSL-G12D/+^* (Jax # 008179 (K))*, R26^LSL-tdTomato^(ai9)* (Jax # 007909 (T)), and *H11^LSL-Cas^*^9^ (Jax # 026816 (C)), *Keap1^flox^, Pten ^flox^* (Jax # 006440)*, Lkb1 ^flox^* (Jax # 014143)*, Nf1 ^flox^ (*Jax # 017640), and *Trp53^flox^* (Jax # 008462) mice have been previously described ^6, 81–87^. All mice were on a C57BL/6:129 mixed background except the mice used for derivation of oncogene-negative *Nf1*, *Rasa1*, *Pten, and Trp53* mutant cell-lines, and some of the *Trp53^flox/flox^;TC* mice that were used for metastasis analysis (**Figure S12a**), which were on a pure C57BL/6 background.

### Tumor initiation and selection of Lenti-sgRNA*/Cre* pools

Tumors were initiated by intratracheal delivery of pooled or individual Lenti-sgRNA*/Cre* vectors. Barcoded Lenti-sgRNA*/Cre* vectors within each viral pool are indicated in each figure. Tumors were initiated with the indicated titers and allowed to develop tumors for between 3 and 12 months after viral delivery, as indicated in each figure.

In **Figure 1** and **Figure S3**, we transduced *Nf1^f/f^;TC, Pten^f/f^;TC, Trp53^f/f^;TC, Lkb1^f/f^;TC, Keap1^f/f^;TC, TC*, and *T* mice with two pre-existing pools of barcoded Lenti-sgRNA/Cre vectors that target ∼50 putative tumor suppressor genes. These two pools have been previously used to studied the effect of these putative tumor suppressor genes in KRAS^G12D^-driven lung tumors (Lenti-sg*TS*15*/Cre* ^88, 89^ and Lenti-sg*TS*102*/Cre* ^90^).

Lenti-sg*TS*15*/Cre* contained vectors targeting 11 tumor suppressors with one sgRNA per gene in addition to four inert sgRNAs (Lenti-sg*TS*15*/Cre)* ^88, 89^. Lenti-sg*TS*102*/Cre* included vectors targeting 48 tumor suppressors, including all five of the “core” tumor suppressors and most of the tumor suppressors targeted in Lenti-sg*TS*15*/Cre* with two or three sgRNAs per gene in addition to five inert sgRNAs (102 sgRNA in total, Lenti-sg*TS*102*/Cre)* ^90^ (See **Table S1**).

We determined the alteration frequency of many putative tumor suppressor genes, including those targeted using our Lenti-sg*TS*15*/Cre* and Lenti-sg*TS*102*/Cre* pools, in oncogene-positive and oncogene-negative tumors from TCGA and GENIE ^60, 61^. Alterations in only 17 tumor suppressor genes were significantly enriched in oncogene-negative tumors in GENIE and most (12/17) were targeted by the Lenti-sg*TS*15*/Cre* and Lenti-sg*TS*102*/Cre* pools (**Table S1**).

We previously found that a small percent of lung tumors initiated with Lenti-sgRNA*/Cre* vectors in other lung cancer models contained multiple sgRNAs, consistent with the transduction of the initial cell with multiple Lenti-sgRNA*/Cre* vectors ^88, 89^. Thus, from tumor suppressor genes that were found to be mutated in the largest tumors and expansions of experiment in **Figure 1**, we selected 7 tumor suppressor genes that showed up in *Nf1^f/f^;TC, Pten^f/f^;TC, Trp53^f/f^;TC* mice in addition to 6 other tumor suppressor genes that showed significant effect in at least one of these three backgrounds. For Studies in **Figure 2** and **Figure S7**, we used higher titers of Lenti-sgRNA/*Cre* vectors to increase the potential of finding higher-order interactions that generate lung tumors. We found that simultaneous alterations of *Nf1*, *Rasa1*, and *Pten* was one of the most frequent co-occurring alterations in the largest tumors. Thus, we focused on studying these three tumor suppressor alterations using Lenti-sg*TripleTS8/Cre*, Lenti- sg*TripleTS6/Cre,* and Lenti-sg*Nf1-sgRasa1-sgPten/Cre* in the following figures.

The sgRNA sequences used in each experiment are summarized below. For a more detailed description see **Table S1**:

| Pool | Pool composition |
| --- | --- |
| Lenti-sg*TS15/Cre* | The exact pool used in<sup>88, 89</sup> |
| Lenti-sg*TS102/Cre* | The exact pool used in <sup>90</sup> |
| Lenti-sg*TS14/Cre* | Version 1 of sg*EP300*, sg*Kmt2c*, sg*Ncoa6*, sg*Rbm10*, sg*Neo*, sg*Nf1*, and sg*Pten*, and version 2 of sg*Arid1a*, sg*Cdkn2a*, sg*Dnmt3a*, sg*Kdm6a*, sg*Rb1*, sg*Tet2*, sg*Rasa1* from <sup>90</sup> |
| Lenti-sg*TS11/Cre* | Lenti-sg*TS14/Cre* pool excluding sg*Nf1*, sg*Rasa1*, and sg*Pten* |
| Lenti-sgTriple*TS8/Cre* | Version 1 of sg*Nf1*, sg*Neo*, and sg*NT*, and version 2 of sg*Rasa1*, sg*Pten*, and sg*Neo* from <sup>90</sup> |
| Lenti-sgTriple*TS6/Cre* | Version 1 of sg*Nf1*, sg*Neo*, and sg*NT*, and version 2 of sg*Rasa1*, sg*Pten*, and sg*Neo* from <sup>90</sup>. Vectors targeting only *Nf1* and only *Rasa1* were removed from this pool. |

### Lentiviral generation, barcoding, and packaging

The sgRNA sequences, cloning, and barcoding of Lenti-sg*RNA/Cre and* Lenti- TriplesgRNA*/Cre* vectors have been previously described ^88, 90, 91^. To generate lentivirus, Lenti- sg*RNA/Cre* vectors were individually co-transfected into 293T cells with pCMV-VSV-G (Addgene #8454) envelope plasmid and pCMV-dR8.2 dvpr (Addgene #8455) packaging plasmid using polyethylenimine. Supernatants were collected 36 and 48 hours after transfection, passed through a 0.45µm syringe filter (Millipore SLHP033RB) to remove cells and cell debris, concentrated by ultracentrifugation (25,000 g for 1.5 hours at 4°C) and resuspended in PBS overnight. Each virus was titered against a standard of known titer using LSL-YFP Mouse Embryonic Fibroblasts (MEFs) (a gift from Dr. Alejandro Sweet-Cordero/UCSF). All lentiviral vector aliquots were stored at -80°C and were thawed and pooled immediately prior to delivery to mice.

### Tumor barcode sequencing and analysis

For DNA extraction from single dissected tumors to generate libraries for Tuba-seq, targeted sequencing of selected oncogenes, and whole-exome sequencing, we used Qiagen AllPrep DNA/RNA Micro kit. For Tuba-seq on bulk lungs, genomic DNA was isolated from bulk tumor-bearing lung tissue from each mouse as previously described ^88^. Briefly, benchmark control cell lines were generated from LSL-YFP MEFs transduced by a barcoded Lenti- sgNT3*/Cre* vector (NT3: an inert sgRNA with a unique sgRNA identifying barcode (sgID) and a random barcode (BC)) and purified by sorting YFP^+^ cells using BD FACS Aria™ II Cell Sorter. Three cell lines (100,000 to 500,000 cells each) were added to each mouse lung sample before lysis to enable the calculation of the absolute number of neoplastic cells in each tumor from the number of sgID-BC reads. Following homogenization and overnight protease K digestion, genomic DNA was extracted from the lung lysates using standard phenol-chloroform and ethanol precipitation methods. Subsequently, Q5 High-Fidelity 2x Master Mix (New England Biolabs, M0494X) was used to amplify the sgID-BC region from 50 ng of DNA from dissected tumors or 32 μg of bulk lung genomic DNA. The unique dual-indexed primers used were Forward: AATGATACGGCGACCACCGAGATCTACAC- 8 nucleotides for i5 index- ACACTCTTTCCCTACACGACGCTCTTCCGATCT-6 to 9 random nucleotides for increasing the diversity-GCGCACGTCTGCCGCGCTG and Reverse: CAAGCAGAAGACGGCATACGAGAT-6 nucleotides for i7 index-GTGACTGGAGTTCAGACGTGTGCTCTTCCGATCT-9 to 6 random nucleotides for increasing the diversity-CAGGTTCTTGCGAACCTCAT. The PCR products were purified with Agencourt AMPure XP beads (Beckman Coulter, A63881) using a double size selection protocol. The concentration and quality of the purified libraries were determined using the Agilent High Sensitivity DNA kit (Agilent Technologies, 5067-4626) on the Agilent 2100 Bioanalyzer (Agilent Technologies, G2939BA). The libraries were pooled based on lung weights to ensure even reading depth, cleaned up again using AMPure XP beads, and sequenced (read length 2x150bp) on the Illumina HiSeq 2500 or NextSeq 500 platform (Admera Health Biopharma Services).

### Tuba-seq analysis of tumor barcode reads

The FASTQ files were parsed to identify the sgID and barcode (BC) for each read. Each read is expected to contain an 8-nucleotide sgID region followed by a 30-nucleotide barcode (BC) region (GCNNNNNTANNNNNGCNNNNNTANNNNNGC), and each of the 20 Ns represents random nucleotides. The sgID region identifies the putative tumor suppressor gene being targeted, for which we require a perfect match between the sequence in the forward read and one of the forward sgIDs with known sequences. Note that all sgID sequences differ from each other by at least three nucleotides. Therefore, the incorrect assignment of sgID due to PCR or sequencing error is extremely unlikely. All cells generated from the clonal expansion of an original cell transduced with a lentiviral vector carry the same BC sequence. To minimize the effects of sequencing errors on calling the BC, we require the forward and reverse reads to agree completely within the 30-nucleotide sequence to be further processed. In our pipeline, any tumor that is within a Hamming distance of two from a larger tumor is assigned as a “spurious tumor”, which likely results from sequencing or PCR errors and the tumor is removed from subsequent analysis. Reads with the same sgID and barcode are assigned to be the same tumor. The tumor size (number of neoplastic cells) is calculated by normalizing the number of reads to the three benchmarks “spike-in” cell lines added to each sample prior to lysis of the lung and DNA extraction step. The median sequencing depth was ∼ 1 read per 4.8 cells, and the minimum sequencing depth is ∼1 read per 16.5 cells. We have high statistical power in identifying tumors with more than 200 cells, which was used as the minimum cell number cutoff for calling tumors. A minimum cell number of 50 was used for calling expansions in **Figures S5** and **S6**). Minimizing the influence of GC amplification bias on tumor-size calling was done as previously described ^88^.

### Measures of tumor size and growth

We used several metrics of tumor number, burden and size (see **Supplemental Figure 4** in ^90^ for additional details on the calculation of these metric).

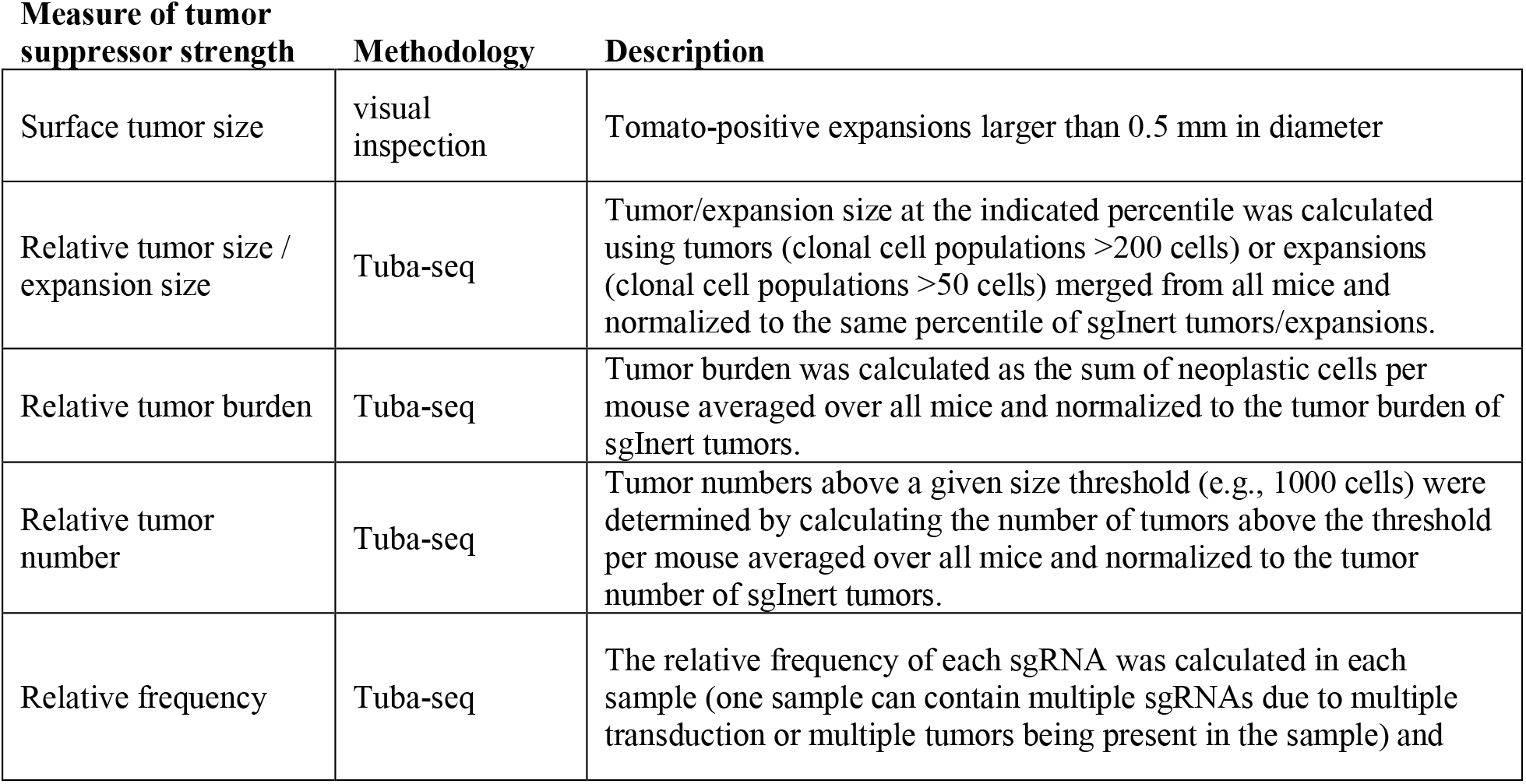
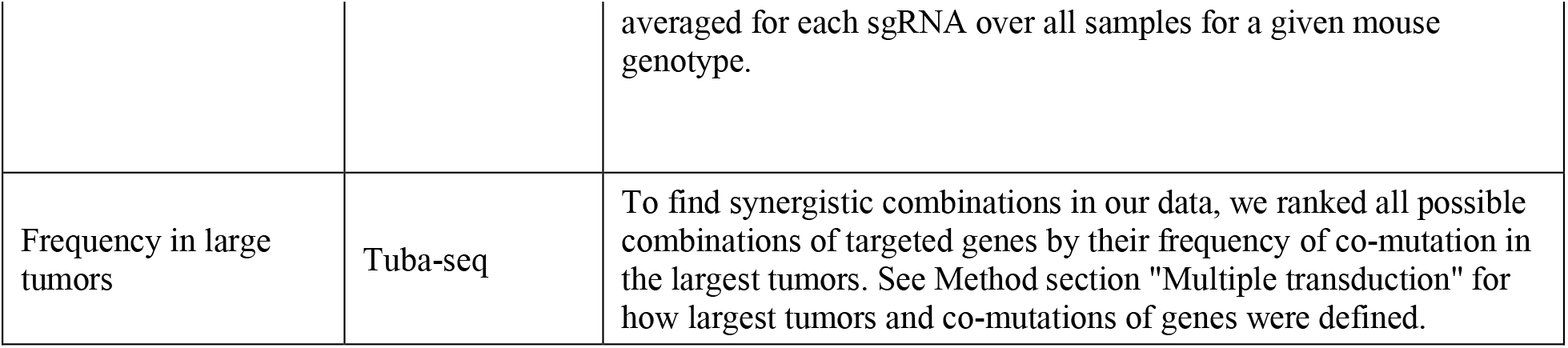

Tumor burden and tumor number are affected linearly by the titer of each Lenti-sgRNA/Cre vector in the pool. When applicable, we used data on the number of tumors from *KT* mice (which lack Cas9) to quantify the representation of each Lenti-sgRNA/Cre vectors in the lentiviral pool. Therefore, when calculating tumor burden and tumor number metrics, we normalized the metric to the effective titer based on data from *KT* mice to account for the viral titer differences among different Lenti-sgRNA/Cre vectors. Tumor/expansion size percentiles, tumor burden, and tumor number were normalized to the values of the same metric for tumors with inert sgRNAs, thus the expression “relative” is used.

For relative tumor/expansion size, relative tumor burden and relative tumor number, confidence intervals and p*-* values were calculated by a nested bootstrap resampling approach to account for variation in sizes of tumors of a given genotype both across and within mice. First, tumors of each mouse were grouped, and these groups (mice) were resampled. Second, all tumors of a given mouse resampling were bootstrapped on an individual basis (10,000 repetitions). For relative frequency, tumors were bootstrap resampled 10,000 times, and the distribution of inert sgRNA frequencies was used to calculate p-values for enrichment of all other sgRNAs. For “frequency in large tumors”, a permutation test was used to calculate p-values (see section **Multiple transduction** for details).

### Multiple transduction

A fraction of lung tumors initiated with Lenti-sg*RNA*/*Cre* vectors contained multiple barcoded Lenti-sgRNA/Cre vectors. If multiple barcodes (sgID-BCs) have unexpectedly similar read counts (as shown in the example plots below), we suspect transduction of the initial cell with multiple Lenti-sg*RNA/Cre* vectors.

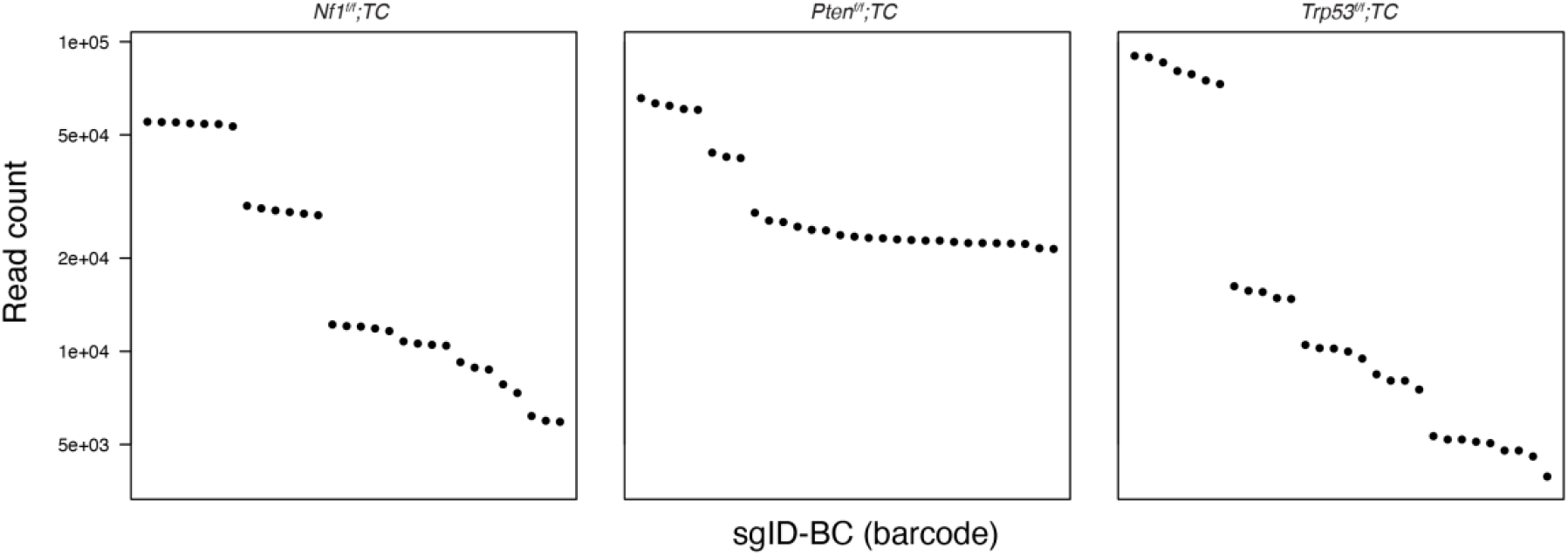
Example plots indicating strong evidence of transduction with multiple barcoded lentiviral vectors in the largest tumors in each genotype. 30 sgID-BCs with the highest read counts from representative mouse samples are shown. Indicated genotypes of mice were transduced with Lenti-sgTS14/*Cre* pool. Each dot represents a sgID-BC, the y-axis shows read count, and the sgID-BCs are sorted on the x-axis by decreasing read counts. Groups of barcodes sgID-BCs that have very similar read counts likely represents a single clonal tumor initiated from a cell transduced with multiple barcoded Lenti-sgRNA/*Cre* vectors.

To capitalize on these multiple transductions as a way to find higher-order interactions between tumor suppressor genes, we developed a method to identify the combinations of sgRNA that appear to cooperate as potent drivers of tumor growth. Accurate identification of coinfected tumors and grouping of barcodes without over grouping was not a trivial task. We developed methods to identify tumors with likely multiple transductions (*i.e*., those tumors with complex genotypes with multiple tumor suppressor genes inactivated). For each sgID-BC, we listed all other sgID-BCs from the same sample with read counts within 10% as possible multiple transduction events. A tumor with multiple transductions can be most easily identified among the largest tumors in each mouse as smaller tumors of similar sizes are too abundant. Multiple transductions that lead to synergistic combinatorial tumor suppressor alterations would confer a growth advantage. Thus, synergistic combinatorial alterations of tumor suppressor genes would be expected to be overrepresented among the largest tumors.

To have a dataset with a higher signal-to-noise ratio, we analyzed the largest tumors that were co-infected with up to 6 Lenti-sgRNA/*Cre* vectors. With this method, for each tumor, we assembled a list of genes that were possibly co-mutated. We then ranked all possible combinations of genes by their frequency in the largest tumors (**Figure 2f-g** and **S6c-h**).

An inherent problem with this analysis is that the genotypes that increase tumor growth will be overrepresented amongst the largest tumors even without multiple transductions and specific synergistic interactions. To account for the different number of tumors with different sgIDs, we performed a permutation test, where we control for the number of tumors of each genotype but randomize the sizes of tumors by randomly matching the genotypes with tumor sizes (10,000 repetitions). Synergistic tumor suppressor combinations will occur at significantly higher than expected frequencies based on this permutation test (**Figure 2f-g** and **S6c-h**).

Reassuringly, while our analysis resulted in significant enrichment of complex genotypes based on the permutation test, a control analysis performed on smaller tumors within the same mice with high noise to signal ratio resulted in a loss of statistical significance, this shows that our permutation test controls for the bias of different frequency of sgIDs among the tumors.

### Fitness landscape and adaptive paths

To investigate the possible adaptive steps that can lead to the complex genotype of coincident inactivation of *Nf1*, *Rasa1*, and *Pten*, we first measured the fitness of all possible combinations of *Nf1*, *Rasa1*, and *Pten* mutations (**Figure 3f** and **S10g**). Relative (Malthusian) fitness was calculated based on the number of individuals (cells) at the end (N1) and the beginning of (N0) of a time period ^92^. For each genotype, the overall sum of neoplastic cells at the end of the experiment (N1) was calculated as the sum of cells from all tumors in each mouse. As we use *KT* mice (which lack Cas9 and all sgRNAs have no effect) to approximate the effective titer of our virus pool (see section Measures of tumor size and growth), the initial number of cells transduced (N0) was calculated from the number of tumors generated in control *KT* mice. Next, the relative fitness for genotype A compared to wild type (wt) was calculated as:

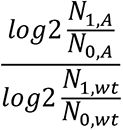

Fitness values relative to wild type are displayed as nodes on the adaptive landscape (**Figure 3f** and **S10g**), where genotypes one mutation away from each other are connected by arrows that represent mutations. In the case of the *Nf1;Rasa1;Pten* triple mutant state, six adaptive paths can lead from wild type to that triple mutant genotype (**Figure 3f** and **S10g**). Arrows are shown if the mutation increases the fitness. In **figure 3f** and **S10g** all arrows are shown since all mutations increase fitness.

Next, we set out to approximate the relative probabilities of different adaptive paths leading from wild type to the triple mutant genotype with a simple population genetic model. In the model, cell populations start from the wild-type genotype, and they can acquire any of the three mutations present in the triple genotype. In the population of cells, a mutation can arise and then change in frequency until one of two outcomes happens: (i) the frequency of the mutation drops to zero, and the mutation is lost from the population or (ii) the frequency of the mutation reaches 1, when it is present in all the cells and hence is fixed in the population. When a mutation fixes in a population, we consider the genotype of the population to change and that constitutes a “step” on the fitness landscape. We assume a “strong selection weak mutation” regime, where there is no more than one mutation simultaneously present with a frequency less than 1. We also assume that mutations appear randomly and with equal probabilities. Mutations can appear and get lost multiple times in a population, and as long as populations have at least one mutation that increases fitness, one of those mutations will fix in the population eventually.

With the model we are estimating the probability of each adaptive step given that the population starts from the wild-type state. Therefore, the probability of each adaptive step will be influenced by the probabilities of previous step(s) and the sum of probabilities of adaptive steps originating from a given genotype must equal the sum of probabilities of all adaptive steps terminating in the given genotype. If there are multiple adaptive steps originating from the same genotype, they will have probabilities proportional to the fixation probabilities of their respective mutations. The fixation probability of a mutation is proportional to its selective advantage 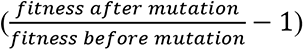^93^. As an example, if there are two adaptive steps originating from a genotype with fitness 1.00, one terminating in a genotype with fitness 1.1, the other in a genotype with fitness 1.2, then they have 10% and 20% selective advantage, respectively. Therefore, one adaptive step will happen half as likely as the other, as the selective advantages and therefore the relative fixation probabilities are in a ratio of 1:2.

### Targeted sequencing of oncogenic loci for potential spontaneous oncogenic mutation

To determine whether the tumors that develop contained spontaneous oncogenic mutations, we performed Sanger Sequencing and Illumina sequencing (HiSeq 2500 platform; read length 2x150 bp, Admera Health Biopharma Services) on select regions of *Kras, Egfr, Braf*, and *Nras* (the 4 frequently mutated oncogenes in lung adenocarcinoma).

PCR products were obtained through amplification with primers listed below on DNA extracted from dissected tumors (**Table S2**) and cleaned up using ExoSAP (ThermoFisher Scientific, Cat# 78-201) treatment before Sanger.

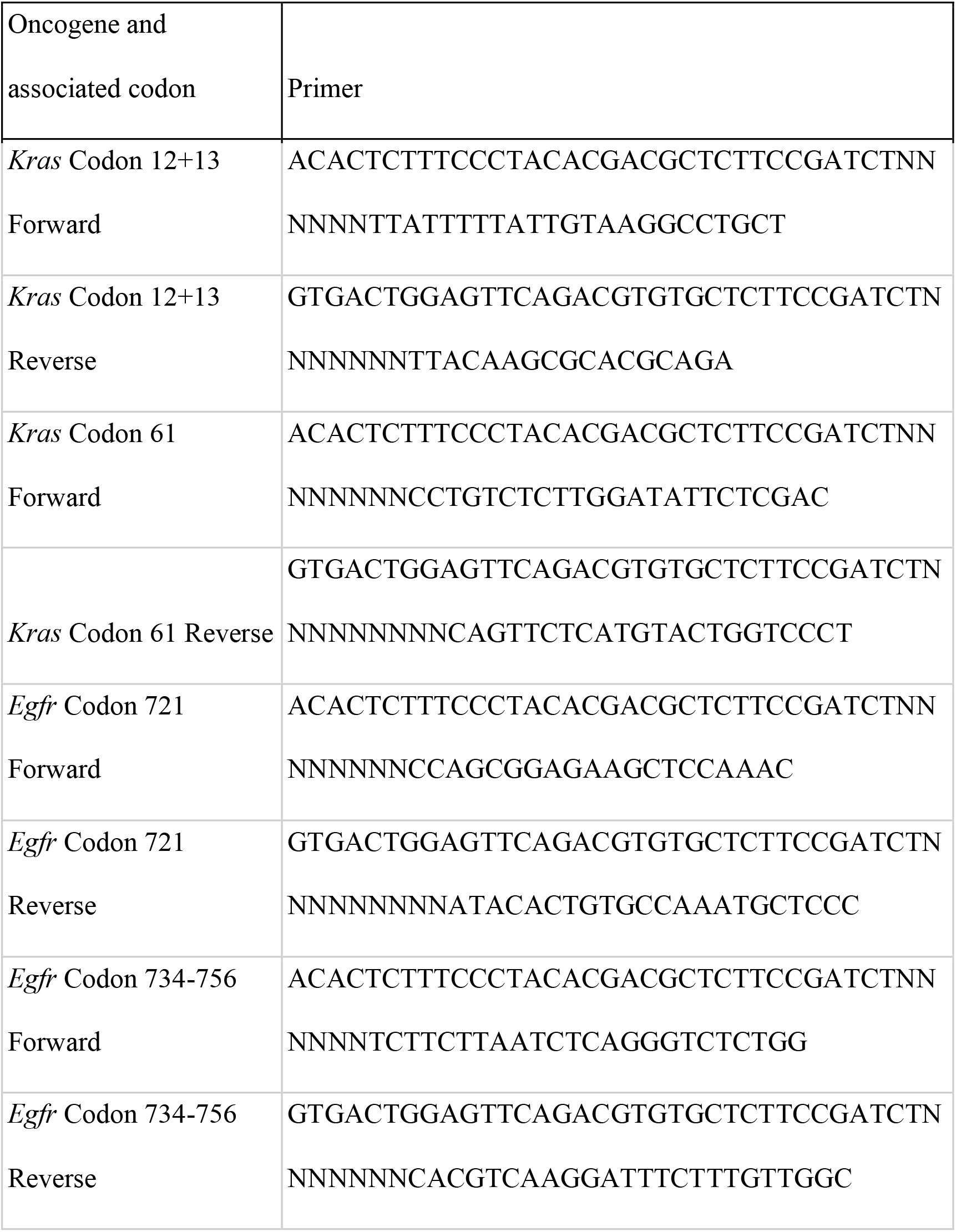
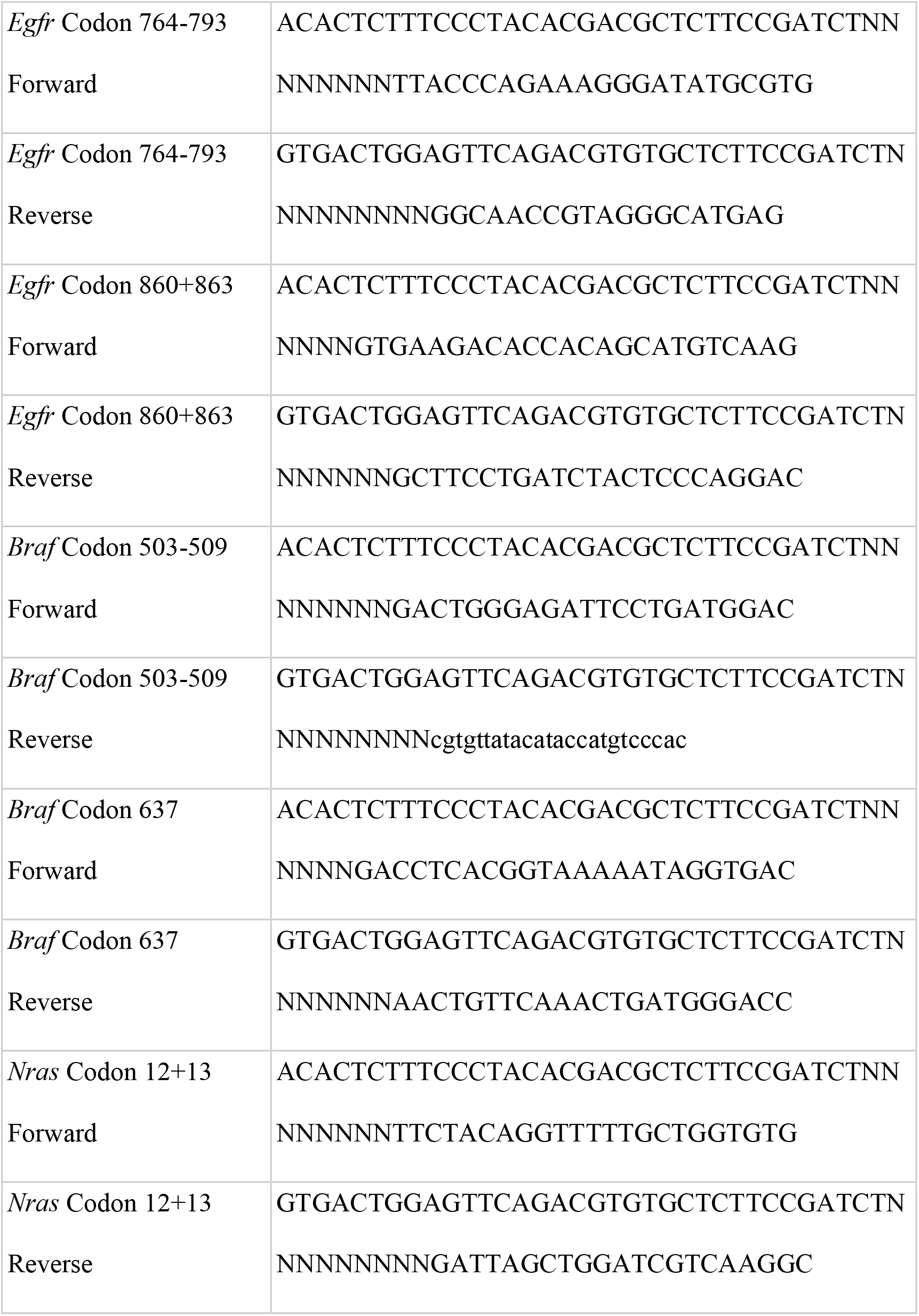
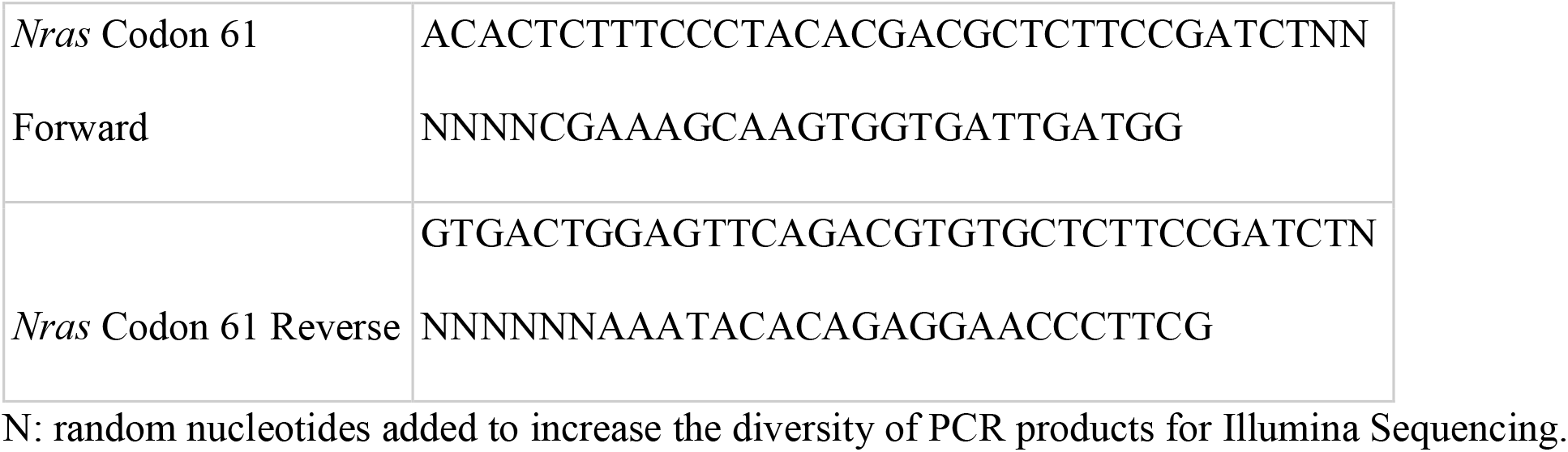

Illumina sequencing was performed on pools of amplicons. The libraries were pooled based on band intensity to ensure even read depth and cleaned up using Sera-Mag Select beads (Thermo Fisher Scientific, Cat# 09-928-107) before undergoing a second round of PCR to attach the sequencing adaptors needed for the HiSeq platform. Second round PCR products were then purified with Sera-Mag Select beads before sequencing.

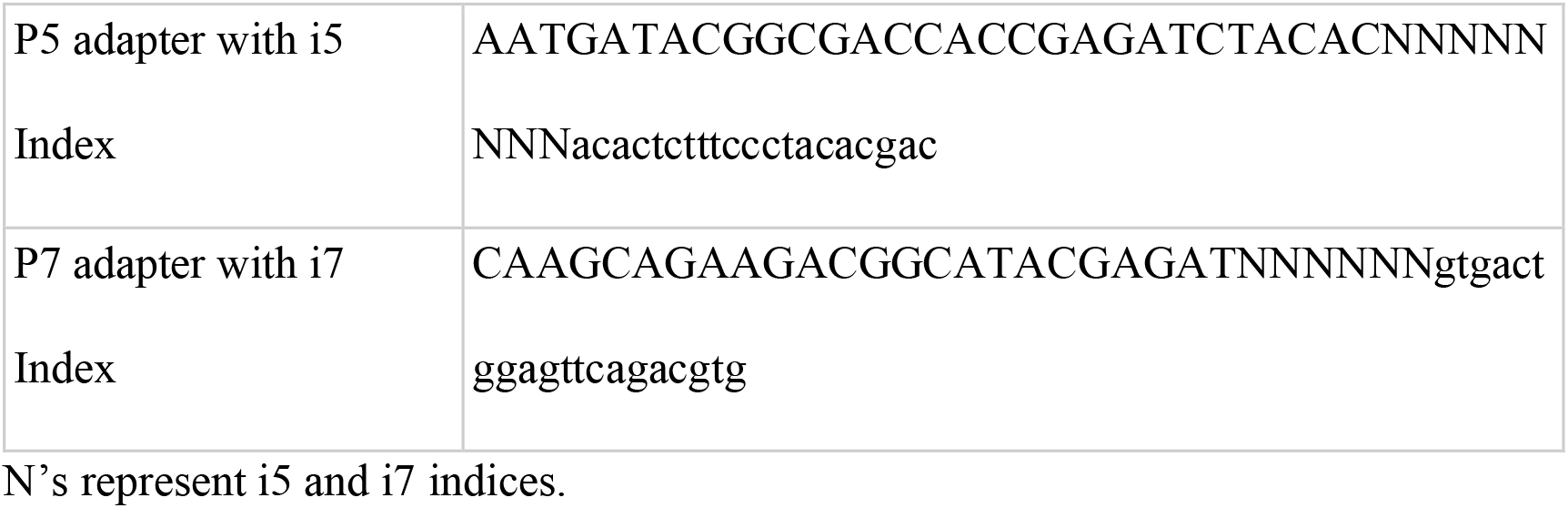

### Analysis of targeted DNA-sequencing of *Kras*, *Egfr*, *Braf*, and *Nras* oncogenic loci

Sequenced reads were analyzed using Genome Analysis Toolkit (GATK, Broad Institute ^94^). “Somatic short variant discovery” best practices pipeline for tumor samples similarly as for whole exome sequencing (see below). However, for targeted sequencing, identification of duplicate reads (Picard MarkDuplicates algorithm) was omitted as that would result in the loss of reads with matching start and end position, which is normal in targeted sequencing and is not a sign of duplicate artifacts. A mean coverage of 6665-7584 reads was achieved for all samples with 90% of regions having a coverage over 275 reads in all samples. Variant calls made and filtered by GATK Mutect2 function were annotated with Ensembl Variant Effect Predictor ^95^. Pick-allele-gene option was used to filter results on the most relevant transcript for each variation. We filtered the results for the known oncogenic codons listed above and variants with a minimum of 5% allele frequency.

### Whole exome sequencing

DNA was extracted from 4 individual tumors from *TC* mice transduced with Lenti-sg*Nf1*- sg*Rasa1*-sg*Pten*, three months after tumor initiation, using Qiagen AllPrep DNA/RNA Micro kit. Whole-exome sequencing library preparation was performed by Admera Health using SureSelect XT Mouse All Exon Kit (Agilent).

Sequenced reads on autosomes were analyzed using Genome Analysis Toolkit (GATK, Broad Institute ^94^) “Somatic short variant discovery” best practices pipeline for tumor samples. Mean coverage of 50-72 reads was achieved for all samples, with 90% of regions having coverage over 20 reads in all samples. Variant calls made and filtered by GATK Mutect2 function and were annotated with Ensembl Variant Effect Predictor (VEP ^95^). The pick-allele-gene option was used to filter results on the most relevant transcript for each variant. The same exact variants appearing in multiple tumor samples were flagged as germline variant and were removed. We filtered the results for protein-coding variation, variants with a minimum of 5% allele frequency, and removed variations in the olfactory OLFR gene family that are likely germline variations.

### Analysis of insertion and deletions

Indel analysis was performed to confirm that insertion and deletions (indels) were generated at the targeted loci as follows: gDNA was isolated from at oncogene-negative mouse cell lines or FACS-sorted Tomato^positive^ cancer cells using either the AllPrep DNA/RNA(Qiagen) or the DNeasy Blood and Tissue Kit. PCR primers were designed to amplify sgRNA-targeted loci, resulting in 500 to 1000 bp amplicons specific to each locus. Amplicons were purified using PCR purification kit (Qiagen) and sequenced by Sanger sequencing. Cutting efficiency was determined by ICE analysis (https://ice.synthego.com/<u>#/</u>)

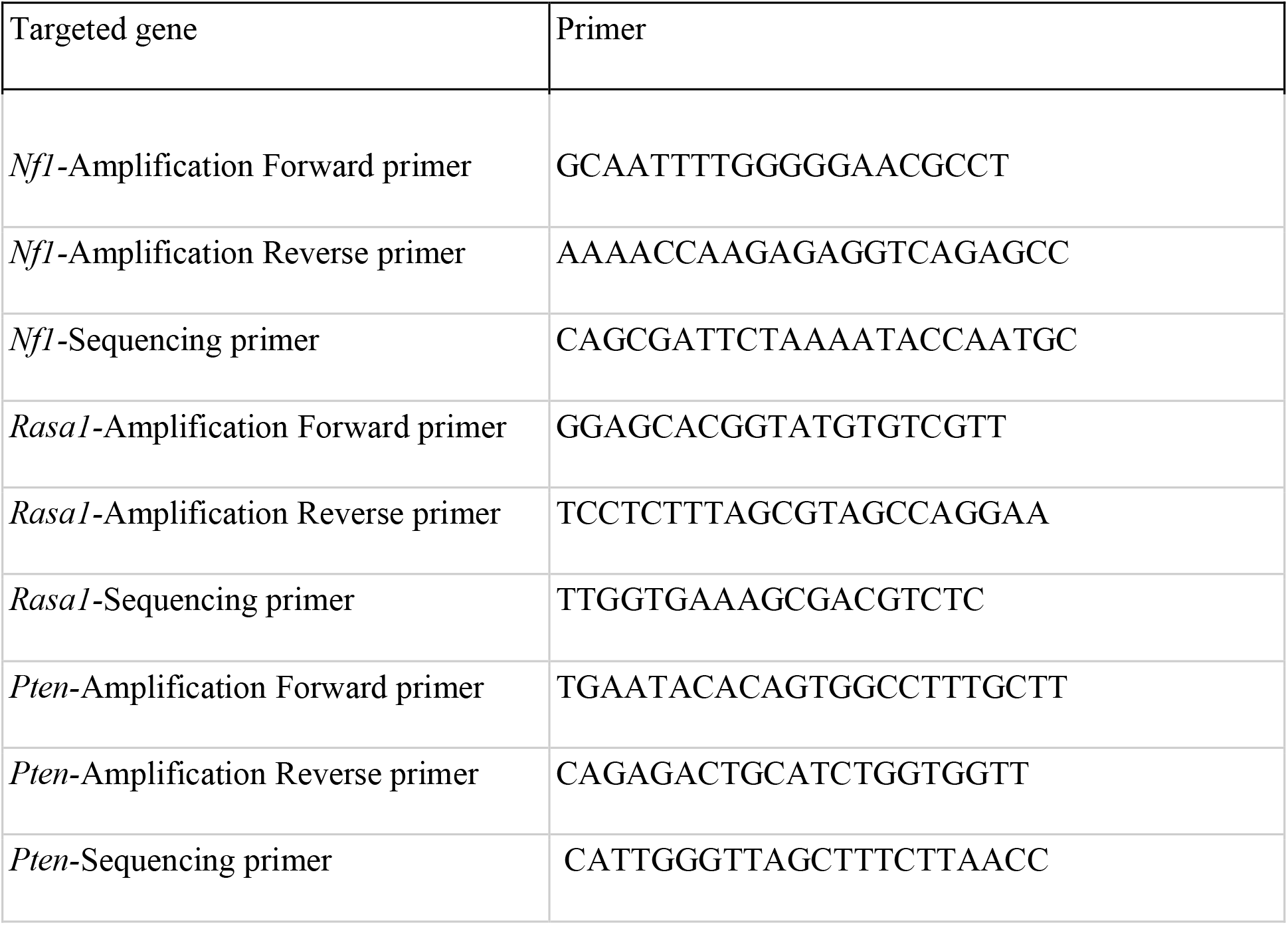

### Histology and immunohistochemistry

Lung lobes were inflated with 4% formalin and fixed for 24 hours, stored in 70% ethanol, paraffin-embedded, and sectioned. 4 μm thick sections were used for Hematoxylin and Eosin (H&E) staining and immunohistochemistry (IHC).

Primary antibodies used for IHC were anti-RFP (Rockland, 600-401-379), anti- TTF1(Abcam, ab76013), anti-UCHL1(Sigma, HPA005993), anti-TP63 (Cell Signaling Technology, 13109), anti-phospho-S6 (Cell Signaling Technology, 4858), anti-PTEN (Cell Signaling Technology, 9559), anti-phospho-ERK (Cell Signaling Technology, 4370), anti-phospho-AKT (Thermo Fisher Scientific, 44-621G), and anti-HMHGA2 (Biocheck, 59170AP). IHC was performed using Avidin/Biotin Blocking Kit (Vector Laboratories, SP-2001), Avidin-Biotin Complex kit (Vector Laboratories, PK-4001), and DAB Peroxidase Substrate Kit (Vector Laboratories, SK-4100) following standard protocols.

Images of the H&E-stained slides were analyzed with ImageJ. Tumor areas were converted from pixels to mm^2^ via a ruler. To quantify the positivity of phospho-ERK and phospho-AKT stained slides, H-scores were calculated using Qupath. The H-score is determined by adding the results of multiplication of the percentage of cells with staining intensity ordinal value (scored from 0 for “no signal” to 3 for “strong signal”) with possible values ranging from 0 to 300 ^96^. To normalize potential variations between different rounds of immunohistochemistry, one patient sample was included and stained for both pERK and pAKT in all rounds of staining as a control.

### Immunoblotting

3 × 10^5^ cells were seeded into 6-well plates and allowed to adhere overnight in regular growth media and cultured in the presence or absence of 10 µM of Capivasertib, RMC-4550, or a combination of both drugs. After 24 hours, the protein was extracted using RIPA lysis buffer (Thermo Fisher Scientific, 89900) and proteinase/phosphatase inhibitor cocktail (Thermo Fisher Scientific, 78442). Protein concentration was measured using BCA protein assay kit (Thermo Fisher Scientific, 23250). Proteins (30 µg from each sample) were separated by SDS-PAGE and immunoblotted and transferred to polyvinyl difluoride (PVDF) membranes (BioRad, 162-0177) according to standard protocols. Membranes were immunoblotted with antibodies against phosphor-ERK (Cell Signaling Technology, 4370), ERK (Cell Signaling Technology, 9102), phosphor-AKT (Thermo Fisher Scientific, 44-621G), AKT (Cell Signaling Technology, 4691), phospho-S6 (Cell Signaling Technology, 4858), S6 (Cell Signaling Technology, 2217), anti- RASA1 (Abcam, ab2922), anti-PTEN (Cell Signaling Technology, 9559), and HSP90 (BD Bioscience, 610418). Immunoblots were developed using Supersignal® West Dura Extended Duration Chemiluminescent Substrate (Thermo Fisher Scientific, 37071). Initially, the membranes were immunoblotted against non-phosphorylated targets, and after stripping these antibodies using Western Blot Stripping Buffer (Thermo Fisher Scientific, 46430), they were immunoblotted against phosphorylated antibodies. Developing the signal was done using Dura Extended Duration Chemiluminescent Substrate (Thermo Fisher Scientific, 37071). All immunoblots were performed at least three times independently.

### Cell Lines and Reagents

Mouse oncogene-negative cell lines were generated from tumors initiated in *Trp53^flox/flox^;TC BL6* mice four months after transduction with Lenti-sg*Nf1*-sg*Rasa1*-sg*Pten*/*Cre.* After dissociation of tumors (described below), cells were cultured in DMEM supplemented with 10% FBS, 1% penicillin/streptomycin (Gibco), and 0.1% Amphotericin (Life Technologies).

HC494 and MMW389T2 (*Kras^G12D^* and *Trp53* mutant) lung adenocarcinoma cells were previously generated in the Winslow Lab. Human oncogene-negative cell lines (NCI-H1838, NCI-H1623) and oncogene-positive cell lines (A549, H2009, NCI-H2009, SW1573, HOP62, NCI-H358, NCI-H1792) were purchased from ATCC and cultured in RPMI supplemented with 5%FBS, 1% penicillin/streptomycin (Gibco), and 0.1% Amphotericin (Life Technologies). We performed mycoplasma testing using MycoAlert™ Mycoplasma Detection Kit (Lonza). Cell were maintained at 37°C in a humidified incubator at 5% CO2. Mutations in components of RAS and PI3K pathways of NCI-H1838, NCI-H1623 (based on **Table S6**) are indicated in the table below (extracted from DepMap):

| Cell line | Pathway | Gene | Protein Change |
| --- | --- | --- | --- |
| NCI-H1838 | RAS | NF1 | p.N184fs |
| NCI-H1838 | RAS | IQGAP2 | p.P780L |
| NCI-H1623 | RAS | FGFR2 | p.A355S |
| NCI-H1623 | RAS | ERF | p.G255C |
| NCI-H1623 | RAS | RASA1 | p.A47fs |

### Clonogenic, apoptosis, and proliferation assays

For clonogenic assays, mouse cells were seeded in triplicate into 24-well plates (4000 cells per well) and allowed to adhere overnight in regular growth media. Cells were then cultured in the absence or presence of the drug as indicated on each figure panel in complete media for 4 days. Growth media with or without drugs was replaced every 2 days. The remaining cells were stained with 0.5% crystal violet in 20% methanol and photographed using a digital scanner. Relative growth was quantified by densitometry after extracting crystal violet from the stained cells using 100% methanol ^97^.

Clonogenic assay of human oncogene-negative lung adenocarcinoma cell lines were done in spheroids ^98^. 400-5000 cells/well were seeded in round bottom ultra-low attachment 96-well plates (Corning) in growth media and incubated for 72 hours at 37°C in 5% CO2. Spheroid formation was confirmed visually, and spheroids were treated in triplicate with dilutions of RMC-4550 and capivasertib in complete growth media. Following drug exposure for five days, cell viability in spheroids was determined using the CellTiter-Glo 3D assay kit (Promega), following the manufacturer’s instructions. Luminescence was read in a Plate Reader. Assay data was normalized to DMSO values.

Drug synergism was analyzed using SynergyFinder (https://synergyfinder.fimm.fi) web application ^99^. The degree of combination synergy, or antagonism, was quantified by comparing the observed drug combination response against the expected response, calculated using Loewe’s model that assumes no interaction between drugs ^100^.

For apoptosis and proliferation assays, 3 × 10^5^ cells were seeded into 6-well plates, and allowed to adhere overnight in regular growth media, and cultured in the presence or absence of 10 µM of Capivasertib, RMC-4550, or a combination of both drugs. After 24 hours, apoptosis and cell proliferation were determined through staining with Fixable Viability Dye eFluor™ 450 (Thermo Fisher Scientific, 65-0863-14), cleaved caspase 3 Antibody (Cell Signaling Technology, 9669), and Click-iT™ EdU Alexa Fluor™ 647 Flow Cytometry Assay Kit (Thermo Fisher Scientific, C-10424) according to the manufacturer’s instructions. Data were acquired using a BD LSR II Flow Cytometer. All experiments were performed independently two times on 3 different cell lines.

### *In vivo* drug response studies

For drug efficacy studies in autochthonous mouse models, *TC* mice (8-12 weeks old) were divided into 4 groups randomly 3.5 months after tumour initiation. They received the vehicle, capivasertib (100 mg/kg, MedChemExpress), RMC-4550 (30 mg/kg, MedChemExpress), or a combination of both dissolved in 10% DMSO, 40% PEG, 5% Tween 80, and 45% PBS through a gavage needle. Mice were treated daily with drugs for eight days, and the treatment was stopped for two days for recovery, and it continued for two more days before the tissue harvest. The last two doses of combination therapy were half of the initial doses.

### Tumor dissociation, cell sorting, and RNA-sequencing

Primary tumors were dissociated using collagenase IV, dispase, and trypsin at 37 °C for 30 min. After dissociation, the samples remained continually on ice, were in contact with ice-cold solutions, and were in the presence of 2 mM EDTA and 1 U/ml DNase to prevent aggregation. Cells were stained with antibodies to CD45 (BioLegened, 103112), CD31 (BioLegend, 303116), F4/80 (BioLegend, 123116), and Ter119 (BioLegend, 116212) to exclude hematopoietic and endothelial cells (lineage-positive (Lin^+^) cells). DAPI was used to exclude dead cells. FACS Aria sorters (BD Biosciences) were used for cell sorting.

RNA was purified using RNA/DNA All Prep kit (Qiagen, 80284). RNA quality of each tumor sample was assessed using the RNA6000 PicoAssay for the Agilent 2100 Bioanalyzer as per the manufacturer’s recommendation. 4.4 ng total RNA per sample was used for cDNA synthesis and library preparation using Trio RNA-Seq, Mouse rRNA kit (Tecan, 0507-32), according to the manufacturer’s instructions. The purified cDNA library products were evaluated using the Agilent bioanalyzer and sequenced on NextSeq High Output 1x75 (Admera Health Biopharma Services).

### Analysis of mouse model-derived RNA-seq datasets

Paired-end RNA-seq reads were aligned to the mm10 mouse genome using STAR (v2.6.1d) 2-pass mapping and estimates of transcript abundance were obtained using RSEM (v1.2.30) ^101, 102^. The differentially expressed genes between different tumor genotypes and treatment groups were called by DESeq2 using transcript abundance estimates via tximport ^103, 104^. The DESeq2-calculated fold changes were used to generate ranked gene lists for input into GSEA ^105^.

The upregulated genes with absolute log2 fold change greater than 1 and a false discovery rate less than 0.05 in the comparison of *Nf1*, *Rasa1*, and *Pten* mutant oncogene-negative tumors with Kras^G12D^-driven tumors (*KTC*+sg*Inert* and *KTC*+sg*Pten)* were compiled into a signature reflecting the oncogene-negative adenocarcinoma state. This gene signature was utilized in the analysis of human oncogene-positive and oncogene-negative tumors. Scaled estimates of transcript abundance for TCGA LUAD samples were obtained from the GDC data portal (gdc-portal.nci.nih.gov). Each expression profile was then scored on the basis of the mouse-derived gene signature using single-sample GSEA within the Gene Set Variation Analysis (GSVA) package ^106^.

### Data availability

Tuba-seq barcode sequencing and RNA-seq data have been deposited in NCBI’s Gene Expression Omnibus (https://www.ncbi.nlm.nih.gov/geo/) and are accessible through GEO Series accession number GSE174393. Whole exome sequencing data generated in our study are publicly available in SRA-NCBI (www.ncbi.nlm.nih.gov/sra), under BioProject accession number PRJNA769722.

## Acknowledgments

The results in Figures 1 and 4 are in part based upon data generated by the TCGA Research Network (https://www.cancer.gov/tcga) and Genomics Evidence Neoplasia Information Exchange (GENIE).

## REFERENCES

1. Barta, J.A., Powell, C.A. & Wisnivesky, J.P. Global Epidemiology of Lung Cancer. Ann Glob Health 85 (2019).

2. Devarakonda, S., Morgensztern, D. & Govindan, R. Genomic alterations in lung adenocarcinoma. Lancet Oncol 16, e342–351 (2015).

3. McDermott, U., Downing, J.R. & Stratton, M.R. Genomics and the continuum of cancer care. N Engl J Med 364, 340–350 (2011).

4. Cancer Genome Atlas Research, N. Comprehensive molecular profiling of lung adenocarcinoma. Nature 511, 543-550 (2014).

5. Carrot-Zhang, J. et al. Whole-genome characterization of lung adenocarcinomas lacking the RTK/RAS/RAF pathway. Cell Rep 34, 108707 (2021).

6. Campbell, J.D. et al. Distinct patterns of somatic genome alterations in lung adenocarcinomas and squamous cell carcinomas. Nat Genet 48, 607–616 (2016).

7. Lawrence, M.S. et al. Discovery and saturation analysis of cancer genes across 21 tumour types. Nature 505, 495–501 (2014).

8. Vaishnavi, A. et al. Oncogenic and drug-sensitive NTRK1 rearrangements in lung cancer. Nat Med 19, 1469–1472 (2013).

9. Jonna, S. et al. Detection of NRG1 Gene Fusions in Solid Tumors. Clin Cancer Res 25, 4966–4972 (2019).

10. Soda, M. et al. Identification of the transforming EML4-ALK fusion gene in non-small- cell lung cancer. Nature 448, 561–566 (2007).

11. Takeuchi, K., et al. RET, ROS1 and ALK fusions in lung cancer. Nat Med 18, 378–381 (2012).

12. Izumi, H. et al. The CLIP1-LTK fusion is an oncogenic driver in non-small-cell lung cancer. Nature (2021).

13. Vogelstein, B. et al. Cancer genome landscapes. Science 339, 1546–1558 (2013).

14. Yaffe, M.B. The scientific drunk and the lamppost: massive sequencing efforts in cancer discovery and treatment. Sci Signal 6, pe13 (2013).

15. Sanchez-Vega, F. et al. Oncogenic Signaling Pathways in The Cancer Genome Atlas. Cell 173, 321–337 e310 (2018).

16. Krogan, N.J., Lippman, S., Agard, D.A., Ashworth, A. & Ideker, T. The cancer cell map initiative: defining the hallmark networks of cancer. Mol Cell 58, 690–698 (2015).

17. George, J. et al. Comprehensive genomic profiles of small cell lung cancer. Nature 524, 47–53 (2015).

18. Gouyer, V. et al. Mechanism of retinoblastoma gene inactivation in the spectrum of neuroendocrine lung tumors. Am J Respir Cell Mol Biol 18, 188–196 (1998).

19. Sekido, Y., Fong, K.M. & Minna, J.D. Molecular genetics of lung cancer. Annu Rev Med 54, 73–87 (2003).

20. Meuwissen, R. et al. Induction of small cell lung cancer by somatic inactivation of both Trp53 and Rb1 in a conditional mouse model. Cancer Cell 4, 181–189 (2003).

21. Govindan, R. et al. Genomic landscape of non-small cell lung cancer in smokers and never-smokers. Cell 150, 1121–1134 (2012).

22. Soria, J.C. et al. Lack of PTEN expression in non-small cell lung cancer could be related to promoter methylation. Clin Cancer Res 8, 1178–1184 (2002).

23. Kazanets, A., Shorstova, T., Hilmi, K., Marques, M. & Witcher, M. Epigenetic silencing of tumor suppressor genes: Paradigms, puzzles, and potential. Biochim Biophys Acta 1865, 275–288 (2016).

24. Ding, L. et al. Somatic mutations affect key pathways in lung adenocarcinoma. Nature 455, 1069–1075 (2008).

25. Lee, J.S., Grisham, J.W. & Thorgeirsson, S.S. Comparative functional genomics for identifying models of human cancer. Carcinogenesis 26, 1013–1020 (2005).

26. Hutter, C. & Zenklusen, J.C. The Cancer Genome Atlas: Creating Lasting Value beyond Its Data. Cell 173, 283–285 (2018).

27. Consortium, A.P.G. AACR Project GENIE: Powering Precision Medicine through an International Consortium. Cancer Discov 7, 818–831 (2017).

28. Jorge, S.E., Kobayashi, S.S. & Costa, D.B. Epidermal growth factor receptor (EGFR) mutations in lung cancer: preclinical and clinical data. Braz J Med Biol Res 47, 929–939 (2014).

29. Skoulidis, F. & Heymach, J.V. Co-occurring genomic alterations in non-small-cell lung cancer biology and therapy. Nat Rev Cancer 19, 495–509 (2019).

30. Saito, M. et al. Gene aberrations for precision medicine against lung adenocarcinoma. Cancer Sci 107, 713–720 (2016).

31. Rogers, Z.N. et al. A quantitative and multiplexed approach to uncover the fitness landscape of tumor suppression in vivo. Nat Methods 14, 737–742 (2017).

32. Rogers, Z.N. et al. Mapping the in vivo fitness landscape of lung adenocarcinoma tumor suppression in mice. Nat Genet 50, 483–486 (2018).

33. Winters, I.P. et al. Multiplexed in vivo homology-directed repair and tumor barcoding enables parallel quantification of Kras variant oncogenicity. Nat Commun 8, 2053 (2017).

34. Winters, I.P., Murray, C.W. & Winslow, M.M. Towards quantitative and multiplexed in vivo functional cancer genomics. Nat Rev Genet 19, 741–755 (2018).

35. Cai, H., et al. A functional taxonomy of tumor suppression in oncogenic KRAS-driven lung cancer. Under consideration (2021).

36. Madisen, L. et al. A robust and high-throughput Cre reporting and characterization system for the whole mouse brain. Nat Neurosci 13, 133–140 (2010).

37. Chiou, S.H. et al. Pancreatic cancer modeling using retrograde viral vector delivery and in vivo CRISPR/Cas9-mediated somatic genome editing. Genes Dev 29, 1576–1585 (2015).

38. Lynch, T.J. et al. Activating mutations in the epidermal growth factor receptor underlying responsiveness of non-small-cell lung cancer to gefitinib. N Engl J Med 350, 2129–2139 (2004).

39. Ohashi, K. et al. Characteristics of lung cancers harboring NRAS mutations. Clin Cancer Res 19, 2584–2591 (2013).

40. Lin, Q. et al. The association between BRAF mutation class and clinical features in BRAF-mutant Chinese non-small cell lung cancer patients. J Transl Med 17, 298 (2019).

41. Jackson, E.L. et al. Analysis of lung tumor initiation and progression using conditional expression of oncogenic K-ras. Genes Dev 15, 3243–3248 (2001).

42. Paez, J.G. et al. EGFR mutations in lung cancer: correlation with clinical response to gefitinib therapy. Science 304, 1497–1500 (2004).

43. Politi, K. et al. Lung adenocarcinomas induced in mice by mutant EGF receptors found in human lung cancers respond to a tyrosine kinase inhibitor or to down-regulation of the receptors. Genes Dev 20, 1496–1510 (2006).

44. Li, D. et al. Bronchial and peripheral murine lung carcinomas induced by T790M-L858R mutant EGFR respond to HKI-272 and rapamycin combination therapy. Cancer Cell 12, 81–93 (2007).

45. van Veen, J.E. et al. Mutationally-activated PI3’-kinase-alpha promotes de-differentiation of lung tumors initiated by the BRAF(V600E) oncoprotein kinase. Elife 8 (2019).

46. Dankort, D. et al. A new mouse model to explore the initiation, progression, and therapy of BRAFV600E-induced lung tumors. Genes Dev 21, 379–384 (2007).

47. McFadden, D.G. et al. Mutational landscape of EGFR-, MYC-, and Kras-driven genetically engineered mouse models of lung adenocarcinoma. Proc Natl Acad Sci U S A 113, E6409–E6417 (2016).

48. Weinreich, D.M., Delaney, N.F., Depristo, M.A. & Hartl, D.L. Darwinian evolution can follow only very few mutational paths to fitter proteins. Science 312, 111–114 (2006).

49. Winslow, M.M. et al. Suppression of lung adenocarcinoma progression by Nkx2-1. Nature 473, 101–104 (2011).

50. Sweet-Cordero, A. et al. An oncogenic KRAS2 expression signature identified by cross- species gene-expression analysis. Nat Genet 37, 48–55 (2005).

51. Agarwal, A. et al. The AKT/I kappa B kinase pathway promotes angiogenic/metastatic gene expression in colorectal cancer by activating nuclear factor-kappa B and beta- catenin. Oncogene 24, 1021–1031 (2005).

52. Yang, S.R. et al. Comprehensive Genomic Profiling of Malignant Effusions in Patients with Metastatic Lung Adenocarcinoma. J Mol Diagn 20, 184–194 (2018).

53. Maertens, O. & Cichowski, K. An expanding role for RAS GTPase activating proteins (RAS GAPs) in cancer. Adv Biol Regul 55, 1–14 (2014).

54. Song, M.S., Salmena, L. & Pandolfi, P.P. The functions and regulation of the PTEN tumour suppressor. Nat Rev Mol Cell Biol 13, 283–296 (2012).

55. Hayashi, T. et al. RASA1 and NF1 are Preferentially Co-Mutated and Define A Distinct Genetic Subset of Smoking-Associated Non-Small Cell Lung Carcinomas Sensitive to MEK Inhibition. Clin Cancer Res 24, 1436–1447 (2018).

56. Kitajima, S. & Barbie, D.A. RASA1/NF1-Mutant Lung Cancer: Racing to the Clinic? Clin Cancer Res 24, 1243–1245 (2018).

57. Nichols, R.J. et al. RAS nucleotide cycling underlies the SHP2 phosphatase dependence of mutant BRAF-, NF1- and RAS-driven cancers. Nat Cell Biol 20, 1064–1073 (2018).

58. Middleton, G. et al. The National Lung Matrix Trial of personalized therapy in lung cancer. Nature 583, 807–812 (2020).

59. Davies, B.R. et al. Preclinical pharmacology of AZD5363, an inhibitor of AKT: pharmacodynamics, antitumor activity, and correlation of monotherapy activity with genetic background. Mol Cancer Ther 11, 873–887 (2012).

60. O’Neill, A.C., Jagannathan, J.P. & Ramaiya, N.H. Evolving Cancer Classification in the Era of Personalized Medicine: A Primer for Radiologists. Korean J Radiol 18, 6–17 (2017).

61. Hanahan, D. & Weinberg, R.A. Hallmarks of cancer: the next generation. Cell 144, 646–674 (2011).

62. Zhao, Z. et al. Cooperative loss of RAS feedback regulation drives myeloid leukemogenesis. Nat Genet 47, 539–543 (2015).

63. Lock, R. & Cichowski, K. Loss of negative regulators amplifies RAS signaling. Nat Genet 47, 426–427 (2015).

64. Lawrence, M.S. et al. Mutational heterogeneity in cancer and the search for new cancer- associated genes. Nature 499, 214–218 (2013).

## References

1. Gao, Q. et al. Driver Fusions and Their Implications in the Development and Treatment of Human Cancers. Cell Rep 23, 227–238 e223 (2018).

2. Lu, X. et al. MET Exon 14 Mutation Encodes an Actionable Therapeutic Target in Lung Adenocarcinoma. Cancer Res 77, 4498–4505 (2017).

3. Liu, J. et al. An Integrated TCGA Pan-Cancer Clinical Data Resource to Drive High-Quality Survival Outcome Analytics. Cell 173, 400–416 e411 (2018).

4. Hallin, J. et al. The KRAS(G12C) Inhibitor MRTX849 Provides Insight toward Therapeutic Susceptibility of KRAS-Mutant Cancers in Mouse Models and Patients. Cancer Discov 10, 54–71 (2020).

5. Canon, J. et al. The clinical KRAS(G12C) inhibitor AMG 510 drives anti-tumour immunity. Nature 575, 217–223 (2019).

6. Jackson, E.L. et al. Analysis of lung tumor initiation and progression using conditional expression of oncogenic K-ras. Genes Dev 15, 3243–3248 (2001).

7. Winters, I.P. et al. Multiplexed in vivo homology-directed repair and tumor barcoding enables parallel quantification of Kras variant oncogenicity. Nat Commun 8, 2053 (2017).

8. Guerra, C. et al. Tumor induction by an endogenous K-ras oncogene is highly dependent on cellular context. Cancer Cell 4, 111–120 (2003).

9. Hingorani, S.R. et al. Preinvasive and invasive ductal pancreatic cancer and its early detection in the mouse. Cancer Cell 4, 437–450 (2003).

10. Zafra, M.P. et al. An In Vivo Kras Allelic Series Reveals Distinct Phenotypes of Common Oncogenic Variants. Cancer Discov 10, 1654–1671 (2020).

11. Paez, J.G. et al. EGFR mutations in lung cancer: correlation with clinical response to gefitinib therapy. Science 304, 1497–1500 (2004).

12. Lynch, T.J. et al. Activating mutations in the epidermal growth factor receptor underlying responsiveness of non-small-cell lung cancer to gefitinib. N Engl J Med 350, 2129–2139 (2004).

13. Politi, K. et al. Lung adenocarcinomas induced in mice by mutant EGF receptors found in human lung cancers respond to a tyrosine kinase inhibitor or to down- regulation of the receptors. Genes Dev 20, 1496–1510 (2006).

14. Li, D. et al. Bronchial and peripheral murine lung carcinomas induced by T790M- L858R mutant EGFR respond to HKI-272 and rapamycin combination therapy. Cancer Cell 12, 81–93 (2007).

15. Ji, H. et al. The impact of human EGFR kinase domain mutations on lung tumorigenesis and in vivo sensitivity to EGFR-targeted therapies. Cancer Cell 9, 485–495 (2006).

16. Zhu, H. et al. Oncogenic EGFR signaling cooperates with loss of tumor suppressor gene functions in gliomagenesis. Proc Natl Acad Sci U S A 106, 2712–2716 (2009).

17. Lin, Q. et al. The association between BRAF mutation class and clinical features in BRAF-mutant Chinese non-small cell lung cancer patients. J Transl Med 17, 298 (2019).

18. van Veen, J.E. et al. Mutationally-activated PI3’-kinase-alpha promotes de- differentiation of lung tumors initiated by the BRAF(V600E) oncoprotein kinase. Elife 8 (2019).

19. Dankort, D. et al. A new mouse model to explore the initiation, progression, and therapy of BRAFV600E-induced lung tumors. Genes Dev 21, 379–384 (2007).

20. Dankort, D. et al. Braf(V600E) cooperates with Pten loss to induce metastatic melanoma. Nat Genet 41, 544–552 (2009).

21. Charles, R.P., Iezza, G., Amendola, E., Dankort, D. & McMahon, M. Mutationally activated BRAF(V600E) elicits papillary thyroid cancer in the adult mouse. Cancer Res 71, 3863–3871 (2011).

22. Davies, H. et al. Mutations of the BRAF gene in human cancer. Nature 417, 949–954 (2002).

23. Chin, L. et al. Essential role for oncogenic Ras in tumour maintenance. Nature 400, 468–472 (1999).

24. Seeburg, P.H., Colby, W.W., Capon, D.J., Goeddel, D.V. & Levinson, A.D. Biological properties of human c-Ha-ras1 genes mutated at codon 12. Nature 312, 71–75 (1984).

25. Der, C.J., Finkel, T. & Cooper, G.M. Biological and biochemical properties of human rasH genes mutated at codon 61. Cell 44, 167–176 (1986).

26. Kwong, L.N. et al. Oncogenic NRAS signaling differentially regulates survival and proliferation in melanoma. Nat Med 18, 1503–1510 (2012).

27. Ackermann, J. et al. Metastasizing melanoma formation caused by expression of activated N-RasQ61K on an INK4a-deficient background. Cancer Res 65, 4005–4011 (2005).

28. Paik, P.K. et al. Response to MET inhibitors in patients with stage IV lung adenocarcinomas harboring MET mutations causing exon 14 skipping. Cancer Discov 5, 842–849 (2015).

29. Drilon, A. et al. Antitumor activity of crizotinib in lung cancers harboring a MET exon 14 alteration. Nat Med 26, 47–51 (2020).

30. Kim, M. et al. Patient-derived lung cancer organoids as in vitro cancer models for therapeutic screening. Nat Commun 10, 3991 (2019).

31. Gow, C.H. et al. Oncogenic Function of a KIF5B-MET Fusion Variant in Non- Small Cell Lung Cancer. Neoplasia 20, 838–847 (2018).

32. Gao, Y. et al. Allele-Specific Mechanisms of Activation of MEK1 Mutants Determine Their Properties. Cancer Discov 8, 648–661 (2018).

33. Cai, D., Choi, P.S., Gelbard, M. & Meyerson, M. Identification and Characterization of Oncogenic SOS1 Mutations in Lung Adenocarcinoma. Mol Cancer Res 17, 1002–1012 (2019).

34. Shaw, A.T. et al. Crizotinib versus chemotherapy in advanced ALK-positive lung cancer. N Engl J Med 368, 2385–2394 (2013).

35. Maddalo, D. et al. In vivo engineering of oncogenic chromosomal rearrangements with the CRISPR/Cas9 system. Nature 516, 423–427 (2014).

36. Pyo, K.H. et al. Establishment of a Conditional Transgenic Mouse Model Recapitulating EML4-ALK-Positive Human Non-Small Cell Lung Cancer. J Thorac Oncol 12, 491–500 (2017).

37. Chiarle, R. et al. NPM-ALK transgenic mice spontaneously develop T-cell lymphomas and plasma cell tumors. Blood 101, 1919–1927 (2003).

38. Soda, M. et al. Identification of the transforming EML4-ALK fusion gene in non- small-cell lung cancer. Nature 448, 561–566 (2007).

39. Subbiah, V. et al. Precision Targeted Therapy with BLU-667 for RET-Driven Cancers. Cancer Discov 8, 836–849 (2018).

40. Huang, Q. et al. Preclinical Modeling of KIF5B-RET Fusion Lung Adenocarcinoma. Mol Cancer Ther 15, 2521–2529 (2016).

41. Saito, M. et al. A mouse model of KIF5B-RET fusion-dependent lung tumorigenesis. Carcinogenesis 35, 2452–2456 (2014).

42. Shaw, A.T. et al. Crizotinib in ROS1-rearranged non-small-cell lung cancer. N Engl J Med 371, 1963–1971 (2014).

43. Arai, Y. et al. Mouse model for ROS1-rearranged lung cancer. PLoS One 8, e56010 (2013).

44. Inoue, M. et al. Mouse models for ROS1-fusion-positive lung cancers and their application to the analysis of multikinase inhibitor efficiency. Carcinogenesis 37, 452–460 (2016).

45. Farago, A.F. et al. Durable Clinical Response to Entrectinib in NTRK1- Rearranged Non-Small Cell Lung Cancer. J Thorac Oncol 10, 1670–1674 (2015).

46. Vaishnavi, A. et al. Oncogenic and drug-sensitive NTRK1 rearrangements in lung cancer. Nat Med 19, 1469–1472 (2013).

47. Drilon, A. et al. Response to ERBB3-Directed Targeted Therapy in NRG1- Rearranged Cancers. Cancer Discov 8, 686–695 (2018).

48. Hyman, D.M. et al. AKT Inhibition in Solid Tumors With AKT1 Mutations. J Clin Oncol 35, 2251–2259 (2017).

49. De Marco, C. et al. Mutant AKT1-E17K is oncogenic in lung epithelial cells. Oncotarget 6, 39634–39650 (2015).

50. Berger, A.H. et al. Oncogenic RIT1 mutations in lung adenocarcinoma. Oncogene 33, 4418–4423 (2014).

51. Pillai, R.N. et al. HER2 mutations in lung adenocarcinomas: A report from the Lung Cancer Mutation Consortium. Cancer 123, 4099–4105 (2017).

52. Perera, S.A. et al. HER2YVMA drives rapid development of adenosquamous lung tumors in mice that are sensitive to BIBW2992 and rapamycin combination therapy. Proc Natl Acad Sci U S A 106, 474–479 (2009).

53. Liu, S. et al. Targeting HER2 Aberrations in Non-Small Cell Lung Cancer with Osimertinib. Clin Cancer Res 24, 2594–2604 (2018).

54. Weinstein, E.J., Kitsberg, D.I. & Leder, P. A mouse model for breast cancer induced by amplification and overexpression of the neu promoter and transgene. Mol Med 6, 4–16 (2000).

55. Guy, C.T. et al. Expression of the neu protooncogene in the mammary epithelium of transgenic mice induces metastatic disease. Proc Natl Acad Sci U S A 89, 10578–10582 (1992).

56. Greulich, H. et al. Functional analysis of receptor tyrosine kinase mutations in lung cancer identifies oncogenic extracellular domain mutations of ERBB2. Proc Natl Acad Sci U S A 109, 14476–14481 (2012).

57. Engelman, J.A. et al. Effective use of PI3K and MEK inhibitors to treat mutant Kras G12D and PIK3CA H1047R murine lung cancers. Nat Med 14, 1351–1356 (2008).

58. Trejo, C.L. et al. Mutationally activated PIK3CA(H1047R) cooperates with BRAF(V600E) to promote lung cancer progression. Cancer Res 73, 6448–6461 (2013).

59. Devarakonda, S. et al. Genomic Profiling of Lung Adenocarcinoma in Never-Smokers. *J Clin Oncol*, JCO2101691 (2021).

60. Sanchez-Vega, F. et al. Oncogenic Signaling Pathways in The Cancer Genome Atlas. Cell 173, 321–337 e310 (2018).

61. Bailey, M.H. et al. Comprehensive Characterization of Cancer Driver Genes and Mutations. Cell 173, 371–385 e318 (2018).

62. Yi, K.H. & Lauring, J. Recurrent AKT mutations in human cancers: functional consequences and effects on drug sensitivity. Oncotarget 7, 4241–4251 (2016).

63. Dobashi, Y. et al. Diverse involvement of isoforms and gene aberrations of Akt in human lung carcinomas. Cancer Sci 106, 772–781 (2015).

64. Arboleda, M.J. et al. Overexpression of AKT2/protein kinase Bbeta leads to up- regulation of beta1 integrins, increased invasion, and metastasis of human breast and ovarian cancer cells. Cancer Res 63, 196–206 (2003).

65. Zhang, Y. et al. A Pan-Cancer Proteogenomic Atlas of PI3K/AKT/mTOR Pathway Alterations. Cancer Cell 31, 820–832 e823 (2017).

66. Chaft, J.E. et al. Coexistence of PIK3CA and other oncogene mutations in lung adenocarcinoma-rationale for comprehensive mutation profiling. Mol Cancer Ther 11, 485–491 (2012).

67. Dbouk, H.A. et al. Characterization of a tumor-associated activating mutation of the p110beta PI 3-kinase. PLoS One 8, e63833 (2013).

68. Pazarentzos, E. et al. Oncogenic activation of the PI3-kinase p110beta isoform via the tumor-derived PIK3Cbeta(D1067V) kinase domain mutation. Oncogene 35, 1198–1205 (2016).

69. Whale, A.D., Colman, L., Lensun, L., Rogers, H.L. & Shuttleworth, S.J. Functional characterization of a novel somatic oncogenic mutation of PIK3CB. Signal Transduct Target Ther 2, 17063 (2017).

70. Liyasova, M.S., Ma, K. & Lipkowitz, S. Molecular pathways: cbl proteins in tumorigenesis and antitumor immunity-opportunities for cancer treatment. Clin Cancer Res 21, 1789–1794 (2015).

71. Maertens, O. & Cichowski, K. An expanding role for RAS GTPase activating proteins (RAS GAPs) in cancer. Adv Biol Regul 55, 1–14 (2014).

72. Ahmad, M.K., Abdollah, N.A., Shafie, N.H., Yusof, N.M. & Razak, S.R.A. Dual-specificity phosphatase 6 (DUSP6): a review of its molecular characteristics and clinical relevance in cancer. Cancer Biol Med 15, 14–28 (2018).

73. Owens, D.M. & Keyse, S.M. Differential regulation of MAP kinase signalling by dual-specificity protein phosphatases. Oncogene 26, 3203–3213 (2007).

74. Sato, K. et al. Fusion Kinases Identified by Genomic Analyses of Sporadic Microsatellite Instability-High Colorectal Cancers. Clin Cancer Res 25, 378–389 (2019).

75. Nonami, A. et al. Spred-1 negatively regulates interleukin-3-mediated ERK/mitogen-activated protein (MAP) kinase activation in hematopoietic cells. J Biol Chem 279, 52543–52551 (2004).

76. McClatchey, A.I. & Cichowski, K. SPRED proteins provide a NF-ty link to Ras suppression. Genes Dev 26, 1515–1519 (2012).

77. Hanafusa, H., Torii, S., Yasunaga, T. & Nishida, E. Sprouty1 and Sprouty2 provide a control mechanism for the Ras/MAPK signalling pathway. Nat Cell Biol 4, 850–858 (2002).

78. Yusoff, P. et al. Sprouty2 inhibits the Ras/MAP kinase pathway by inhibiting the activation of Raf. J Biol Chem 277, 3195–3201 (2002).

79. Kim, H.J. & Bar-Sagi, D. Modulation of signalling by Sprouty: a developing story. Nat Rev Mol Cell Biol 5, 441–450 (2004).

80. Chakravarty, D., et al. OncoKB: A Precision Oncology Knowledge Base. JCO Precis Oncol 2017 (2017).

81. Liu, C. et al. Mosaic analysis with double markers reveals tumor cell of origin in glioma. Cell 146, 209–221 (2011).

82. Zhu, Y. et al. Ablation of NF1 function in neurons induces abnormal development of cerebral cortex and reactive gliosis in the brain. Genes Dev 15, 859–876 (2001).

83. Madisen, L. et al. A robust and high-throughput Cre reporting and characterization system for the whole mouse brain. Nat Neurosci 13, 133–140 (2010).

84. Chiou, S.H. et al. Pancreatic cancer modeling using retrograde viral vector delivery and in vivo CRISPR/Cas9-mediated somatic genome editing. Genes Dev 29, 1576–1585 (2015).

85. Okawa, H. et al. Hepatocyte-specific deletion of the keap1 gene activates Nrf2 and confers potent resistance against acute drug toxicity. Biochem Biophys Res Commun 339, 79–88 (2006).

86. Bardeesy, N. et al. Loss of the Lkb1 tumour suppressor provokes intestinal polyposis but resistance to transformation. Nature 419, 162–167 (2002).

87. Jonkers, J. et al. Synergistic tumor suppressor activity of BRCA2 and p53 in a conditional mouse model for breast cancer. Nat Genet 29, 418–425 (2001).

88. Rogers, Z.N. et al. A quantitative and multiplexed approach to uncover the fitness landscape of tumor suppression in vivo. Nat Methods 14, 737–742 (2017).

89. Rogers, Z.N. et al. Mapping the in vivo fitness landscape of lung adenocarcinoma tumor suppression in mice. Nat Genet 50, 483–486 (2018).

90. Cai, H., et al. A functional taxonomy of tumor suppression in oncogenic KRAS-driven lung cancer. Under consideration (2021).

91. Murray, C.W. et al. An LKB1-SIK Axis Suppresses Lung Tumor Growth and Controls Differentiation. Cancer Discov 9, 1590–1605 (2019).

92. Orr, H.A. Fitness and its role in evolutionary genetics. Nat Rev Genet 10, 531–539 (2009).

93. J.B.S., H. The mathematical theory of natural and artificial selection. Mathematical Proceedings of the Cambridge Philosophical Society 23, 607–615 (1927).

94. Geraldine A. Van der Auwera, B.D.O.C. Genomics in the Cloud. (2020).

95. McLaren, W. et al. The Ensembl Variant Effect Predictor. Genome Biol 17, 122 (2016).

96. Fedchenko, N. & Reifenrath, J. Different approaches for interpretation and reporting of immunohistochemistry analysis results in the bone tissue - a review. Diagn Pathol 9, 221 (2014).

97. Feoktistova, M., Geserick, P. & Leverkus, M. Crystal Violet Assay for Determining Viability of Cultured Cells. Cold Spring Harb Protoc 2016, pdb prot087379 (2016).

98. Nichols, R.J. et al. RAS nucleotide cycling underlies the SHP2 phosphatase dependence of mutant BRAF-, NF1- and RAS-driven cancers. Nat Cell Biol 20, 1064–1073 (2018).

99. Ianevski, A., Giri, A.K. & Aittokallio, T. SynergyFinder 2.0: visual analytics of multi-drug combination synergies. Nucleic Acids Res 48, W488–W493 (2020).

100. Loewe, S. The problem of synergism and antagonism of combined drugs. Arzneimittelforschung 3, 285–290 (1953).

101. Dobin, A. et al. STAR: ultrafast universal RNA-seq aligner. Bioinformatics 29, 15–21 (2013).

102. Li, B. & Dewey, C.N. RSEM: accurate transcript quantification from RNA-Seq data with or without a reference genome. BMC Bioinformatics 12, 323 (2011).

103. Soneson, C., Love, M.I. & Robinson, M.D. Differential analyses for RNA-seq: transcript-level estimates improve gene-level inferences. F1000Res 4, 1521 (2015).

104. Love, M.I., Huber, W. & Anders, S. Moderated estimation of fold change and dispersion for RNA-seq data with DESeq2. Genome Biol 15, 550 (2014).

105. Subramanian, A. et al. Gene set enrichment analysis: a knowledge-based approach for interpreting genome-wide expression profiles. Proc Natl Acad Sci U S A 102, 15545–15550 (2005).

106. Hanzelmann, S., Castelo, R. & Guinney, J. GSVA: gene set variation analysis for microarray and RNA-seq data. BMC Bioinformatics 14, 7 (2013).

